# High-throughput characterization of photocrosslinker-bearing ion channel variants to map residues critical for function and pharmacology

**DOI:** 10.1101/2020.11.24.392498

**Authors:** Nina Braun, Søren Friis, Christian Ihling, Andrea Sinz, Jacob Andersen, Stephan A. Pless

## Abstract

Incorporation of non-canonical amino acids (ncAAs) can endow proteins with novel functionalities, such as crosslinking or fluorescence. In ion channels, the function of these variants can be studied with great precision using standard electrophysiology, but this approach is typically labor intensive and low throughput. Here, we establish a high-throughput protocol to conduct functional and pharmacological investigations of ncAA-containing hASIC1a (human acid-sensing ion channel 1a) variants in transiently transfected mammalian cells. We introduce three different photocrosslinking ncAAs into 103 positions and assess the function of the resulting 309 variants with automated patch-clamp (APC). We demonstrate that the approach is efficient and versatile, as it is amenable to assessing even complex pharmacological modulation by peptides. The data show that the acidic pocket is a major determinant for current decay and live-cell crosslinking provides insight into the hASIC1a-psalmotoxin-1 interaction. Further, we provide evidence that the protocol can be applied to other ion channels, such as P2X2 and GluA2 receptors. We therefore anticipate the approach to enable future APC-based studies of ncAA-containing ion channels in mammalian cells.

## Introduction

Genetic code expansion approaches allow the incorporation of non-canonical amino acids (ncAAs) with unique chemical properties into proteins. Over the past two decades, this method has greatly facilitated protein modification and functionalization beyond the confines of the genetic code [1]. Ion channels have proven highly suited to ncAA incorporation, as evidenced by the success in introducing photocrosslinking, photoswitchable or fluorescent ncAAs into numerous members of this large and diverse protein family [2–4]. Among the ncAA subclasses, photocrosslinkers have proven particularly versatile, as they allow for the trapping of ion channels in certain conformational states [5–8], capturing of protein-protein interactions [9–12] and covalent linking of receptor-ligand complexes to delineate ligand binding sites [13–17].

Typically, incorporation of ncAAs is achieved by repurposing a stop codon to encode for a ncAA supplied by an orthogonal tRNA/aminoacyl tRNA synthetase (aaRS) pair. But the incorporation efficiency can be variable and unspecific incorporation of naturally occurring amino acids can result in inhomogeneous protein populations [2]. Verification of site-specific ncAA incorporation can therefore be laborious and time-consuming, especially in combination with detailed functional characterization. As a result, most studies have focused on only a limited number of incorporation sites, and the evaluation of potential functional or pharmacological effects of ncAA incorporation often remained minimal. In principle, automated patch-clamp (APC) devices offer fast and efficient high-throughput testing and have recently gained increasing popularity for electrophysiological interrogation of a diverse set of ion channels [18–22]. However, a combination of low efficiency of transient transfection in mammalian cells and limited ncAA incorporation rates have thus far prevented functional screening of ncAA-containing ion channel variants on APC platforms.

Here, we sought to overcome these limitations by developing a fluorescence-activated cell sorting (FACS)-based approach to enrich the population of transiently transfected cells expressing ncAA-containing ion channels. Using the human acid-sensing ion channel 1a (hASIC1a) as an example, we incorporated three different ncAA photocrosslinkers (AzF (4-Azido-L-phenylalanine), Bpa (4-Benzoyl-L-phenylalanine) and Se-AbK ((R)-2-Amino-3-{2-[2-(3-methyl-3H-diazirin-3-yl)-ethoxycarbonylamino]-ethylselanyl}-propionic acid)) at 103 positions throughout its intracellular, extracellular and transmembrane domains.

ASICs are trimeric ligand-gated ion channels that open a weakly sodium-selective pore in response to proton binding to the so-called acidic pocket and likely other sites in the extracellular domain [23]. Apart from contributions to synaptic plasticity [24, 25], ASICs have recently gained increasing attention as potential drug targets for pain and stroke [26–35]. The six different human ASIC isoforms (ASIC1a, 1b, 2a, 2b, 3 and 4) are modulated by an impressive array of neuropeptides and venom-derived toxins that bind to the large extracellular domain [24, 36, 37]. Intriguingly, the extent and type of modulation (e.g. inhibition vs potentiation) are often highly dependent on ambient proton concentration, as well as subtype and species origin [38, 39]. This poses challenges for pharmacological profiling and motivates a detailed understanding of the mechanism and site of action of these peptides, to eventually generate lead compounds that could potentially target pain or stroke.

In this study, we establish a protocol to functionally screen ncAA-containing ion channels in transiently transfected cells on an APC platform. The 384-well setup of the SyncroPatch 384PE (Nanion Technologies) allows the efficient characterization of 309 hASIC1a variants and we show that ncAA incorporation is tolerated in over 50% of the positions. Incorporation of bulky ncAA photocrosslinkers generally results in lower pH sensitivity, especially around the acidic pocket, where ncAA incorporation also greatly accelerates current decay kinetics. We further demonstrate differential channel modulation by the neuropeptide big dynorphin (BigDyn; [40]) and by psalmotoxin-1 (PcTx1; [41]), a toxin derived from tarantula venom. Lastly, we turn to UV-induced photocrosslinking to covalently trap channel-toxin complexes and thus map the hASIC1a-PcTx1 interaction in live cells. Overall, our work highlights that ncAA-containing ion channels, including ASICs, ionotropic glutamate and P2X receptors, are amenable to APC-based high-throughput screening. We further demonstrate how this approach, when used with ncAA photocrosslinkers, can be harnessed to investigate protein-peptide or protein-protein interactions *in cellulo*.

## Results

### Development of an APC screen to validate ncAA incorporation into hASIC1a

In order to efficiently assess functional incorporation of ncAAs into human ASIC1a (hASIC1a), we developed an APC screen to record proton-gated channel activation (Figure 1). To this end, we co-transfected 103 different hASIC1a variants containing individual TAG stop codons throughout the protein together with the suppressor tRNA/ncAA-RS pair for either AzF, Bpa or Se-AbK and a GFP-reporter carrying a TAG at Y40 (for Bpa and Se-AbK) or Y151 (for AzF, as we observed a higher degree of unspecific incorporation in the Y40TAG variant with AzF) into custom-made ASIC1a-KO HEK 293T cells [17, 42–44]. The corresponding ncAA was supplied in the cell culture medium six hours after transfection or omitted from the experiment in incorporation control samples. To increase cell viability and uptake efficiency, we synthesized the methylester derivates of AzF and Bpa [8, 45]. This allowed us to supplement the cell media with 50- and 100-fold lower ncAA concentration compared to previous studies, respectively [7, 13].

**Figure 1:**
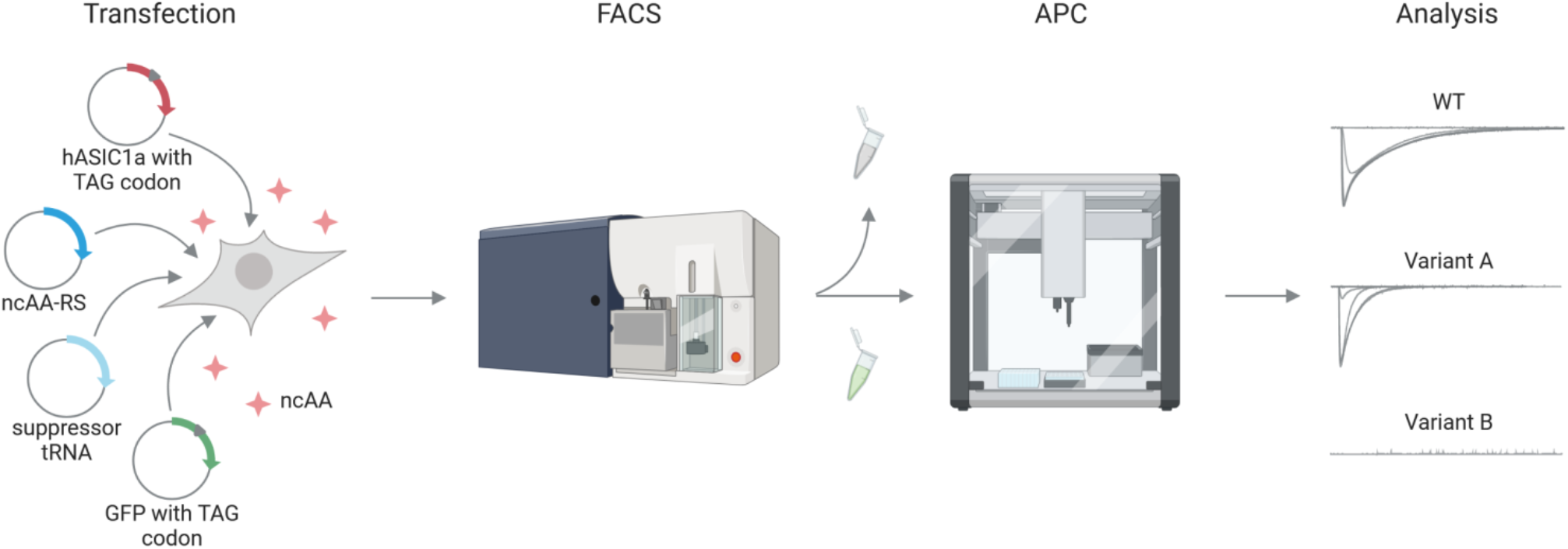
Schematic illustration of the workflow to assess ncAA incorporation into hASIC1a. HEK 293T ASIC1a-KO cells are transfected with hASIC1a containing a TAG stop codon at the site of interest, a co-evolved suppressor tRNA/ncAA-RS pair and a TAG-containing GFP reporter. ncAA is supplied in the cell culture medium. 48 hours after transfection, cells are sorted for green fluorescence on a FACS BD Aria I and those showing fluorescence are subjected to APC on a SyncroPatch 384PE to measure proton concentration response curves.

After 48 hours, cells grown in the presence of ncAA were sorted for green fluorescence to enrich the population of transfected cells, which were then submitted to APC to record proton-gated currents. Using GFP fluorescence as a proxy, we determined a transfection efficiency of 62.9 ± 9.5% for hASIC1a WT and an average of 11.2 ± 5% for the ncAA variants (Figure S1, Table S1). Without the FACS step, the latter rate would translate into less than 10% of the APC wells being occupied by transfected cells, precluding efficient APC experiments. By contrast, the cell sorting improves occupation to around 46% of wells with successful patch also displaying proton-gated currents (62% for AzF, 29% for Bpa and 48% for Se-AbK) and is therefore an indispensable element for the use of transiently transfected cells in APC (Figure S1).

The 384-well system of the SyncroPatch 384PE allows for parallel concentration response curve measurements on 24 different samples, enabling us to test 11 different channel variants with corresponding incorporation controls (cells grown in the absence of ncAA), as well as hASIC1a WT and untransfected cells in less than one hour, with up to 16 replicates per sample. Specifically, we embarked to functionally interrogate 103 positions throughout the hASIC1a sequence: 38 positions in the N-terminal domain (Figure S2), 24 positions in the transmembrane domain and interface region (Figure S3), 29 positions in the C-terminal domain (Figure S4) and 12 positions around the acidic pocket (Figure 3 and S8). The current traces in Figure 2A show typical pH-induced inward currents of hASIC1a WT with a pH_50_ of 6.64 ± 0.12 (n=182), in line with previous studies [46, 47], as well as a variant with lower proton sensitivity containing AzF in the acidic pocket (T236AzF, pH_50_ 6.17 ± 0.14, n=10). Interestingly, the incorporation of Bpa, AzF and Se-AbK at position W46 did not result in proton-gated currents (Figure 2A, Figure S3), despite a previous report showing functional incorporation of a bulky ncAA at this conserved Trp in the M1 helix [48]. We analysed all variants for mean peak current size and pH_50_ to compare incorporation efficiency and proton sensitivity, respectively (Figures S2-4 and S8, Table S1). Furthermore, we routinely assessed the extent of tachyphylaxis [49] and variants displaying >20% current decrease after reaching the peak current are indicated in Figures 3 and S2-8 as well as Table S1.

**Figure 2:**
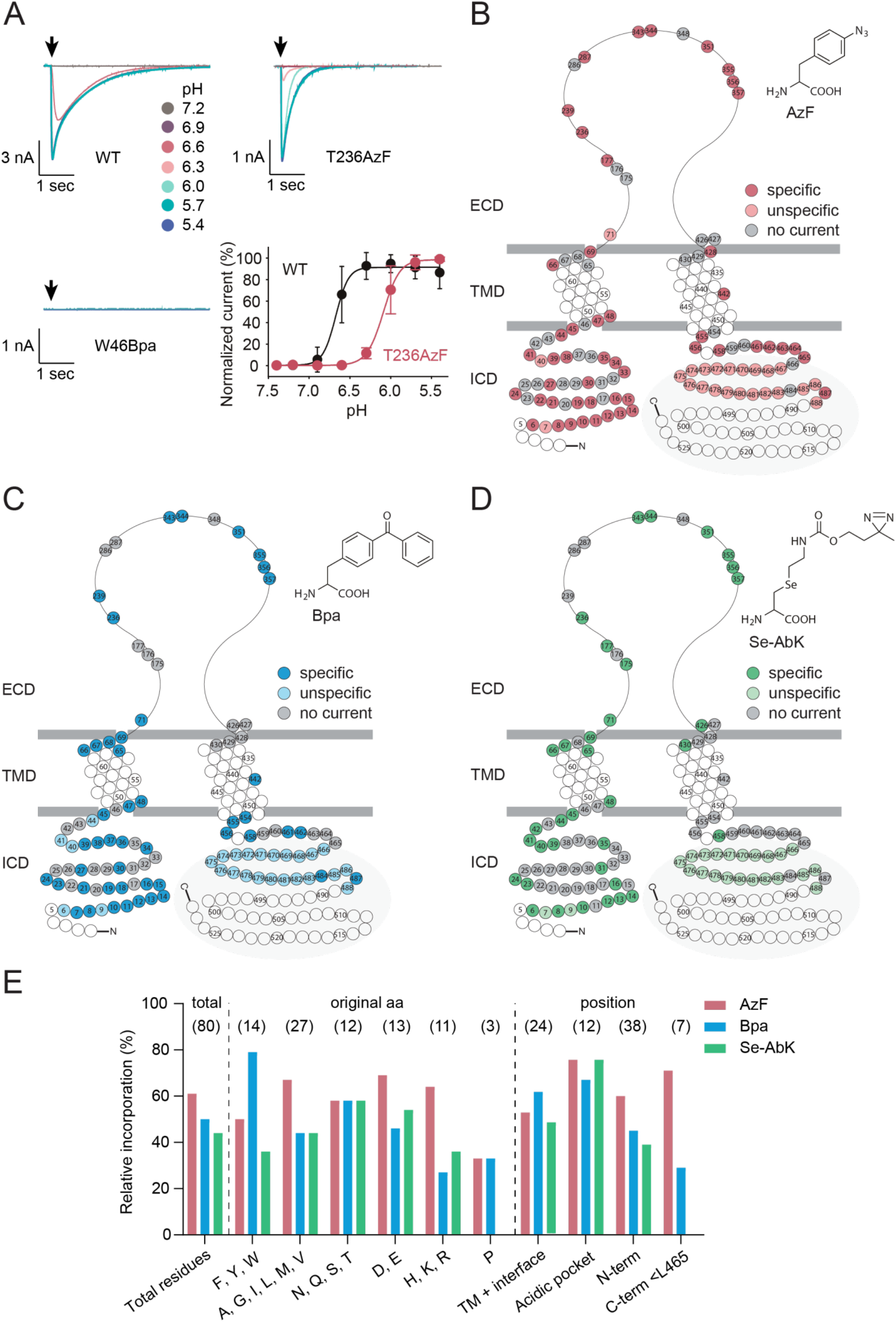
Incorporation of ncAA crosslinkers is tolerated in all domains of hASIC1a and produces functional channel variants. (A) Representative current traces for pH response curves of hASIC1a WT, T236AzF and W46Bpa recorded on the SyncroPatch 384PE, pH response curve in bottom right corner (WT pH_50_ 6.64 ± 0.12, n=182; T236AzF pH_50_ 6.17 ± 0.14, n=10). (B-D) Snake plot representations indicating specific, unspecific and unsuccessful incorporation (no current) for AzF (B), Bpa (C) and Se-AbK (D). Specific incorporation (circles with darker shade) is defined as pH-dependent peak currents >1 nA observed in cells grown in the presence, but not in the absence of ncAA, whereas unspecific incorporation (circles with lighter shade) indicates that currents were observed both in the presence and absence of ncAA. Positions indicated by grey circles did not yield functional channel variants when replaced by an ncAA (no current), while those coloured in white were not tested. The grey area highlights positions distal of L465. (E) Relative incorporation rates of AzF (red), Bpa (blue) and Se-AbK (green) at 80 different hASIC1a positions. Exchanged amino acids are grouped for original side chain properties and position within the channel, respectively (TM: transmembrane helices; N-term: N-terminus; C-term <L465: C-terminus up to and including L465). Relative incorporation rates were calculated by dividing the number of positions successfully replaced with a ncAA by the total number of positions at which incorporation was attempted. Positions distal of L465 were excluded from the analysis (highlighted in grey in B-D), as more distal deletions result in truncated, but functional channels (see Figures S4-6).

**Figure 3:**
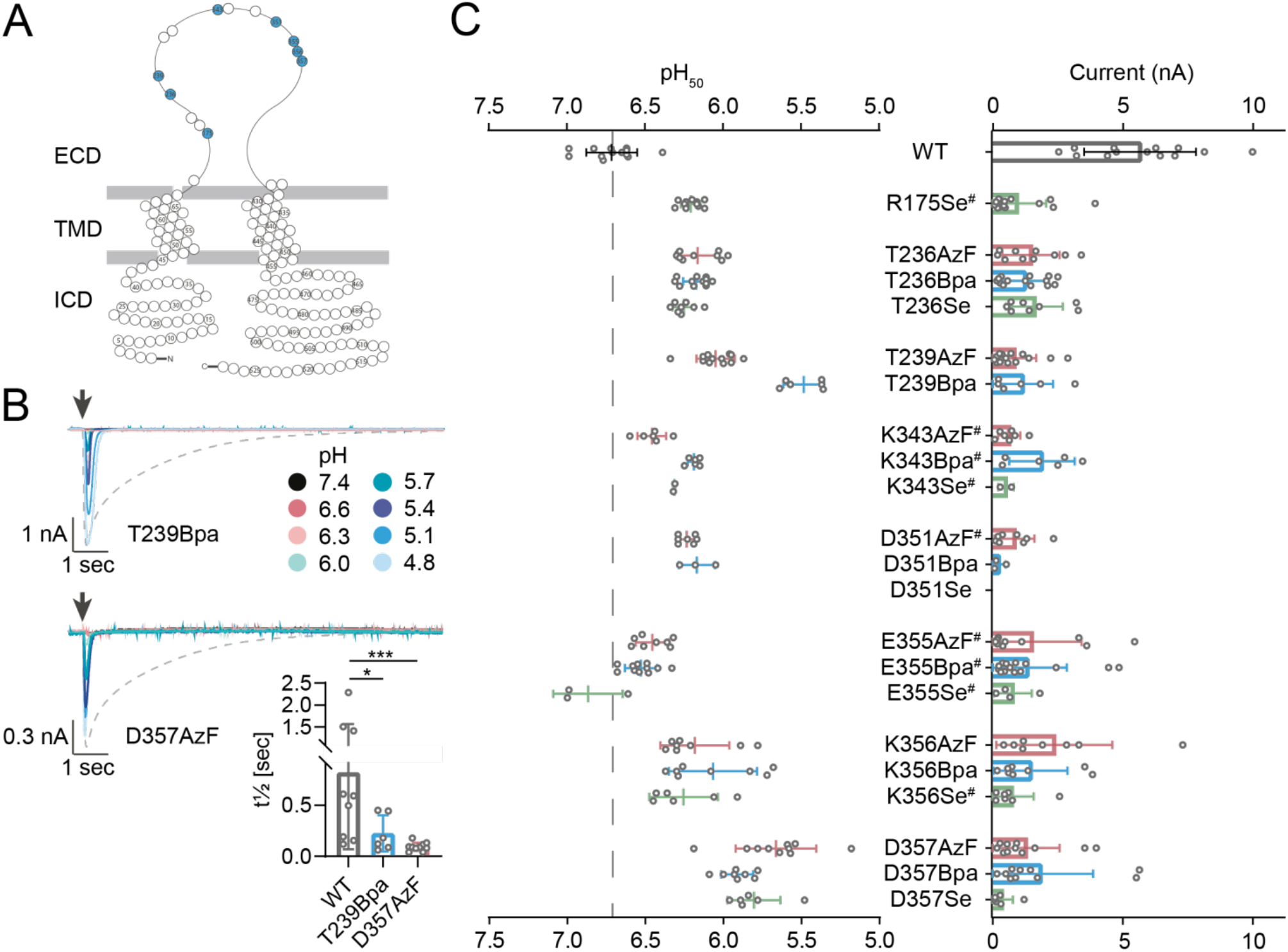
Incorporation of ncAA photocrosslinkers into the acidic pocket results in channel variants with lowered proton sensitivity and accelerated current decay. (A) Snake plot of hASIC1a with the assessed positions highlighted in blue. (B) Representative current traces of variants T239Bpa and D357AzF as recorded on the SyncroPatch 384PE, with arrows indicating the time of proton application. Dashed lines indicate WT current in response to pH 6.0 application. Bar graph shows mean t½ ± S.D. of current decay. (C) Incorporation of AzF (red), Bpa (blue) and Se-AbK (green) at 8 positions around the acidic pocket results in lowered proton sensitivity for several variants. Dot plots comparing pH_50_ (left) and peak current sizes (right), bars indicate mean ± S.D., (^#^) indicates >20% tachyphylaxis (see also Figure S8 and Table S1), (*) denotes significant difference between t½ of current decay, p < 0.05; (***): p < 0.001; Mann-Whitney test (see also Table S3).

To provide a comprehensive overview, we mapped incorporation patterns for the three photocrosslinkers onto snake plots schematically depicting an ASIC1a subunit (Figure 2B-D). We defined specific incorporation (circles with dark colour shade) as proton-gated currents of >1 nA observed in the presence of ncAA, and minimal (<500 pA) proton-gated currents in the absence of ncAA. If currents >500 pA were observed in the absence of ncAA, incorporation was considered unspecific (circles with lighter colour shade), while positions labelled in grey did not yield substantial currents in either condition (<1 nA). However, we cannot exclude the possibility of underestimating the degree of unspecific incorporation, as enriching transfected cells grown in the absence of ncAA by FACS was not feasible due to the low number of cells displaying GFP fluorescence (2.2 ± 1.7%). On the other hand, by defining incorporation as not successful for currents <1 nA, we are aware that we may have potentially excluded variants in which specific ncAA incorporation resulted in reduced open probability or lower conductance.

As is apparent from the snake plots, we observed robust incorporation in the N-terminus, around the acidic pocket and in the proximal C-terminus. Indeed, among the 80 positions tested up to and including L465, AzF resulted in functional channel variants in 61% of cases, compared to 50% for Bpa and 44% for Se-AbK (Figure 2E).

By contrast, all three crosslinkers showed mostly unspecific incorporation distal of L465, with WT-like current phenotypes from position 467 onwards (Figure S4 and S5A-C). This led us to hypothesize that channel constructs truncated in this region are functional. To investigate this further, we inserted an additional TGA stop codon for several variants, confirmed channel truncation by comparing molecular weight on a Western blot and measured concentration response curves in APC and two electrode voltage-clamp (TEVC) (Figure S5D-E). We found that channels truncated after H463 or K464 yielded no current in either APC or TEVC, but truncation after L465 produced a variant with strong tachyphylaxis in HEK 293T cells (Figure S5D) and truncation after C466 or R467 resulted in channels with WT-like proton sensitivity in both APC and TEVC. We conclude that the C-terminus distal of position 465 is not essential for proton-gated channel activity and that it is not possible to differentiate between currents originating from truncated and full-length protein to evaluate ncAA incorporation. We therefore added a C-terminal 1D4-tag to the hASIC1a construct to selectively purify full-length protein and compare the amounts in cells grown in the presence or absence of ncAA. This strategy confirms efficient incorporation in the distal C-terminus (Figure S6A). Additionally, liquid chromatography/tandem mass spectrometry data revealed that Bpa can be specifically incorporated at positions distal of L465 (A480, Figure S6B).

For the 80 positions up to and including L465, we evaluated incorporation efficiency by comparing how many positions could be functionally replaced by each of the ncAA photocrosslinkers, based on the nature of the side chain occupying the position in the native channel and the position within the protein overall. We did not find evidence for pronounced global trends, but for instance Bpa incorporation was tolerated best at originally aromatic side chains (79%), while replacement of basic residues was least successful (27%) (Figure 2E). The three tested prolines could not be exchanged for any of the ncAAs. Interestingly, and in contrast to our expectations, Se-AbK incorporation only produced functional variants in 33% of cases when replacing structurally similar Lys and Arg side chains, while success rates were higher at polar and acidic side chains (58% and 54%, respectively). AzF incorporation rates were similar throughout all protein domains, whereas Bpa was better tolerated in the transmembrane regions and less in the N- and C-termini and Se-AbK incorporation in the M2 helix and C-terminus was negligible (Figure 2E). Overall, incorporating the three photocrosslinkers produced functional variants in all protein domains, albeit with varying success rates.

Together, we show that combining FACS with APC affords the time-efficient functional characterization of over 300 hASIC1a variants and provides a versatile platform to assess successful ncAA incorporation throughout all protein domains.

To evaluate if the established APC screen can also serve as a platform for other ion channels, we applied it to selected TAG variants of the rat P2X2 and rat GluA2 receptors. Specifically, we compared currents upon exposure to two different concentrations of ATP or glutamate, respectively (Figure S7 and Table S2, [5, 7, 50]). Incorporation of AzF into position K296 of the rP2X2 receptor is unspecific, whereas that of Bpa is efficient and specific (Figure S7A). For GluA2, incorporation patterns at Y533 and S729 are identical to those observed in previous studies using manual patch-clamp (Figure S7B, [5, 7]). Incorporation of AzF at Y533 is tolerated with currents of 1.21 ± 0.96 nA (n=30, compared to 600 ± 100 pA (n=15) reported by Poulsen *et al.*), while incorporation of Bpa does not produce a functional channel. At position S729, we observe small currents of 390 ± 330 pA for AzF (n=17) and 280 ± 240 pA for Bpa (n=16, compared to 470 ± 50 pA reported by Klippenstein *et al.* for S729Bpa). Importantly, as GluA2 gating is fast compared to the perfusion speed of the SyncroPatch 384PE and Klippenstein *et al.* report increased desensitization rates for S729 variants, we pre-incubated cells with 100 μM cyclothiazide to slow desensitization and increase the likelihood of resolving the GluA2 peak current [51]. While our GluA2 and P2X2 data show that target-specific optimization of the ligand-application protocols is required, they illustrate that our APC screening approach can be applied to a variety of different ion channels and yields results comparable to those obtained with conventional ncAA-incorporation protocols.

### Photocrosslinker incorporation in the acidic pocket decreases proton sensitivity and accelerates current decay

During the design of the construct library for the APC screen, we consulted the 2.8 Å resolution structure of PcTx1 bound to chicken ASIC1 (PDB 4FZ0) to select 12 positions around the acidic pocket that are in sufficiently close proximity to potentially form covalent crosslinks with PcTx1 if replaced by a ncAA [52] (Figure S8A). Most of the resulting ncAA channel variants were functional, but in several instances, the initially applied proton concentration range of up to pH 5.4 did not yield saturating currents (Figure S8B/C). Consequently, we re-evaluated these variants using a lower pH range to resolve the pH_50_ and re-assess peak current size (Figure 3). This allowed us to determine EC_50_ values for all variants and confirmed that hASIC1a variants containing ncAAs in the acidic pocket display markedly reduced proton sensitivity, with pH_50_ values as low as 5.49 ± 0.13 (T239Bpa, mean ± S.D., n=6) and 5.66 ± 0.26 (D357AzF, mean ± S.D., n=10). Additionally, we observed substantial changes in current shape compared to WT. For example, current decay rates were increased for T239Bpa (t½ 224 ± 176 ms, n=6) and D357AzF (t½ 93.8 ± 40.9 ms, n=10) compared to WT (t½ 818 ± 750 ms, n=9), indicating possible effects of the photocrosslinkers on channel gating (rates of desensitization or closure, Figure 3B and Table S3). Overall, we found that incorporation of Se-AbK was least efficient, so all subsequent experiments focused on AzF- and Bpa-containing channel variants.

As hASIC1a variants with ncAAs around the acidic pocket displayed markedly altered proton sensitivity and current decay rates, we next wanted to assess if these variants can still be modulated by two peptide gating modifiers that interact with the acidic pocket, BigDyn and PcTx1.

### Peptide modulation is retained in hASIC1a variants containing photocrosslinkers in the acidic pocket

First, we investigated the neuropeptide BigDyn, which interacts with the acidic pocket and shifts the proton dependence of both activation and SSD [17]. A key physiological function of BigDyn is to limit ASIC1a steady-state desensitization (SSD) [40]. In order to define the appropriate pH for BigDyn application on each variant, we first established an APC-based protocol to determine SSD curves. Due to the open-well system of the SyncroPatch 384PE, lowering the conditioning pH to assess SSD required multiple mixing steps, which we simulated on a pH meter to determine the apparent pH the cells are exposed to before each activation (Figure S9). Using this approach, we obtained a pH_50_ SSD of 6.91 ± 0.02 for hASIC1a WT (n=40), which is lower than the value reported in *Xenopus laevis* oocytes (pH_50_ SSD = 7.05 ± 0.01, Figure S10A+B). Notably, we also observed a more shallow Hill slope for WT compared to oocytes (n_H_ 3.16 ± 0.42 vs 9.45 ± 2.84), but not for any of the tested variants in the acidic pocket or interface region (Figure S10B-F, Table S4). SSD profiles of the ncAA-containing variants varied with pH_50_ SSD values ranging from 7.15 ± 0.01 (E177Bpa, n=12) to 6.76 ± 0.06 (K356AzF, n=7, Table S4), with most variants displaying a slightly increased proton sensitivity compared to WT. This is in contrast to the observed pattern of reduced proton sensitivity for proton-gated activation, suggesting that incorporation of ncAA photocrosslinkers in the acidic pocket modulates proton sensitivity of activation and SSD differentially. For our subsequent APC experiments to assess BigDyn modulation, we chose a conditioning pH that led to around 10% remaining current upon activation.

Here, we focused on AzF-containing variants for which we had previously detected crosslinking to BigDyn on Western blots to evaluate if the observed peptide-channel interaction also results in functional modulation [17]. Cells were exposed to SSD-inducing pH conditions in the presence or absence of 3 μM BigDyn and the resulting currents upon pH 5.6 activation were normalized to control currents after incubation at pH 7.6 (Figure 4A+B). Control cells not exposed to BigDyn exhibited SSD to 0-30% mean remaining current (Figure S11, Table S5), while BigDyn co-application during conditioning limited SSD to varying degrees (Figure 4B). BigDyn increased rescue from pH-induced SSD in all tested AzF-containing variants, but did not do so in WT, despite a similar trend (Figure 4B). For all tested variants, we regularly observed incomplete SSD after the first conditioning step, but this typically increased after the second conditioning step (see Figure S11). This could point towards possible confounding effects by the repeated solution mixing to achieve the desired conditioning pH described above. However, despite the reduced control over the conditioning pH compared to using a perfusion system with continuous flow, it was still possible to determine if BigDyn modulates hASIC1a SSD. In short, the APC setup enables rapid evaluation of several channel variants with different SSD profiles for BigDyn modulation in a single experiment.

**Figure 4:**
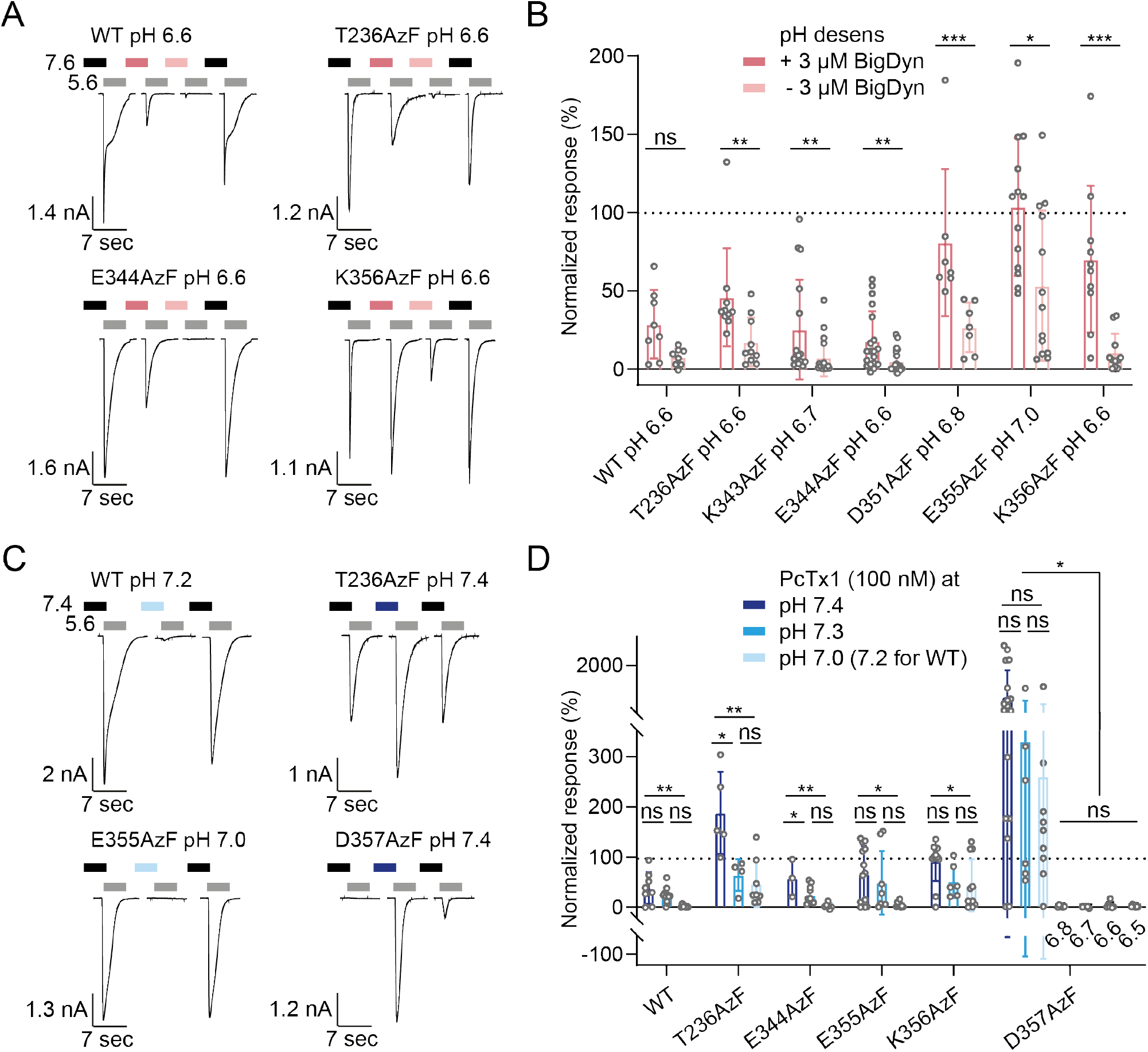
Peptide modulation of hASIC1a WT and selected variants containing AzF in the acidic pocket. (A) Characteristic current traces and (B) normalized response after SSD in absence or presence of BigDyn for hASIC1a WT and six ncAA variants. Cells were incubated at the desensitizing pH specified for each variant with or without 3 μM BigDyn for 2 min (pink bars) before activation at pH 5.6 (grey bars, 5 sec) and the currents were normalized to the average of two control currents after conditioning at pH 7.6 (black bars; control traces shown in Figure S11). (C) Exemplary current traces and (D) bar graph for PcTx1 modulation of hASIC1a WT and selected variants containing AzF in the acidic pocket at different pH. Cells were incubated with 100 nM PcTx1 at varying pH for 2 min (blue bars) before activation at pH 5.6 (grey bars, 5 sec) and the current was normalized to the average of the four preceding and following control currents after conditioning at pH 7.4 (black bars). Bar graphs show mean ± S.D, dashed line indicates 100%, values in Table S5 and S6. (*) denotes significant difference between groups, p < 0.05; (**): p < 0.01; (***): p < 0.001; ns: not significant; Mann Whitney test (B) or one-way ANOVA with Tukey’s multiple comparisons test (D). Coloured and black bars in (A) and (C) not to scale.

We next tested a subset of AzF-containing acidic pocket variants for modulation by the gating modifier PcTx1, which was originally isolated from the venom of the *Psalmopoeus cambridgei* tarantula [41]. PcTx1 has previously been shown to increase the apparent proton affinity of both activation and steady-state desensitization of ASIC1a, resulting in inhibition or potentiation, depending on the application pH [39, 41, 53, 54]. Here, we assessed hASIC1a modulation by co-applying 100 nM PcTx1 at varying conditioning pH and compared the resulting current upon activation with pH 5.6 to the average of the preceding and following control currents after conditioning at pH 7.4 (Figure 4C). For hASIC1a WT, we observed increasing inhibition from 38.2 ± 31.7% of current remaining at pH 7.4 to 2.06 ± 2.50% at pH 7.2 (Figure 4D, Table S6). This is in agreement with previous findings that the PcTx1 IC_50_ decreases at lower pH values [39]. Channel variants with AzF in positions 344, 355 or 356 showed a similar trend (Figure 4D). In contrast, we saw potentiation for T236AzF at pH 7.4 and for D357AzF at pH 7.4 to 7.0 (Figure 4C+D). This is consistent with the observation that these variants are among those with most pronounced reduction in the pH_50_ of activation (Figure 3, S8 and Table S1, pH_50_ 6.17 ± 0.14 (n=10) and 5.66 ± 0.26 (n=10), respectively). D357AzF in particular exhibited an unusual phenotype: the first two control applications of pH 5.6 led to only very small or no detectable channel activation, but pH 5.6 after pre-application of the toxin induced a substantial inward current, after which the channels also activated in response to the following control applications. We therefore chose to evaluate PcTx1 modulation of D357AzF in more detail. Specifically, we used lower pH during conditioning and observed that at pH 6.8 and below, the variant is inhibited. In light of the strong potentiation at pH 7.0, this highlights that PcTx1 modulation of D357AzF exhibits a striking pH dependence, which far exceeds that of WT and the other mutants examined here (Figure 4D, Table S6) [39].

Overall, the APC assay established here enabled the time-efficient characterization of pharmacological modulation of selected hASIC1a variants, providing an overview on their PcTx1 modulation profile at different application pH. Together, these results confirm that hASIC1a variants containing ncAA photocrosslinkers in the acidic pocket can still be modulated by known peptide gating modifiers, opening avenues to efficiently study peptide-channel interactions with a combination of APC and photocrosslinking.

### Photocrosslinking confirms PcTx1 binding to the hASIC1a acidic pocket

Nine out of the originally targeted 12 positions around the PcTx1 binding site exhibited specific AzF incorporation (Figure 5A, left inset) and were used for photocrosslinking experiments followed by Western blotting following the workflow in Figure 5B. In parallel, six positions in the lower extracellular domain, F69, Y71, V80, D253, W287 and E413 were also replaced by AzF to confirm the specificity of potential photocrosslinking around the acidic pocket. (Figure 5A, right insets). hASIC1a variants were expressed in HEK293T ASIC-KO cells and 100 nM biotinylated PcTx1 was added before cells were exposed to UV light (365 nm) for 15 min to induce photocrosslinking. We then isolated full-length hASIC1a via a C-terminal 1D4-tag and analysed protein samples on a Western blot with antibodies against biotin and the 1D4-tag to detect PcTx1 and hASIC1a, respectively. Biotinylated PcTx1 was absent in UV-exposed hASIC1a WT and in all control positions containing AzF in the lower extracellular domain (F69, Y71, V80, D253, W287 and E413), as well as in samples containing AzF in the acidic pocket not exposed to UV light (Figure 5C). By contrast, PcTx1 was detected at four out of nine AzF-containing positions (344, 355, 356 and 357) after UV exposure, indicating covalent photocrosslinking at these positions (marked in red in Figure 5A), but at none of the five other sites in the acidic pocket tested (marked in green).

**Figure 5:**
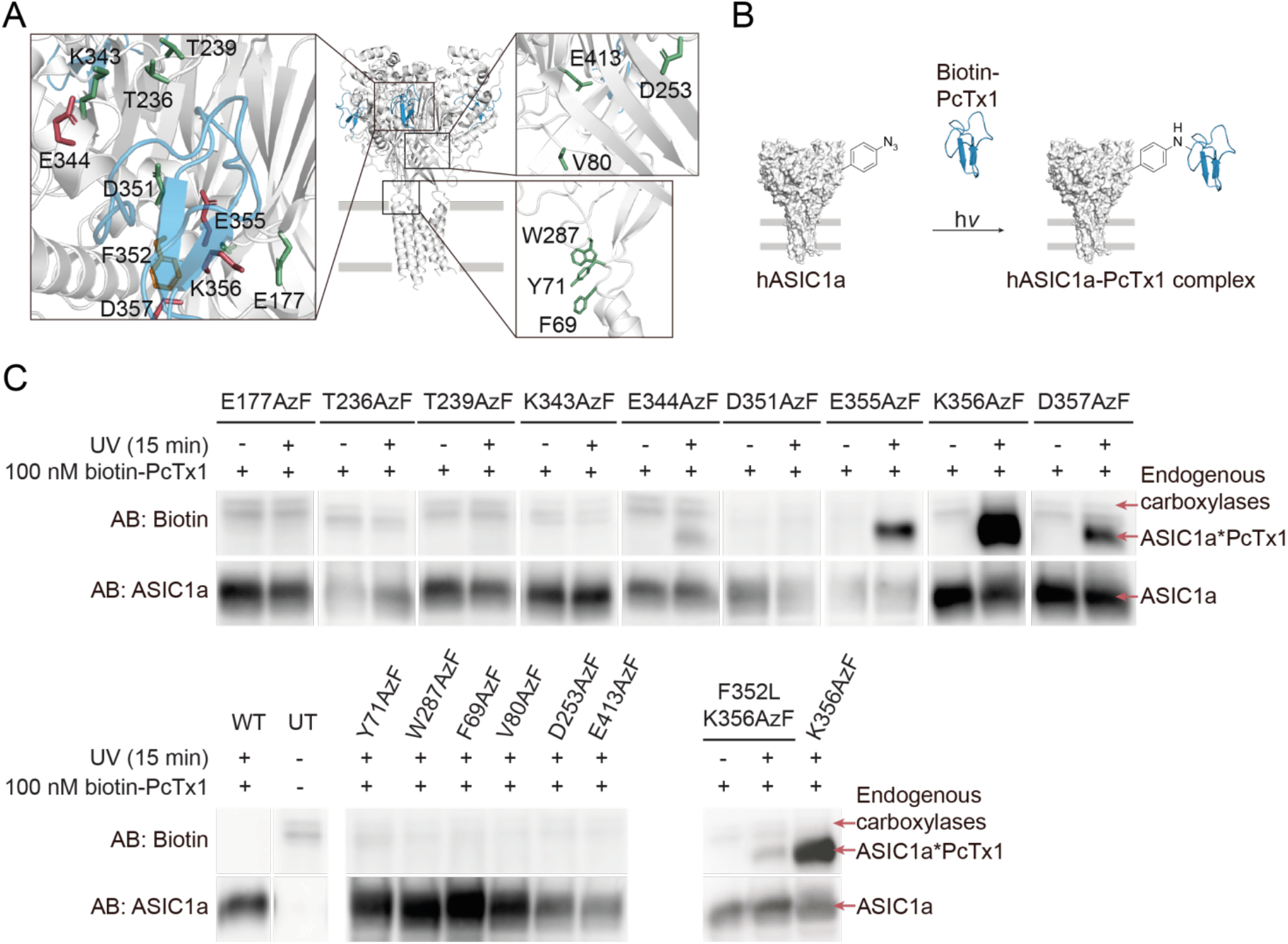
Live-cell photocrosslinking delineates the PcTx1 binding site at the ASIC1a acidic pocket. (A) Structure of cASIC1 (white) in complex with PcTx1 (blue, PDB: 4FZ0), insets show individual side chains replaced by AzF in the acidic pocket (left inset) and lower extracellular domain (right insets). Positions that crosslinked to biotin-PcTx1 are coloured red, F352 is marked in orange and positions that did not crosslink are coloured green. (B) Schematic workflow for crosslinking to biotin-PcTx1. HEK 293T ASIC1a-KO cells expressing AzF-containing hASIC1a variants are incubated with 100 nM biotin-PcTx1 and exposed to UV light for 15 min to form covalent hASIC1a-PcTx1 complexes, which are purified via a C-terminal 1D4-tag on hASIC1a and visualized via Western blotting. (C) Western blot of purified hASIC1a WT, untransfected cells (UT) and variants carrying AzF in the extracellular domain detected using the specified antibodies (AB). Biotin-PcTx1 is detected in UV-exposed samples containing AzF at positions 344, 355, 356 and 357 in the acidic pocket (coloured red in A, left inset), but not at positions 177, 236, 239, 343 or 351 (coloured green in A, left inset). PcTx1 is also absent in control samples not exposed to UV, those carrying AzF in the lower extracellular domain (right insets in A), WT or UTs. PcTx1 can be detected upon UV-exposing the toxin-insensitive F352L K356AzF double mutant (left inset in A, F352 coloured orange). Of note, the anti-biotin AB detects endogenous biotin-dependent carboxylases, which are also found in purified samples from UTs and have been described before [55, 56]. Data is representative of three individual experiments, see Figures S12-15 for original blots and crosslinking attempts with Bpa.

Previous studies have shown that the F352L mutation at the base of the acidic pocket eliminates the modulatory effect of PcTx1 on hASIC1a [57, 58], but it remained unclear if the toxin is still able to bind to hASIC1a. To test this possibility directly, we combined the F352L mutation with one of the crosslinking variants, resulting in the hASIC1a F352L K356AzF double mutant variant. Upon UV exposure, we were able to detect the PcTx1-hASIC1a complex even in the presence of the F352L mutation, albeit in lower amounts as assessed by the lower band intensity compared to the K356AzF single variant (Figure 5C, lower panel). This suggests that the F352L mutation does not eliminate toxin binding *per se*, but likely primarily abolishes the functional effects caused by PcTx1.

Attempts to photocrosslink PcTx1 using Bpa in the equivalent positions around the acidic pocket did not succeed (Figure S12). We therefore tested PcTx1 modulation of selected Bpa variants on the SyncroPatch 384PE to assess if the toxin binds to the acidic pocket when Bpa is present (Figure S13, Table S6). We observed robust inhibition at pH 7.0, indicating the interaction persists despite incorporating a bulky ncAA within the acidic pocket. However, most of the variants showed only weak modulation at the pH used during the UV exposure (7.4), which might partly explain the lack of crosslinking with Bpa.

Overall, our photocrosslinking experiments confirm that PcTx1 interacts with the acidic pocket of hASIC1a, even in the presence of a mutation that abolishes the functional effects of PcTx1.

## Discussion

### First comprehensive functional assessment of ncAA-containing ion channels on an APC platform

Since their introduction, APC platforms have greatly aided ion channel research with their high throughput capabilities [59]. However, the requirement for high transfection rates to express the ion channels of interest limits the types of experiments that can be performed with this approach. Our FACS-assisted ncAA incorporation assay represents, to our knowledge, the first example of using an APC platform to functionally interrogate ncAA-containing ion channels. By transiently transfecting the protein of interest into mammalian cells and selecting those that express all components with FACS, we circumvent the need for stable cell lines. This method therefore greatly expands the scope of experiments that can be addressed using APC-based approaches.

Our extensive scanning of 309 ncAA-containing variants emphasizes the amenability of hASIC1a to ncAA incorporation, with the highest tolerance observed for AzF (61% functional variants) followed by Bpa (50%) and Se-AbK (44%) (Figure 2E). Previous studies on incorporation of AzF and Bpa into the human serotonin transporter (hSERT) and α-amino-3-hydroxy-5-methyl-4-isoxazolepropionic acid receptor (AMPAR) also show preferred functional incorporation of AzF and attribute this to its smaller size [7, 13]. Rannversson *et al*. report lowest ncAA tolerance in the hSERT TMD (44% and 20% for AzF and Bpa, respectively), contrasting our findings in the TM segments of hASIC1a (52% and 61%). However, it should be noted that we specifically selected the outer turns of the TM helices, where the study on AMPARs observed better incorporation compared to the more tightly packed central pore [7].

Previous work on hSERT shows higher success rates for replacing aromatic vs non-aromatic side chains, a trend we only observe for Bpa. Generally, genetic encoding of ncAAs does not appear to depend on the original properties of the replaced amino acid when assessed via protein expression [14, 60]. Indeed, a systematic examination of the effect of the similarly bulky ncAA acridonylalanine on protein solubility found no correlation to amino acid conservation, hydrophobicity or accessibility, but a close dependence on the location within the overall tertiary structure [61]. Consequently, the authors suggest that scientists broaden rather than narrow screens when aiming to introduce a ncAA into a new target protein. In the present study, we cover around 20% of hASIC1a and functionally assess three different ncAAs, likely the most comprehensive investigation of genetic code expansion in a transmembrane protein to date.

### Mechanistic insights into ASIC function

A beneficial side-effect of replacing native side chains with ncAA photocrosslinkers is that, in addition to their photoactivatable properties, these bulky side chains can also inform on basic biophysical aspects of the protein domain in question. Here, we show that incorporation of bulky, non-polar side chains leads to functional channels in about 50% of all cases, and we observe a general trend towards lower apparent proton affinity in the ncAA-bearing hASIC1a channels. This is particularly evident at positions in or near the acidic pocket, where previous studies have shown that mutations to acidic side chains in thumb and finger domains result in increased pH_50_ values (reviewed in [23]). By contrast, we only found a few positions in M1 (L45, Q66, F69) that resulted in higher apparent proton affinity. This is consistent with previous work on the nearby pre-M1 region [62], as well as a number of M1 and M2 mutations that mostly resulted in left-shifted pH_50_ values [48, 63]. Together, this suggests that mutations in M1 and M2 of ASIC1a have a general tendency to increase apparent proton affinity.

Generally, we observe that the time course of current decay is relatively heterogeneous (Figure 2 and 3, S7), likely due to the slow and incomplete solution exchange (see also below). This makes an exact quantification of the changes to activation or desensitization rates difficult. Nevertheless, we observe that the same sites around the acidic pocket that show a pronounced decrease in apparent proton affinity also display a marked acceleration in current decay rates (t_1/2_ analysis in Figure 3B, S8 and Table S3). This was consistently observed at all of the eight sites around the acidic pocket assessed in Figure 3 and was independent of the nature of the incorporated ncAA. This finding is coherent with a previous study that showed the thumb domain affects rates of fast desensitization [64]. Alternatively, it is conceivable that the observed phenotype is due to greatly increased channel deactivation rates [65]. Although we cannot discriminate between these possibilities, our data clearly show that the physico-chemical properties of side chains lining the acidic pocket are a major determinant for current decay in ASIC1a.

We also noticed varying degrees of tachyphylaxis, especially when positions in the external turns of the TM helices were replaced with ncAAs (Figure S3, Table S1). In light of previous work suggesting a contribution by permeating protons and an effect of hydrophobicity of TM1 side chains on tachyphylaxis, this warrants further investigation [49, 66].

### Complex pharmacological modulation studied in ncAA-containing channels using APC

The complex pharmacological modulation pattern of hASIC1a by BigDyn and PcTx1 is notoriously challenging to study. However, we were able to optimize the APC protocols to replicate and even expand on the differential effects of this highly state-dependent peptide modulation (Figure 4). Specifically, we were able to show that despite the prominently lowered proton sensitivity of acidic pocket variants, all tested ncAA-containing hASIC1a variants retained some degree of modulation by both BigDyn and PcTx1. We observed varying degrees of BigDyn-dependent rescue from SSD for the different variants (Figure 4B). Under our conditions, rescue from SSD was incomplete when we applied 3 μM BigDyn, a concentration well above the reported EC_50_ range of 26-210 nM [17, 40]. In combination with the steep pH dependence of modulation, this resulted in considerable variability in the BigDyn modulation data, as evident by the reported range in S.D. values. While this can, at least in part, be attributed to our limited control over the BigDyn-application pH, we have made similar observations in a previous study using TEVC [17].

PcTx1 inhibited or potentiated AzF-containing hASIC1a variants in a pH dependent manner, in line with previous reports [39]. We examined a total of five variants, of which all except T236AzF also formed covalent complexes with the toxin upon UV exposure (Figure 5C). While PcTx1 still modulates and therefore interacts with hASIC1a T236AzF (Figure 4C+D), we cannot exclude that introduction of AzF at positions 177, 239, 343 or 351 prevents toxin interaction, as these variants were not assessed for PcTx1 modulation with APC and did not crosslink to the peptide upon UV exposure (Figure 5C).

### Live-cell crosslinking provides a detailed map of the PcTx1-hASIC1a interaction

The acidic pocket is now well established both as a hotspot for channel activation and as a binding site for pharmacological modulators [17, 23]. In the case of PcTx1, structural data had already outlined the toxin binding site on ASICs [52, 67], but unlike previous work, the crosslinking approach outlined in this study enables us to covalently trap ligand-channel complexes in living cells. This represents a notable advantage, especially for highly state-dependent interactions, such as those between hASIC1a and BigDyn or PcTx1. Additionally, comparing the crosslinking pattern between two ligands, the approach can indirectly inform on the varying degrees of conformational flexibility of the ligands: BigDyn is likely to be highly flexible without a strong propensity to adopt a secondary fold [68, 69], therefore samples a greater conformational space and is thus more likely to undergo covalent crosslinking at multiple sites (9/9 sites tested at the acidic pocket, [17]). By contrast, PcTx1 folds into a compact and highly stable conformation and will consequently undergo covalent crosslinking at relatively fewer sites (4/9 sites tested at the acidic pocket, Figure 5). These findings also complement an earlier investigation of the key interactions between PcTx1 and ASIC1a that concluded for the majority of contacts observed in the crystal structures to not persist during MD simulations or to not be functionally relevant for PcTx1-mediated inhibition of ASIC1a [58].

The ability to covalently trap ligand-receptor complexes offers a unique opportunity to directly assess if ASIC mutations shown to alter or abolish ligand effects still bind to the same site on the receptor. For example, the hASIC1a F352L mutation at the base of the acidic pocket is known to almost completely abolish the PcTx1-dependent modulation of ASIC1a channels [57, 58]. Yet it remained unclear if the toxin also interacts with the acidic pocket in these mutant channels. Here, we directly demonstrate that PcTx1 still binds to the acidic pocket, even at a concentration that is far too low to have a functional effect on the mutant channels (100 nM). This leads us to propose that the F352L mutation primarily affects conformational changes responsible for the PcTx1 effect on WT hASIC1a, but not toxin binding *per se*.

We note that unlike AzF, we were unable to employ Bpa for crosslinking experiments with PcTx1. To test if introduction of the more bulky photocrosslinker prevents toxin interaction, we assessed PcTx1 modulation of selected variants with APC and found robust inhibition for most variants, indicating that Bpa does not fully occlude the acidic pocket (Figure S13). We therefore speculate that steric constraints due to the positioning of the benzophenone diradical and the more selective reactivity of Bpa (reacts exclusively with C-H bonds) may play a role [70, 71]. Together, this emphasizes that screens with multiple redundant ncAAs significantly increase chances of observing successful crosslinking.

### Limitations of the outlined APC-based approach

While our work establishes that ncAA-containing ion channels can be screened on an APC platform, some limitations persist. Firstly, our present approach relies on simultaneous transfection of four plasmids (Figure 1), which can negatively impact transfection efficiency and/or result in cells not containing all four components. Careful optimization of DNA amounts and transfection conditions is therefore necessary and a revised construct design to reduce the number of plasmids could further improve yields. For example, the Plested group achieved co-expression of TAG-containing AMPAR and GFP with an internal ribosome entry site (IRES) [5, 7], while Rook *et al.* used ASIC1a with a C-terminally fused GFP-tag [8]. Furthermore, both Zhu and co-workers and Rook *et al.* created bidirectional plasmids to encode both AzF-RS or Bpa-RS and tRNA, respectively [6, 8]. This latter strategy might be particularly fruitful for the incorporation of Se-AbK, which was generally less efficient than that of AzF and Bpa (Figure 2E and Table S1), despite others reporting robust incorporation of a similar ncAA [72].

Secondly, while GFP fluorescence indicates successful transfection and ncAA incorporation and thereby increased likelihood of observing proton-gated currents in cells grown in the presence of ncAA, it is not a reliable proxy for incorporation specificity in control cells grown in the absence of ncAA. This is due to the fact that the degree of unspecific incorporation in GFP does not correlate with that of the ncAA-containing hASIC1a variants. We consistently observed GFP fluorescence in only around 2% of the control cells, independent of the co-expressed channel variant, which translated to insufficient cell numbers for APC (requires a minimal concentration of 100.000 cells/ml). Assuming that the transfection rates are similar in the presence and absence of ncAA (i.e. around 11%, Figure S1B), we concluded that recording a larger number of unsorted control cells is the more stringent approach to assess incorporation efficiency. We therefore did not subject the incorporation control cells to FACS and instead conducted APC with the entire unsorted cell population. To evaluate this strategy, we randomly selected 45 hASIC1a variants assessed for ncAA incorporation in the N-terminus, ECD or C-terminus and compared the number of wells harbouring a patched cell with >100 MΩ seal and those showing proton-gated currents in presence and absence of ncAA (Figure S1C). While the percentage of cells with current is generally lower for cells grown in the absence of ncAA, we do observe currents for those positions where incorporation is unspecific, e.g. throughout the C-terminus and in some positions in the N-terminus. For the control samples, an average of 9.8 out of 16 possible wells contained a cell with >100 MΩ seal, and we observed currents in 1.6 wells on average. We therefore conclude that despite some shortcomings, the employed strategy using non-sorted controls detects at least those positions with unspecific incorporation of >15% (i.e. 1.6/9.8).

Thirdly, while APC platforms offer unprecedented throughput and speed, there are limitations with regards to the rate and extent of perfusion exchange. This can be particularly challenging for ligand application to fast-gating ligand-gated ion channels (i.e. pH changes for ASIC1a) in general and strongly state-dependent pharmacological modulation (by e.g. BigDyn or PcTx1) in particular. Although we were able to partially overcome these issues by employing a solution stacking approach, we cannot draw detailed conclusions about activation or desensitization kinetics. Similarly, values for proton-dependent activation and especially SSD can be determined with greater precision using TEVC or manual patch-clamp electrophysiology. However, note that the values reported here are generally in agreement with previous reports, both with regards to WT values [46, 47] and relative shifts caused by mutations, i.e. in the acidic pocket [23].

Lastly, limitations arise from the accessibility and running costs of APC platforms compared to conventional patch-clamp set ups. But we anticipate that the establishment of academic core facilities for high-throughput electrophysiology (e.g. Northwestern University, Il, US; University of Nantes, France; Illawarra Health and Medical Research Institute, Wollongong, Australia) and collaborations between academia and industry (this study, [73–76]) will likely contribute to a broader accessibility. This is also evident from the rising number of publications involving APC (currently >80 publications according to vendor information).

### Conclusions and outlook

The ability to functionally screen ncAA-containing ion channels on APC platforms has the potential to greatly expand the use of ncAAs in both academic and industry settings. The intrinsically high throughput enables rapid assessment of incorporation efficiencies, functional properties and even complex pharmacological modulation. In principle, the approach can be used for both site-specific (this study) and global ncAA incorporation [77, 78], thus further increasing the number and type of chemical modifications that can be introduced. In the case of incorporation of photocrosslinking ncAAs, the approach can be exploited to crosslink to peptides (Figure 5, [17]), small molecules [13] or establish intra-protein crosslinking, including in protein complexes [8, 9]. Furthermore, the recently developed ability for on-chip optostimulation on related APC platforms [79] offers exciting prospects for potentially conducting UV-mediated crosslinking during live APC experiments in the future. Paired with MS and/or biochemical approaches [80, 81], the overall strategy could also be expanded to define interaction sites of unknown or known protein-protein interactions. Given that there are now well over 100 different ncAAs available for incorporation into proteins in mammalian cells [1, 82], the above approach will enable the efficient study of ion channels endowed with a wide range of properties or functionalities.

## Material and Methods

### Molecular biology

The complementary DNA (cDNA) encoding human ASIC1a (hASIC1a) was kindly provided by Dr. Stephan Kellenberger. Plasmids containing AzF-RS, Bpa-RS and tRNA were gifts from Dr. Thomas P. Sakmar [43]. AbK-RS and tRNA_pyl_ in pcDNA3.1 were kindly provided by Dr. Chris Ahern [44]. The dominant negative eukaryotic release factor (DN-eRF) was a gift from Dr. William Zagotta [83]. Plasmids containing rat GluA2 Q607 Y533TAG or S729TAG were gifts from Dr. Andrew Plested [5, 7], rat GluA2 Q607 WT was kindly provided by Dr. Anders Skov Kristensen. Rat P2X2 WT 3T was a gift from Dr. Thomas Grutter [50], the K296TAG variant was generated in-house.

Site-directed mutagenesis was performed using PfuUltraII Fusion polymerase (Agilent, Denmark) and custom DNA mutagenesis primers (Eurofins Genomics, Germany). All sequences were confirmed by sequencing of the full coding frame (Eurofins Genomics). For hASIC1a constructs, a C-terminal 1D4-tag was added for protein purification and Western blot detection and two silent mutations were inserted at V10 and L30 to reduce the risk of potential reinitiation [84].

### Cell culture and transfection

HEK 293T cells (ATCC^®^), in which endogenous hASIC1a was removed by CRISPR/Cas9 [17], were grown in monolayer in T75 or T175 flasks (Orange Scientific, Belgium) in DMEM (Gibco, Denmark) supplemented with 10 % FBS (Thermo Fisher Scientific, Denmark) and 1 % penicillin-streptomycin (Thermo Fisher Scientific) and incubated at 37 °C in a humidified 5 % CO_2_ atmosphere. For APC experiments, cells were seeded into six-well plates (Orange Scientific) at a density of 300.000 cells/well and transfected the next day with Trans-IT LT1 (Mirus, WI, USA) and 1:1:1:1 μg DNA encoding hASIC1a TAG variants, ncAA-RS, tRNA and eGFP Y40TAG or Y151TAG, respectively. For the WT control, cells were transfected with 1 μg hASIC1a WT and 0.3 μg eGFP WT. Six hours after transfection, cell medium was replaced with supplemented DMEM containing 10 μM AzF- or Bpa-methylester (synthesis in SI) or 100 μM Se-AbK (custom-synthesized by ChiroBlock, Germany). FACS and APC recordings were performed 48 hours after transfection. The same procedure was used for GluA2 and P2X2R recordings.

For crosslinking studies, cells were seeded into 15 cm dishes (VWR, Denmark) at a density of 5-7 million cells and transfected the next day with PEI (Polysciences, Germany) and 16:4:4:8 μg DNA encoding hASIC1a TAG variants, AzF-RS, tRNA and DN-eRF, respectively. For WT controls, 2 million cells were seeded into a 10 cm dish (VWR) and transfected with 8 μg hASIC1a WT. Six hours after transfection, cell medium was replaced with supplemented DMEM containing 0.5 mM AzF (Chem Impex, IL, USA) or 1 mM Bpa (Bachem Bio, Switzerland) and crosslinking studies were performed 48 hours after transfection. Please note that for crosslinking studies followed by Western blot, the free acid version of the ncAAs was used to increase protein yields.

### FACS

HEK 293T cells were washed with PBS, treated with Accutase (Sigma Aldrich, Denmark) or Trypsin-EDTA (Thermo Fisher Scientific), pooled and centrifuged at 1000 rpm for 5 min. They were resuspended in 350 μl of a 1:1 mixture of serum-free Hams F-12 nutrient mixture and extracellular patch-clamp solution (140 mM NaCl, 4 mM KCl, 1 mM MgCl_2_, 2 mM CaCl_2_, 10 mM HEPES, pH 7.4) supplemented with 20 mM HEPES and transported to the FACS core facility at ambient temperature. A FACSAria I or III (BD Biosciences, CA, USA) with a 70 μm nozzle was used to sort cells for singularity, size and GFP fluorescence (Excitation 488 nm, Emission 502 nm (low pass) and 530/30 nm (band pass)). Cells were filtered through a sterile 50 μm cup filcon (BD Biosciences) directly before sorting to prevent clogging of the nozzle. The WT control was used to set the fluorescence cutoff between GFP-positive and GFP-negative populations and to check the purity of the sort before sorting 1 million GFP-positive cells for subsequent patch-clamp experiments. Where possible, a minimum of 200000 GFP-positive cells were collected for hASIC1a TAG variants grown in presence of ncAA, while controls grown in absence of ncAA and untransfected cells were not sorted. Cells were collected in 1.5 ml tubes containing the 1:1 mixture mentioned above and transported to the APC instrument at ambient temperature.

### Automated patch-clamp

Automated whole-cell patch-clamp recordings were conducted on a SyncroPatch 384PE (Nanion Technologies, Germany) directly after FACS sorting. Cells were loaded into a teflon-coated plastic boat at concentrations of 1 million cells/ml (WT, controls grown in absence of ncAA and untransfected cells) or 200000–400000 cells/ml (variants grown in presence of ncAA) and incubated at 20 °C and 200 rpm. For patch-clamp recordings, a NPC^®^-384 medium resistance single hole chip (Nanion Technologies) was filled with intracellular solution (120 mM KF, 20 mM KCl, 10 mM HEPES, pH 7.2) and extracellular solution (140 mM NaCl, 4 mM KCl, 1 mM MgCl2, 2 mM CaCl_2_, 10 mM HEPES, pH 7.4). 30 μl of cells were loaded into each well and the cells were caught on the holes by brief application of −200 mbar pressure and washed with 30 μl seal enhancer solution (extracellular solution with 8 mM Ca^2+^) under a holding pressure of −50 mbar. After a wash step with extracellular solution, two more pulses of −200 mbar were applied to reach whole cell configuration and the cells were clamped at 0 mV under atmospheric pressure (Figure S9A). For recordings of concentration-response curves, extracellular solutions at different pH were applied using a liquid stacking approach. Briefly, pipette tips were loaded with 45 μl of pH 7.4 wash solution followed by 5 μl of activating extracellular solution (pH 7.2-4.8). For each sweep, the baseline current was recorded for 1 sec before application of the 5 μl activating solution, while the pH 7.4 wash solution was dispensed with a delay of 5 sec to allow for recording of channel opening and desensitization in the presence of ligand. The second dispension was directly followed by aspiration of liquid and a second wash step with pH 7.4 before application of the next activating pH (interval between stimuli 140 sec, Figure S9B).

For SSD curve recordings, cells were exposed to an activating pH of 5.6 using the stacked addition protocol described above, while the conditioning pH was varied (pH 7.6-6.4). The open-well system of the SyncroPatch 384PE does not allow a single exchange of the entire liquid surrounding the cell, as this would result in destabilization or loss of the seal. Instead, the conditioning pH was adjusted stepwise by repeated addition and removal of 50% of the solution in the well, leading to 6 min conditioning intervals between stimuli (Figure S9C). While this process was simulated at the pH meter to determine the apparent conditioning pH, small variations may occur due to mixing effects. The authors note that APC instruments operating with microfluidic flow channels might offer superior control of the conditioning pH. At the end of each SSD curve recording, a control application of pH 5.6 after conditioning pH 7.6 was used to assess the extent of current rescue and exclude cells that did not recover from SSD.

For peptide modulation experiments, 0.1 % (w/v) bovine serum albumin (BSA, Sigma Aldrich) was added to the conditioning solutions to reduce peptide loss on boat and tip surfaces. To investigate modulation by BigDyn (synthesis described in [17]), cells were first exposed to two activations with pH 5.6 after conditioning at pH 7.6 to determine the control current, followed by two rounds of activation after 2 min conditioning with a pH that induces SSD (total interval between stimuli: 8 min, due to the conditioning protocol described above) and a control activation to evaluate current recovery. For half of the cell population, 3 μM BigDyn were co-applied during the second conditioning period to measure rescue from SSD. This assessment of SSD and recovery was repeated with peptide co-application during the first SSD-conditioning to also evaluate peptide wash out. To assess modulation by PcTx1 (Alomone labs, Israel, >95% purity), cells were exposed to two control measurements of activation with pH 5.6 after conditioning at pH 7.4 (interval 3.75 min), followed by pH 5.6 activation after incubation with 100 nM PcTx1 at varying pH (pH 7.4-7.0) for 2 min (total interval between stimuli 7 min), as well as two further controls to assess recovery from modulation. For recordings on GluA2 and P2X2R variants, cells were clamped at −60 mV and currents activated by application of 30 μM and 300 μM/10 mM ATP or glutamate, respectively. Cells expressing GluA2 were pre-incubated with 100 μM cyclothiazide (in 0.8% (v/v) DMSO) for 60 sec before activation to slow desensitization (total interval between stimuli: 220 sec) [51].

### Data analysis

Current traces were acquired at 2 kHz and filtered in the DataControl384 software using a Butterworth 4th order low pass filter at 45 Hz to remove solution artefacts. Only cells with initial seals >100 MΩ were considered for biophysical characterization using GraphPad Prism 7 or 8, while wells with lower seals, no current or no caught cell were excluded. The relatively low seal cutoff in combination with the large proton-gated currents (up to 10 nA) recorded for WT and some of the ncAA-containing variants resulted in suboptimal voltage-clamp conditions for a subpopulation of cells, as also apparent from the current shapes. However, we have no evidence that this adversely affected activation parameters or pharmacological modulation. Where possible, APC data was pooled from a minimum of three cells and two separate recording days. On several occasions, an *n* of five or more was acquired during the first screening trial, in which case the experiment was not repeated. Current sizes were normalized to the respective control currents and half-maximal concentrations (EC_50_ values) and Hill slopes (nH) calculated using equation (1). pH_50_ values were calculated in Excel using equation (2). All values are expressed as mean ± S.D. (Standard Deviation). The extent of tachyphylaxis for each recording was calculated by subtraction of the normalized current at lowest pH from the normalized maximal current (> 20 % tachyphylaxis is marked by (^#^)). Bar graphs and dot plots were made using GraphPad Prism 7 or 8 and SigmaPlot 13.0, while current traces were exported to Clampfit 10.5 and Adobe Illustrator CC 2019.

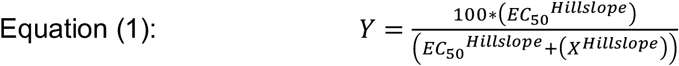

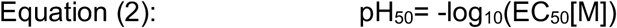

Mean current sizes and pH_50_ values of different cell lines and constructs were compared using student’s t-test, Mann Whitney test or one-way ANOVA followed by Tukey’s multiple comparisons test.

### Crosslinking studies, protein purification, western blotting

Cells were washed with PBS and dislodged using cell scrapers (Orange Scientific). After centrifugation (1000 rpm, 5 min), cell pellets were resuspended in 1 mL PBS pH 7.4 containing 100 nM biotinyl-PcTx1 (Phoenix Pharmaceuticals, CA, USA) and transferred into 12 well plates (Orange Scientific). Cells were placed on ice and crosslinked at a distance of 7-10 cm to a Maxima ML-3500 S UV-A light source (Spectronics corporation, 365 nm) for 15 min (AzF) or up to 60 min (Bpa). Control samples without UV exposure were kept at 4 °C. After crosslinking, cells were centrifuged (1000 rpm, 5 min) and resuspended in 1 mL solubilisation buffer (50 mM Tris-HCl, 145 mM NaCl, 5 mM EDTA, 2 mM DDM, pH 7.5) supplemented with cOmplete™ EDTA-free protease inhibitor cocktail (Sigma Aldrich). Cells were lysed (2 h, 4 °C) and centrifuged for 30 min (18000 g/4 °C). In parallel, 40 μL Dynabeads Protein G (Thermo Fisher Scientific) were washed with 200 μL PBS/0.2 mM DDM and incubated with 4 μg RHO 1D4 antibody (University of British Columbia) in 50 μL PBS/0.2 mM DDM on a ferris wheel (VWR, 30 min). After washing the beads PBS/0.2 mM DDM (with 200 μL), the cell lysate supernatant was incubated with the beads on a ferris wheel (4 °C, 90 min). Beads were washed with 200 μL PBS three times to remove nonspecifically bound proteins and incubated in 25 μL elution buffer (2:1 mixture between 50 mM glycine, pH 2.8 and 62.5 mM Tris-HCl, 2.5 % SDS, 10 % Glycerol, pH 6.8) supplemented with 80 mM DTT at 70 °C for 10 min. Protein samples (12 μL) were mixed with 3 μL 5 M DTT and 5 μL 4x NuPAGE™ LDS sample buffer (Thermo Fisher Scientific) and incubated (95 °C, 20 min) before SDS-PAGE using 3–8 % Tris-Acetate protein gels (Thermo Fisher Scientific). After transfer onto PVDF membranes (iBlot 2 Dry Blotting System, Thermo Fisher Scientific) and blocking in TBST/3% non-fat dry milk for 1 hour, hASIC1a was detected using RHO 1D4 antibody (1 μg/μL, University of British Columbia) and 1:5000 goat anti-mouse IgG HRP-conjugate (Thermo Fisher Scientific). Biotinyl-PcTx1 was detected using 1:1000 rabbit anti-biotin antibody (abcam, UK) and 1:5000 goat anti-rabbit IgG HRP-conjugate (Promega, Denmark). Samples used for incorporation controls were treated as above, but were not exposed to UV light.

## Acknowledgements

We acknowledge the FACS core facility at the Biotech Research & Innovation Center (University of Copenhagen) for technical support. We thank Dr. Iacopo Galleano for the synthesis of AzF- and Bpa-ME, Dr. Christian Bernsen Borg for the synthesis of big dynorphin and members of the Pless laboratory for comments on the manuscript. Figure 1 was created with BioRender.com.

