## Supplementary information for "High-throughput characterization of photocrosslinker-bearing ion channel variants to map residues critical for function and pharmacology"

- Supplementary methods
- Supplementary figures 1-15
- Supplementary tables 1-6
- Supplementary references

### Supplementary methods

**Synthesis of ncAA-MEs.** AzF (4-Azido-L-phenylalanine) and Bpa (4-Benzoyl-L-phenylalanine) were purchased from Chem Impex (IL, USA) and Bachem Bio (Switzerland), respectively. For the synthesis of ncAA-methylesters [1], TMSCl (chlorotrimethylsilane, 2 equivalents) was added to the amino acid in a round-bottom flask. Anhydrous methanol was added to dissolve the starting material and the reaction mixture was stirred at room temperature over night under nitrogen atmosphere. TMSCl (1 equiv.) was added every 24h until complete conversion of starting material (reaction monitored by LC-MS (liquid-chromatography - mass spectrometry)). The solvents were evaporated yielding an off-white solid, which was purified by preparative HPLC (high-performance liquid chromatography) when necessary. ncAA-methylesters were dissolved in DMSO (dimethyl sulfoxide) and stock solutions were stored at -20 ˚C.

**Mass spectrometry.** For filter-assisted in-solution digestion, eluted protein samples (25 µl) were incubated with 250 µl UA solution (8 M Urea in 0.1 M Tris/HCl (pH 8.5), 10 mM TCEP) for 10 min and passed through a 0.5 ml Amicon Ultra centrifugal filter (30 kD cutoff, Sigma Aldrich, Germany) by centrifugation at 14000 g for 10 min. Samples were washed with 200 µl UA by centrifugation and incubated with 100 µl iodoacetamide solution (50 mM iodoacetamide in UB) at RT for 20 min in the dark. After centrifugation, samples were washed twice with 200 µl UB (8 M Urea in 0.1 M Tris/HCl (pH 8.5)) and twice with 200 µl of 50 mM ammonium bicarbonate before incubation with 40 µl of chymotrypsin (0.5 µg) in 25 mM ammonium bicarbonate at 37 °C over night. The next morning, 20 µl trypsin (0.25 µg) in 25 mM ammonium bicarbonate were added and the samples were incubated at 37 °C for 4 hours. Using a new collection tube, samples were centrifuged and the columns washed with 50 µl 0.5 M NaCl. After addition of 10 µl 10 % (v/v) trifluoroacetic acid, peptide solutions were concentrated to 40 µl on a Savant SPD1010 SpeedVac Concentrator (Thermo Fisher Scientific, Germany) and loaded onto the LC-MS system.

Peptide solutions were analyzed by liquid chromatography followed by tandem mass spectrometry (LC-MS/MS) on Ultimate 3000 RSLC nano-HPLC systems coupled to an Orbitrap Fusion mass spectrometer (all from Thermo Fisher Scientific)[2]. The mass spectrometer was equipped with a nano-ESI source (Nanospray Flex or EASY-Spray™ ion source, Thermo Fisher Scientific) and external column heater (Phoenix S&T, PA, USA) to enable the use of self-packed emitter columns (PicoFrit, New Objective, MA, USA). Samples were loaded onto an RP (reverse phase) C18 pre-column (Acclaim PepMap, 300 µm × 5 mm, 5 µm, 100 Å, Thermo Fisher Scientific) at a flow rate of 30 µl/min and washed with 0.1 % (v/v) TFA for 15 min at 30 µL/min before elution and separation on a self-packed RP C18 separation column (PicoFrit, 75 µM × 250–500 mm, 15 µm tip diameter, packed with *ReproSil-Pur* C18-AQ, 1.9 µm, 120 Å, Dr. Maisch, Germany) equilibrated with 3%

solvent B (solvent A: 0.1 % (v/v) FA (formic acid), solvent B: ACN, 0.08 % (v/v) FA). A gradient from 3–40 % solvent B within 90 min at 300 nl/min was used to elute peptides from the separation column. A voltage of 1.9 kV was applied between the column and the mass spectrometer entrance for positive ionization. Data were acquired in data-dependent MS/MS mode using HCD (high energy collisional dissociation, stepped normalized collision energies (NCE): 28 %) for fragmentation. For data acquisition, each high-resolution full scan (*m/z* 300 to 1500, R = 120000) in the Orbitrap was followed by high-resolution product ion scans (R = 15000, minimum charge states 2+ to 6+) within 5 seconds, starting with the most intense signal in the full scan mass spectrum (isolation window 2 Th); the target value and maximum accumulation time were 50000 and 200 ms. Dynamic exclusion (duration 60 s, window ± 2 ppm) was enabled. The Xcalibur software (version 4.1, Thermo Fisher Scientific) was used for data acquisition and raw data inspection.

Raw data generated by LC-MS/MS was processed using the Thermo Proteome Discoverer (version 2.0.0.802, Thermo Fisher Scientific) by matching MS and MS/MS data to the human reference proteome downloaded from the UniProt database (http://www.uniprot.org, FASTA file) or an *in-house* database containing the amino acid sequence of hASIC1a as well as common contaminants. For identification of proteins and evaluation of the sequence coverage, the Sequest search engine was used (maximum mass differences of 10 ppm and 0.02 Da for MS and MS/MS data). Further parameters were unspecific enzymatic cleavage, alkylation of cysteines as a static modification and oxidation of methionines as well as replacement of alanine by Bpa as variable modifications. Mass spectra were exported from Thermo Proteome Discoverer and further annotated in Illustrator CC 2019.

**Supplementary figures S1-15**

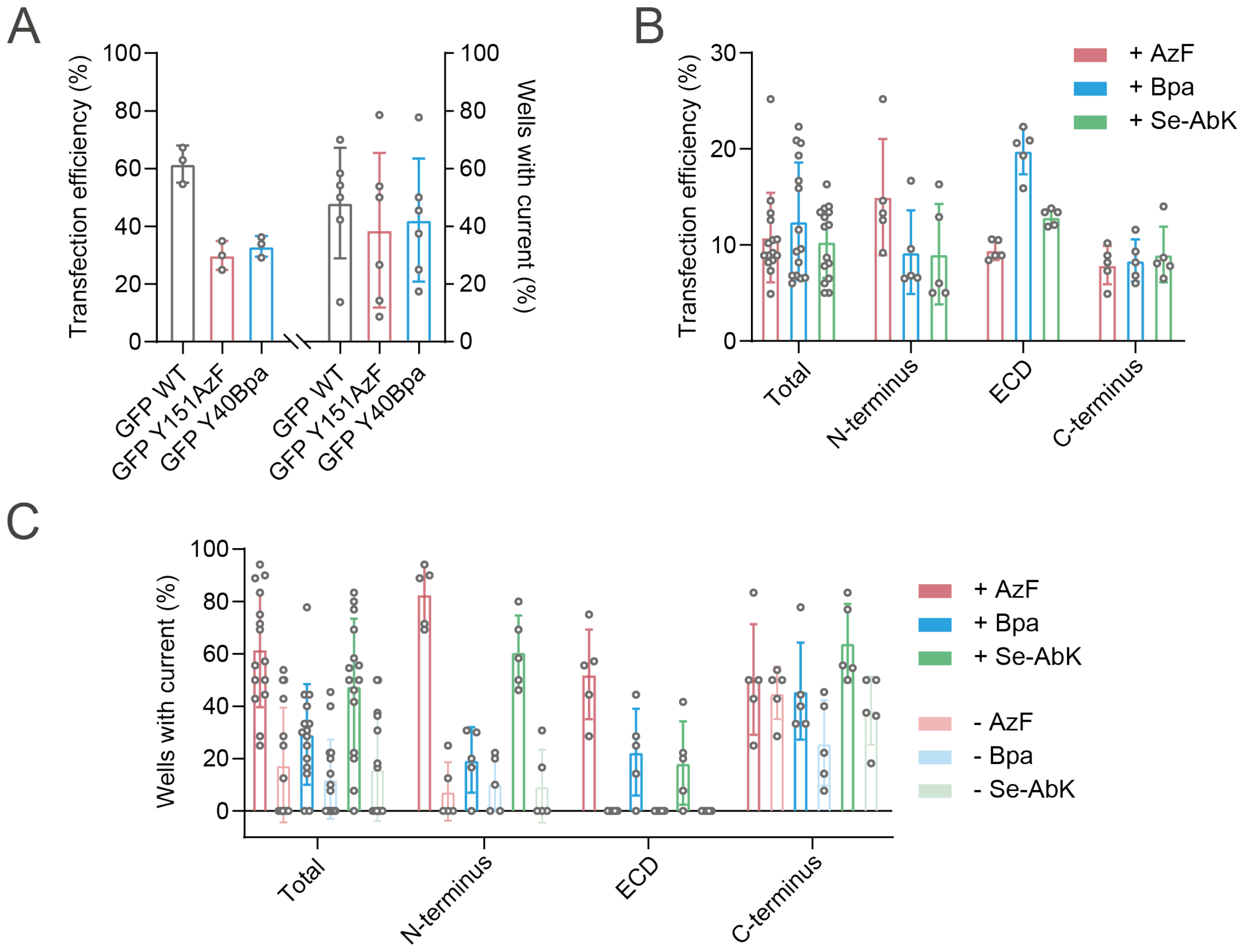

Figure S1: FACS prior to APC increases current density on the 384-well chip. (A) Transfection efficiency and percentage of wells showing proton-gated currents (of wells harbouring a patched cell with >100 MΩ seal) for hASIC1a WT co-transfected with WT GFP, GFP Y151AzF or GFP Y40Bpa. While the transfection efficiency is reduced considerably from 61.6% to 29.9% and 33.1% upon co-transfection of GFP Y151AzF or GFP Y40Bpa, respectively, the decrease in mean current density is less pronounced from around 48% to 39% and 42%. This illustrates that non-sense suppression in GFP results in a decrease in apparent TE compared to WT and shows that both WT and TAG-containing GFP can be used as a reporter to enrich transfected cells for APC. (B) Transfection efficiency of 45 randomly selected hASIC1a variants assessed for ncAA incorporation in the N-terminus, ECD or C-terminus. Of note, only cells grown in the presence of ncAA were FACS-sorted and assessed for transfection efficiency. (C) Percentage of wells showing proton-gated currents (of wells harbouring a patched cell with >100 MΩ seal). Cells grown in the absence of ncAA (lighter shades in right panel) show currents in 0-50% of the wells, depending on incorporation specificity, which varies in the different protein domains. All values shown as mean ± S.D..

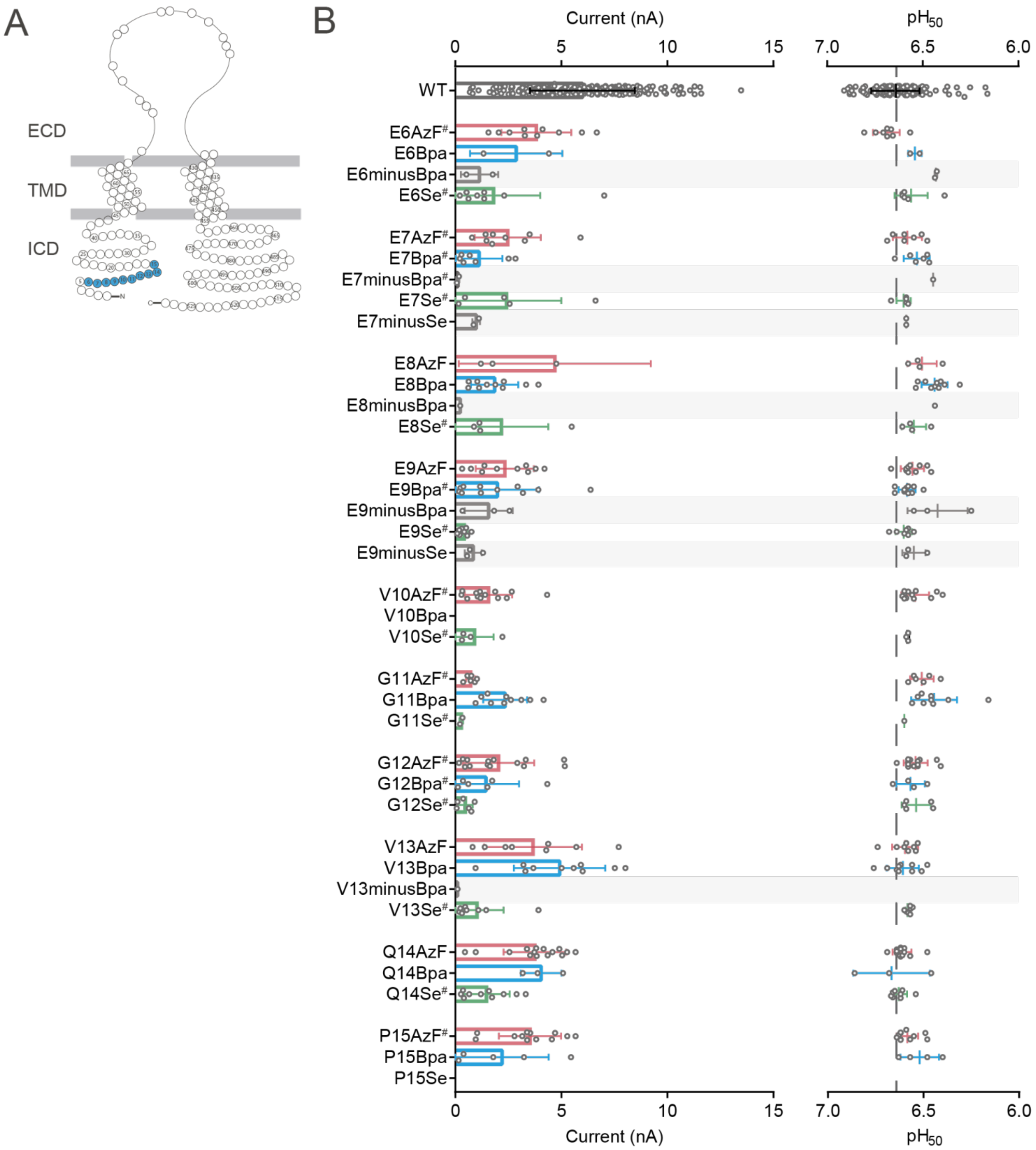

Figure S2 (including next 3 pages): Incorporation of ncAA photocrosslinkers into the hASIC1a N-terminus is specific from position 10 onwards and mostly produces variants with WT-like properties. (A) Snake plots with tested positions marked in blue. (B) Dot plots comparing peak current sizes (left) and $pH_{50}$ (right), bars indicate mean ± S.D., (#) marks >20% tachyphylaxis (see also Table S1). For variants expressed in the absence of ncAAs that yielded currents, results are marked by underlying grey bars.

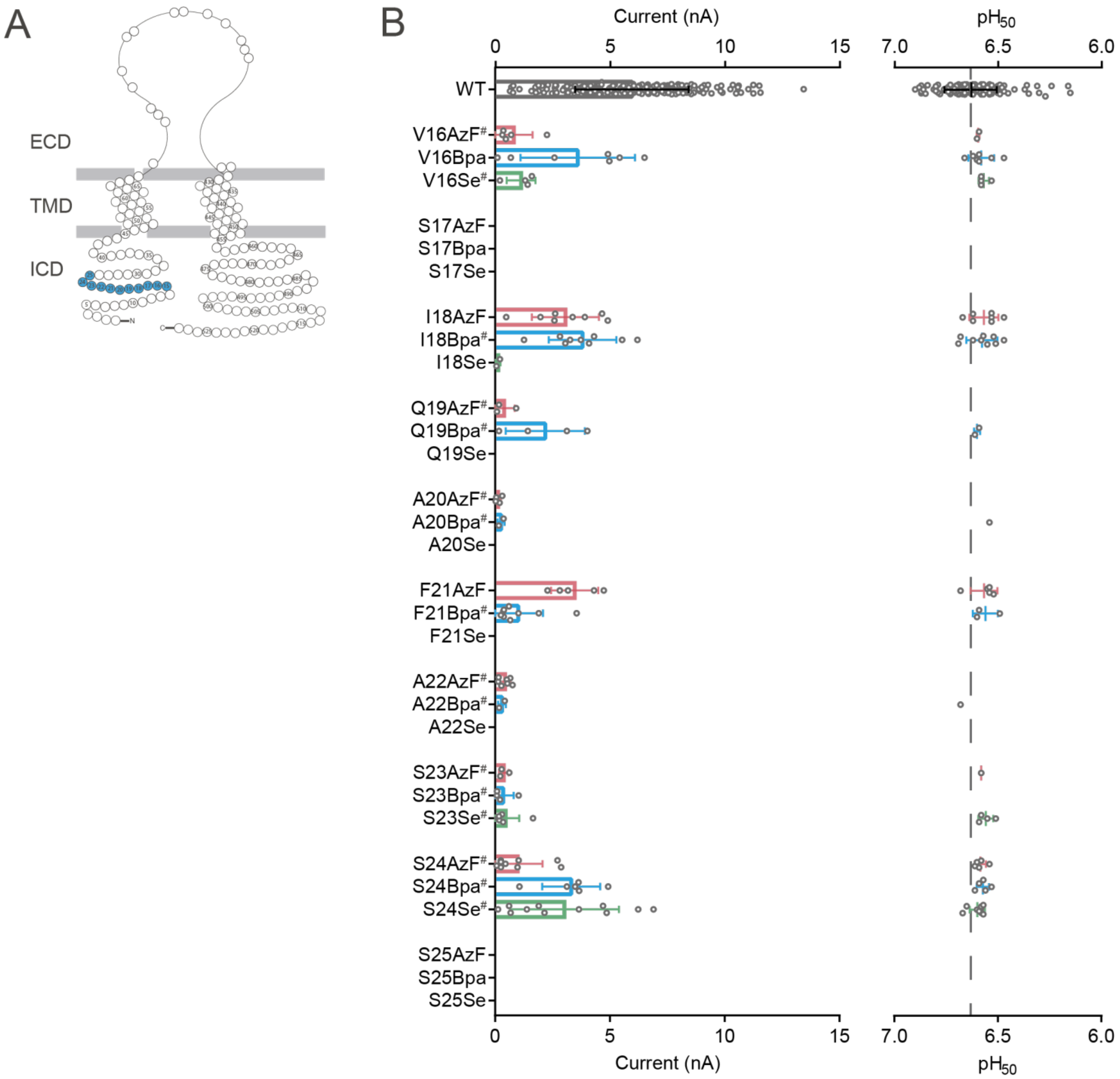
A
ECD
TMD
ICD
B
Current (nA)
pH50
WT
V16AzF#
V16Bpa
V16Se#
S17AzF
S17Bpa
S17Se
I18AzF
I18Bpa#
I18Se
Q19AzF#
Q19Bpa#
Q19Se
A20AzF#
A20Bpa#
A20Se
F21AzF
F21Bpa#
F21Se
A22AzF#
A22Bpa#
A22Se
S23AzF#
S23Bpa#
S23Se#
S24AzF#
S24Bpa#
S24Se#
S25AzF
S25Bpa
S25Se

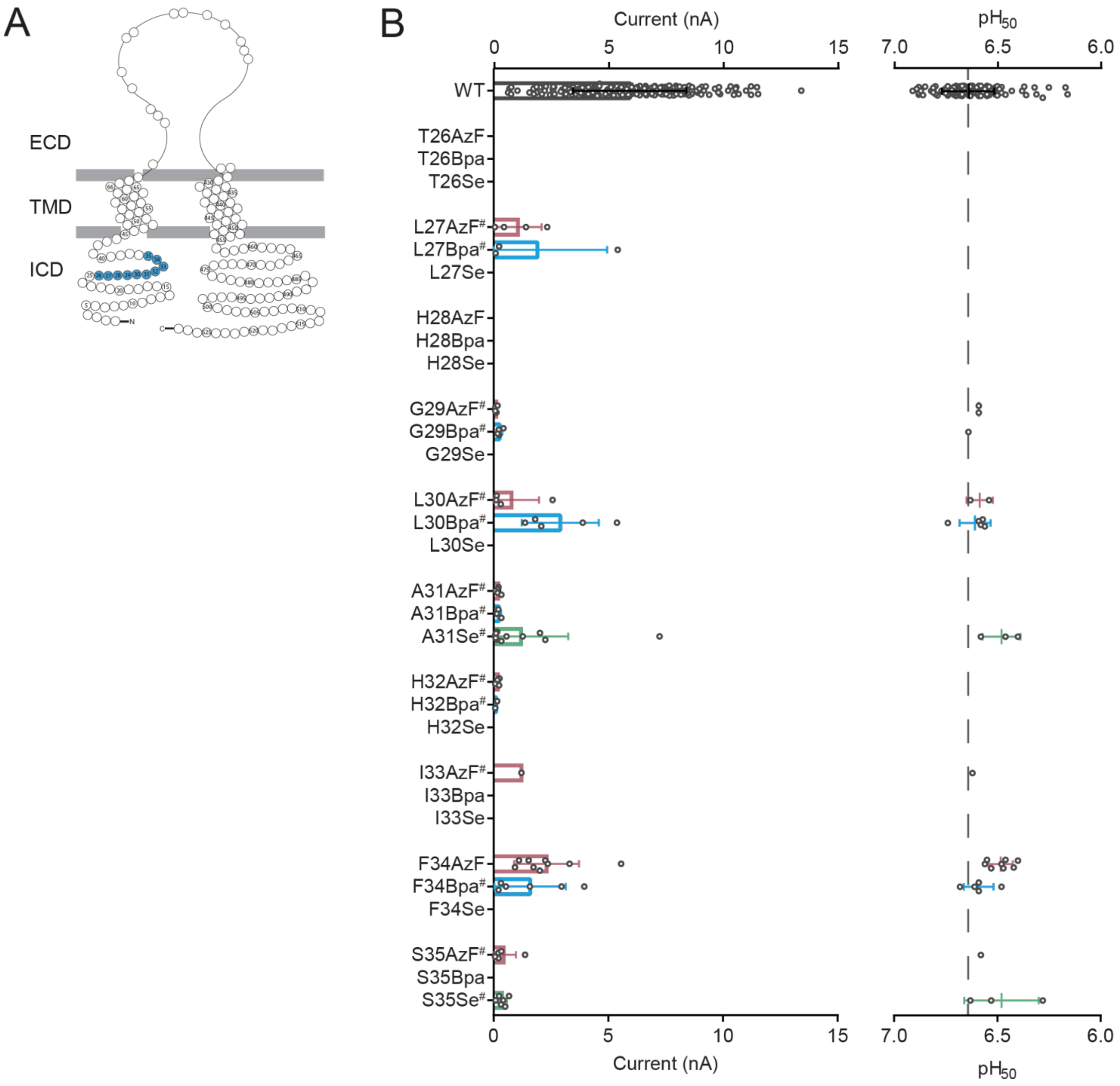
A
ECD
TMD
ICD
B
Current (nA)
pH50
WT
T26AzF
T26Bpa
T26Se
L27AzF#
L27Bpa#
L27Se
H28AzF
H28Bpa
H28Se
G29AzF#
G29Bpa#
G29Se
L30AzF#
L30Bpa#
L30Se
A31AzF#
A31Bpa#
A31Se#
H32AzF#
H32Bpa#
H32Se
I33AzF#
I33Bpa
I33Se
F34AzF
F34Bpa#
F34Se
S35AzF#
S35Bpa
S35Se#

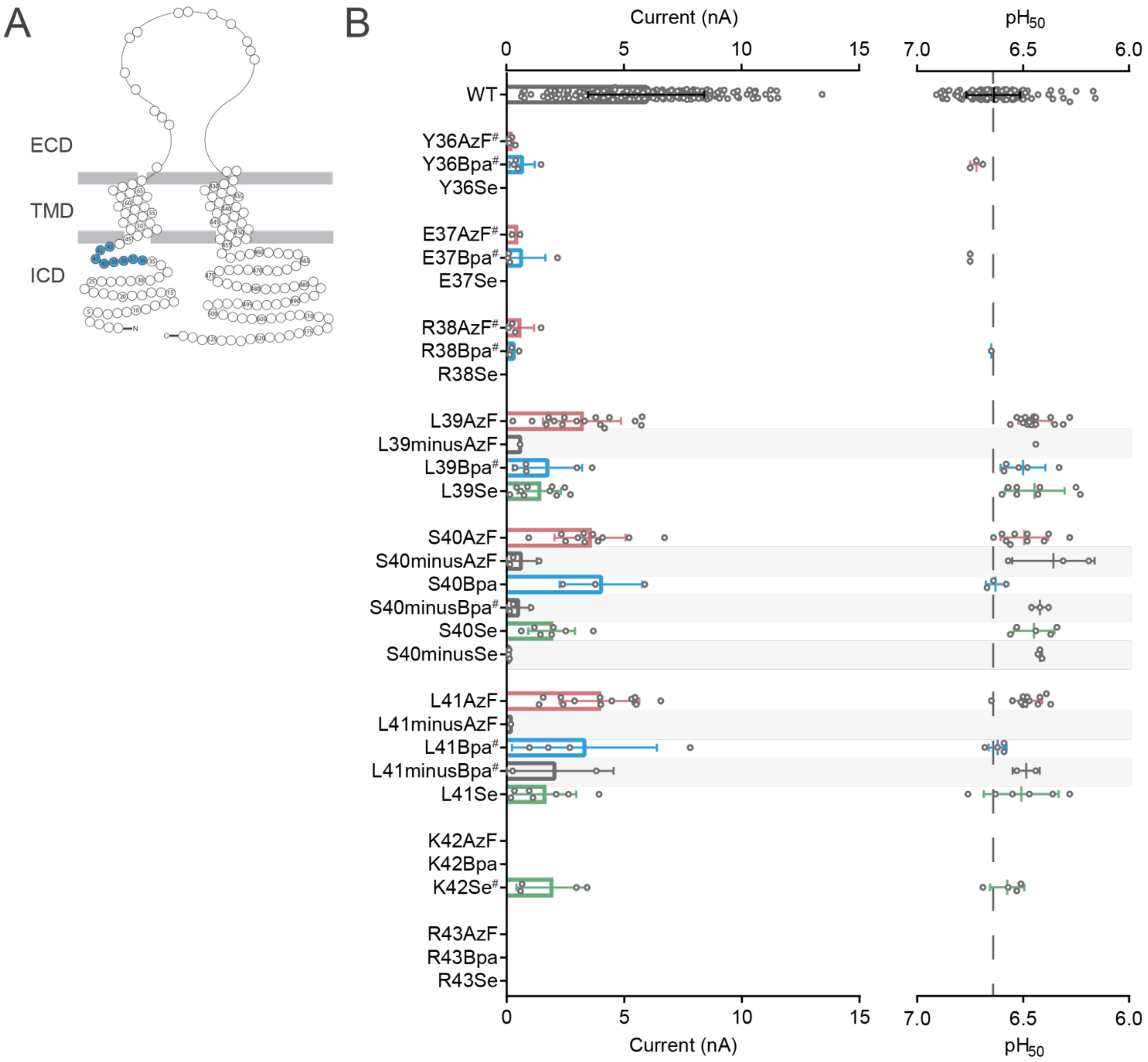

Figure S3 (next 2 pages): In the hASIC1a outer transmembrane helix loops and interface region, incorporation of ncAA photocrosslinkers is better tolerated in the M1 than the M2 helix and produces variants with varying degrees of tachyphylaxis. (A) Snake plots with tested positions marked in blue. (B) Several variants, e.g. A47AzF, Y68Bpa, G340Se and Y458AzF, undergo tachyphylaxis after reaching the peak current. (C) Dot plots comparing $pH_{50}$ (left) and peak current sizes (right), bars indicate mean ± S.D., (#) marks >20% tachyphylaxis (see also Table S1). For variants expressed in the absence of ncAAs that yielded currents, results are marked by underlying grey bars.

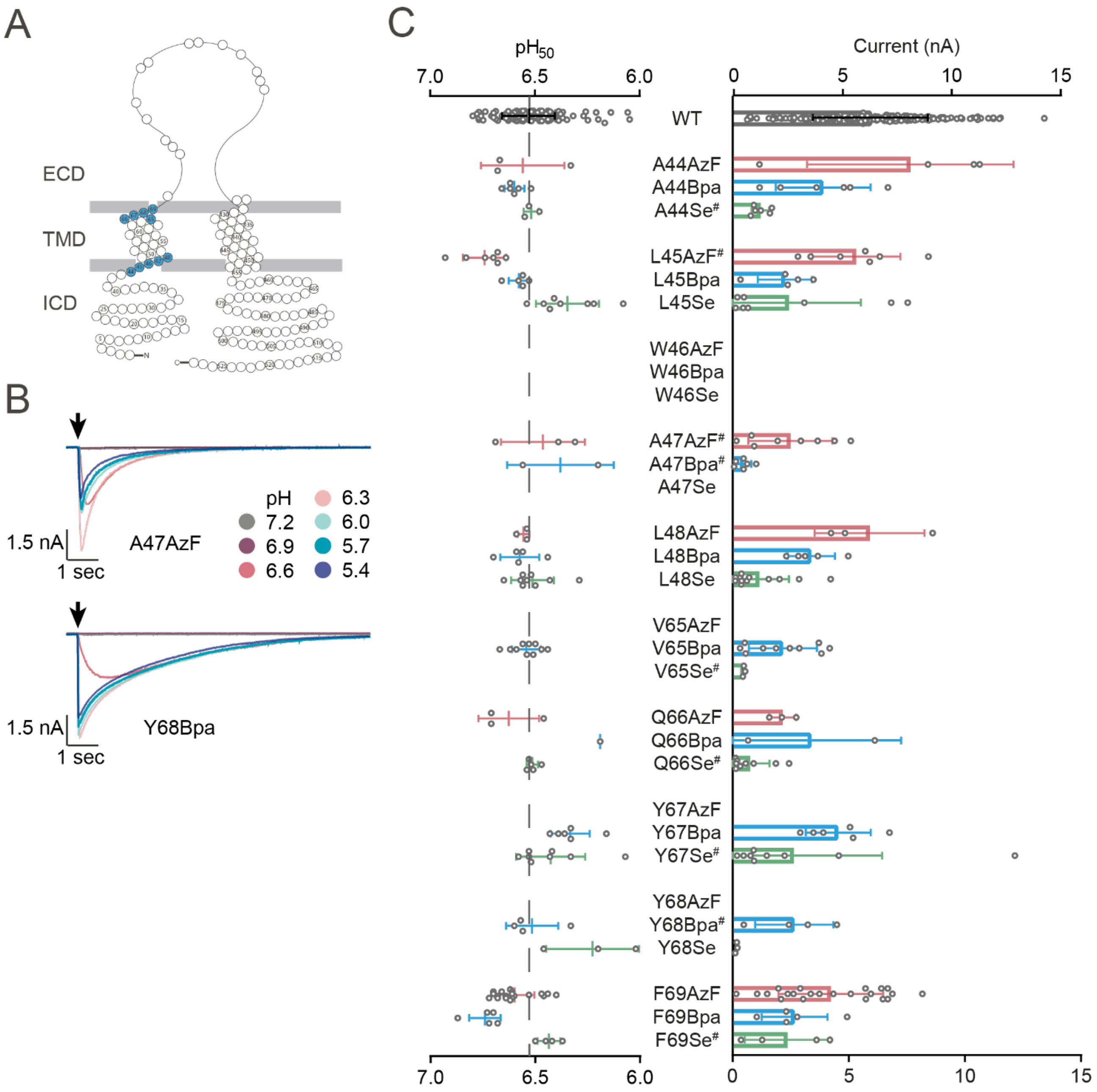
A
ECD
TMD
ICD
B
A47AzF
1.5 nA
1 sec
pH
7.2
6.9
6.6
6.3
6.0
5.7
5.4
Y68Bpa
1.5 nA
1 sec
C
pH50
7.0
6.5
6.0
Current (nA)
0
5
10
15
WT
A44AzF
A44Bpa
A44Se#
L45AzF#
L45Bpa
L45Se
W46AzF
W46Bpa
W46Se
A47AzF#
A47Bpa#
A47Se
L48AzF
L48Bpa
L48Se
V65AzF
V65Bpa
V65Se#
Q66AzF
Q66Bpa
Q66Se#
Y67AzF
Y67Bpa
Y67Se#
Y68AzF
Y68Bpa#
Y68Se
F69AzF
F69Bpa
F69Se#

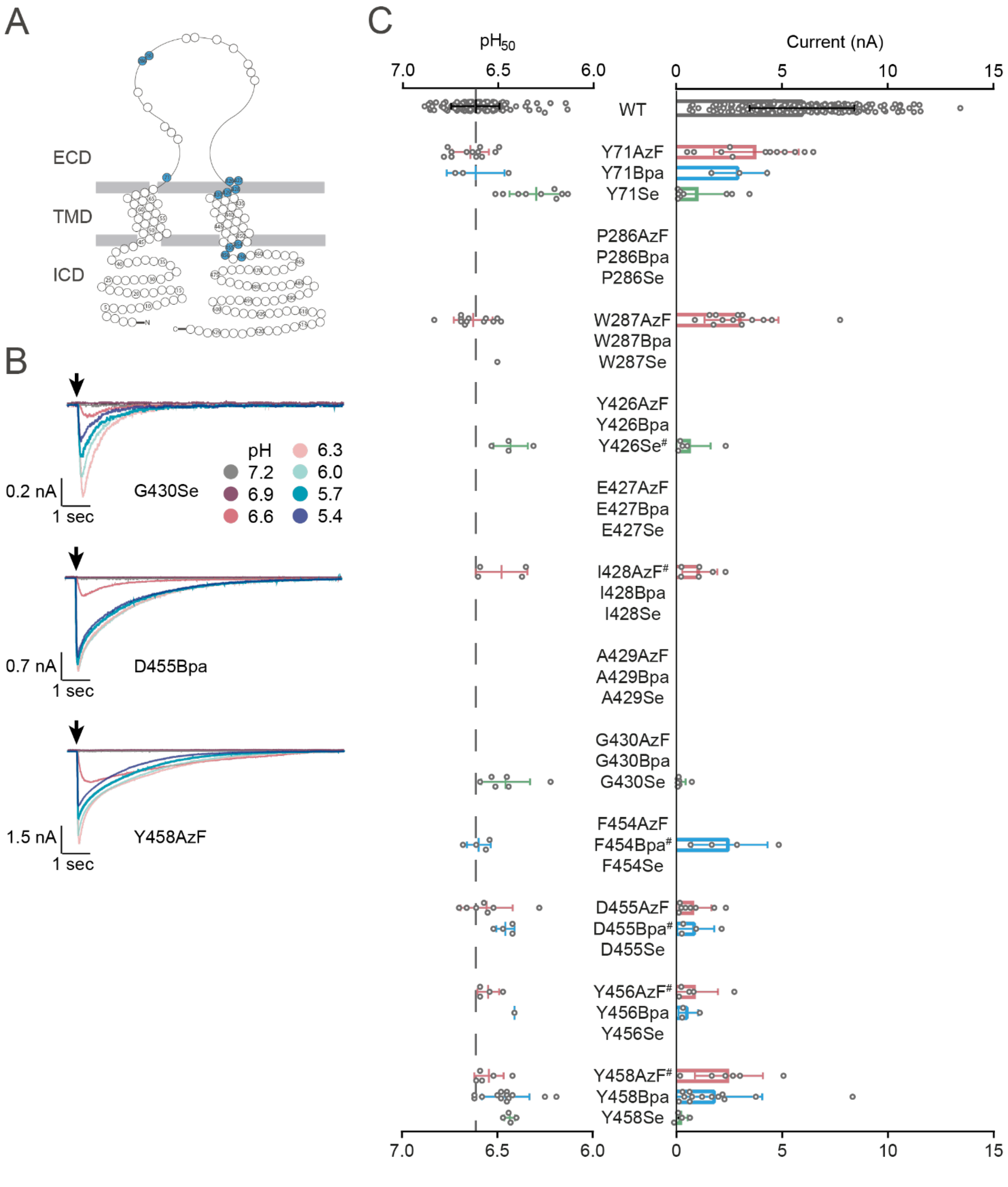

A
ECD
TMD
ICD
B
0.2 nA
1 sec
G430Se
pH
7.2
6.9
6.6
6.3
6.0
5.7
5.4
0.7 nA
1 sec
D455Bpa
1.5 nA
1 sec
Y458AzF
C
pH50
Current (nA)
7.0
6.5
6.0
0
5
10
15
WT
Y71AzF
Y71Bpa
Y71Se
P286AzF
P286Bpa
P286Se
W287AzF
W287Bpa
W287Se
Y426AzF
Y426Bpa
Y426Se#
E427AzF
E427Bpa
E427Se
I428AzF#
I428Bpa
I428Se
A429AzF
A429Bpa
A429Se
G430AzF
G430Bpa
G430Se
F454AzF
F454Bpa#
F454Se
D455AzF
D455Bpa#
D455Se
Y456AzF#
Y456Bpa
Y456Se
Y458AzF#
Y458Bpa
Y458Se

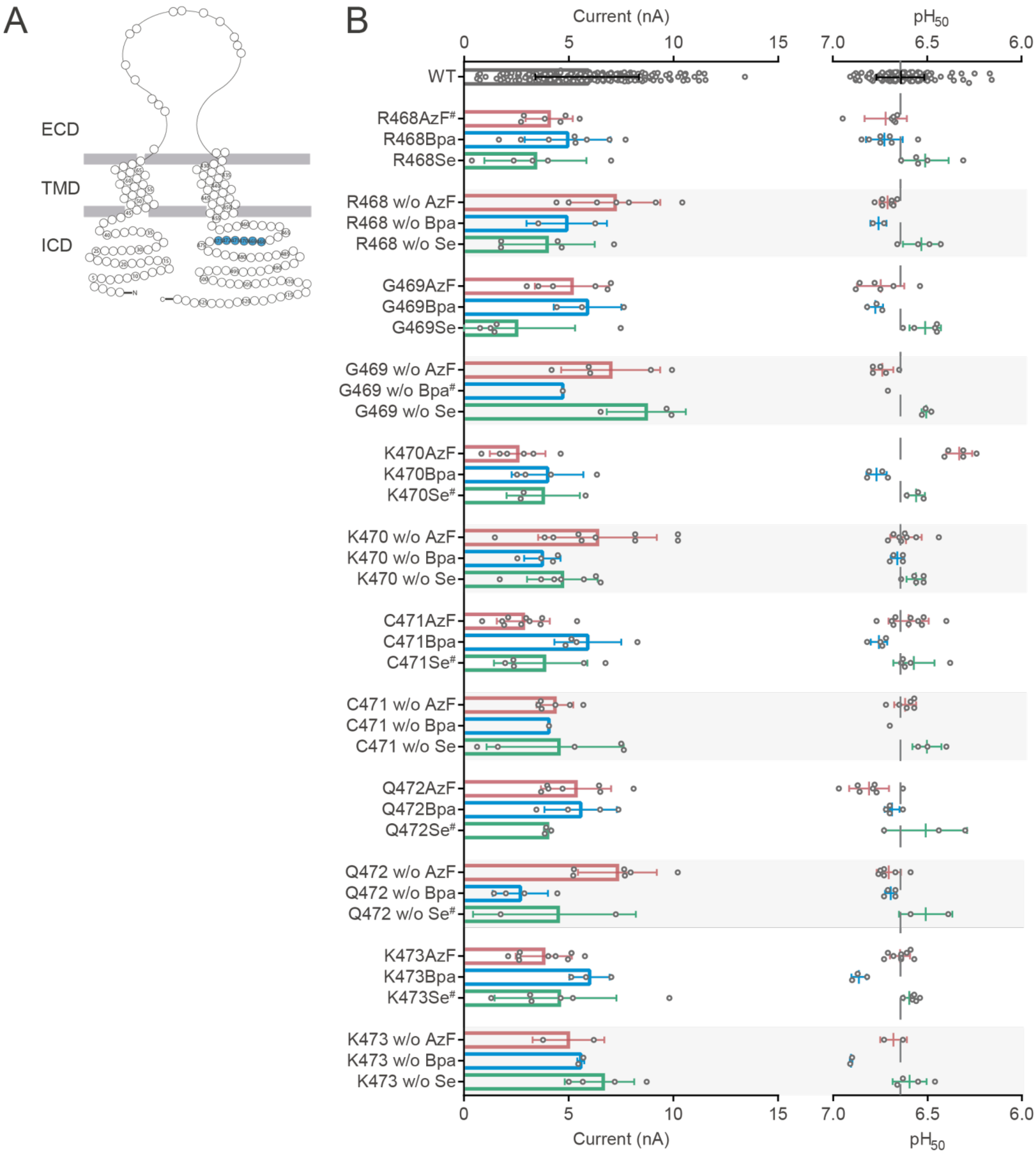

Figure S4 (including next 3 pages): Incorporation of ncAA photocrosslinkers into the hASIC1a C-terminus is unspecific from position 465 onwards. (A) Snake plots with tested positions marked in blue. (B) Dot plots comparing peak current sizes (left) and $pH_{50}$ (right), bars indicate mean ± S.D., ($^{\#}$) marks >20% tachyphylaxis (see also Table S1). For variants expressed in the absence of ncAAs that yielded currents, results are marked by underlying grey bars.

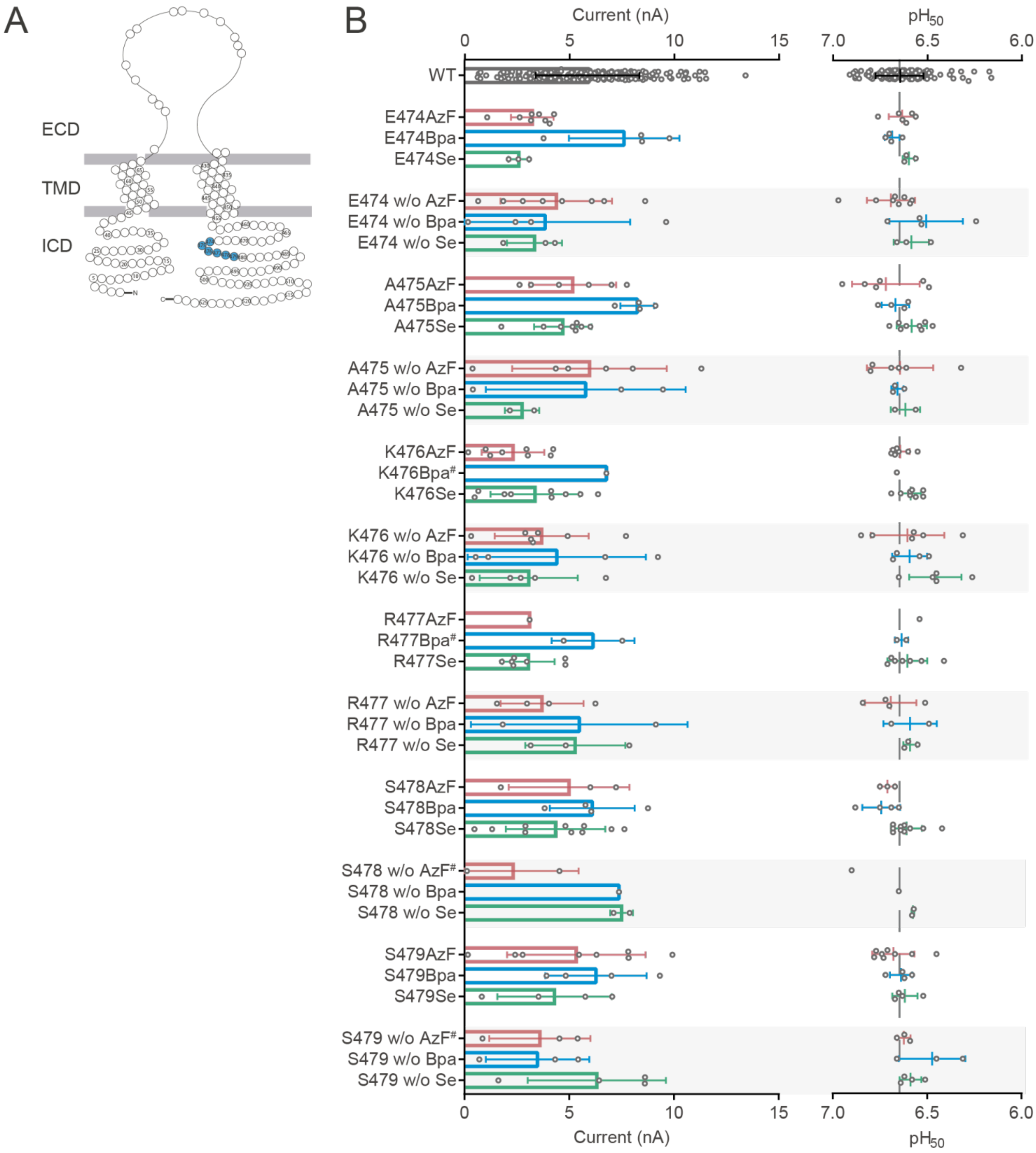
A
ECD
TMD
ICD
B
Current (nA)
0
5
10
15
pH50
7.0
6.5
6.0
WT
E474AzF
E474Bpa
E474Se
E474 w/o AzF
E474 w/o Bpa
E474 w/o Se
A475AzF
A475Bpa
A475Se
A475 w/o AzF
A475 w/o Bpa
A475 w/o Se
K476AzF
K476Bpa#
K476Se
K476 w/o AzF
K476 w/o Bpa
K476 w/o Se
R477AzF
R477Bpa#
R477Se
R477 w/o AzF
R477 w/o Bpa
R477 w/o Se
S478AzF
S478Bpa
S478Se
S478 w/o AzF#
S478 w/o Bpa
S478 w/o Se
S479AzF
S479Bpa
S479Se
S479 w/o AzF#
S479 w/o Bpa
S479 w/o Se

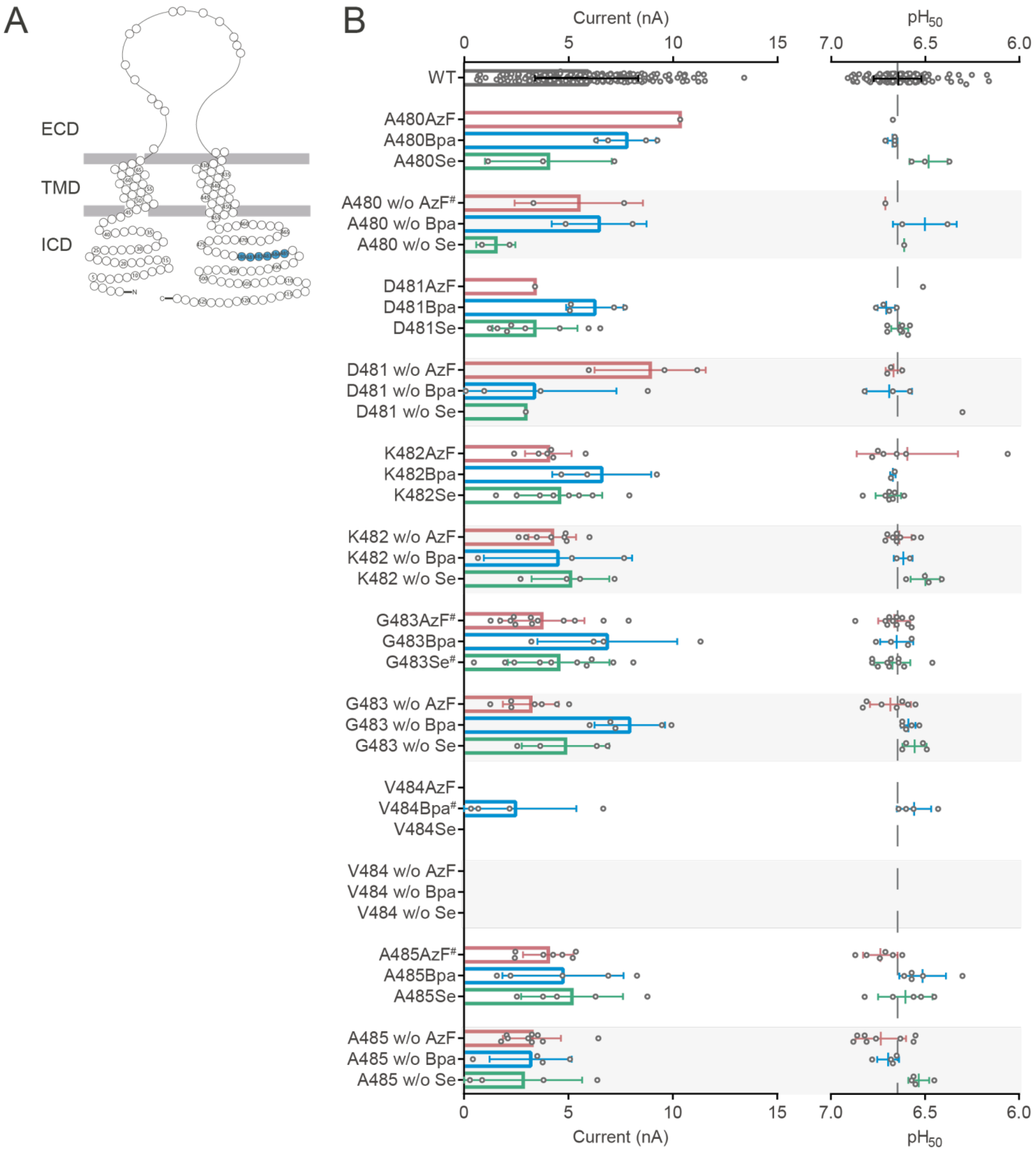
A
ECD
TMD
ICD
B
Current (nA)
pH50
WT
A480AzF
A480Bpa
A480Se
A480 w/o AzF#
A480 w/o Bpa
A480 w/o Se
D481AzF
D481Bpa
D481Se
D481 w/o AzF
D481 w/o Bpa
D481 w/o Se
K482AzF
K482Bpa
K482Se
K482 w/o AzF
K482 w/o Bpa
K482 w/o Se
G483AzF#
G483Bpa
G483Se#
G483 w/o AzF
G483 w/o Bpa
G483 w/o Se
V484AzF
V484Bpa#
V484Se
V484 w/o AzF
V484 w/o Bpa
V484 w/o Se
A485AzF#
A485Bpa
A485Se
A485 w/o AzF
A485 w/o Bpa
A485 w/o Se

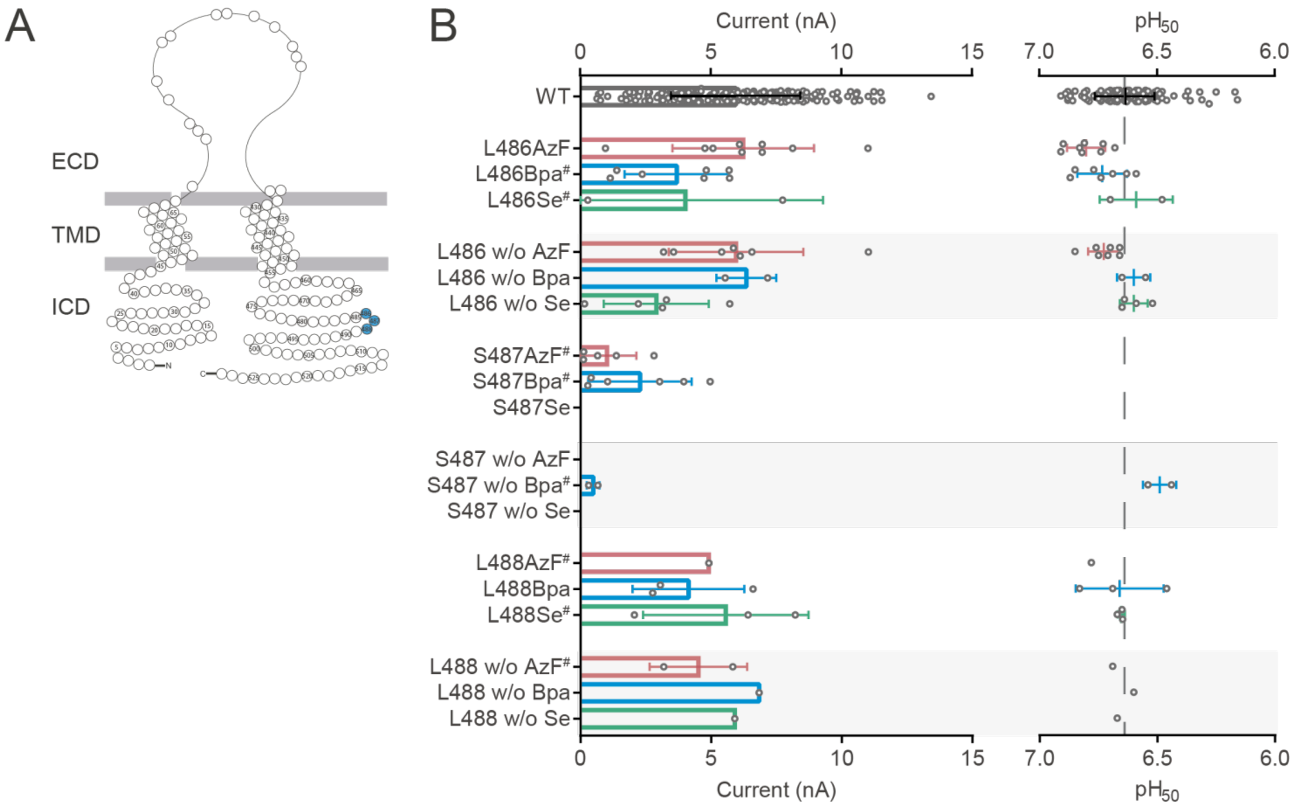

Figure S5 (next page): hASIC1a C-terminal positions distal of L465 are not essential for proton-gated current responses. (A) Snake plot of hASIC1a highlighting C-terminal positions in blue. (B) Representative current traces of C466TAG with and without Bpa (upper panels) and R467TAG with and without Se-AbK (termed Se, lower panels) as recorded on the SyncroPatch 384PE. (C) Dot plots comparing $pH_{50}$ (left) and peak current sizes (right), bars indicate mean ± S.D., (#) marks >20% tachyphylaxis (see also Table S1). For variants expressed in the absence of ncAAs that yielded currents, results are marked by underlying grey bars. (D) Concentration response curves of hASIC1a WT (black) and C-terminally truncated constructs recorded in HEK 293T cells (APC, left panel) and *X. laevis* oocytes (TEVC, right panel). (E) Western blot using an ASIC1a-antibody targeting an extracellular epitope documents truncation of the protein.

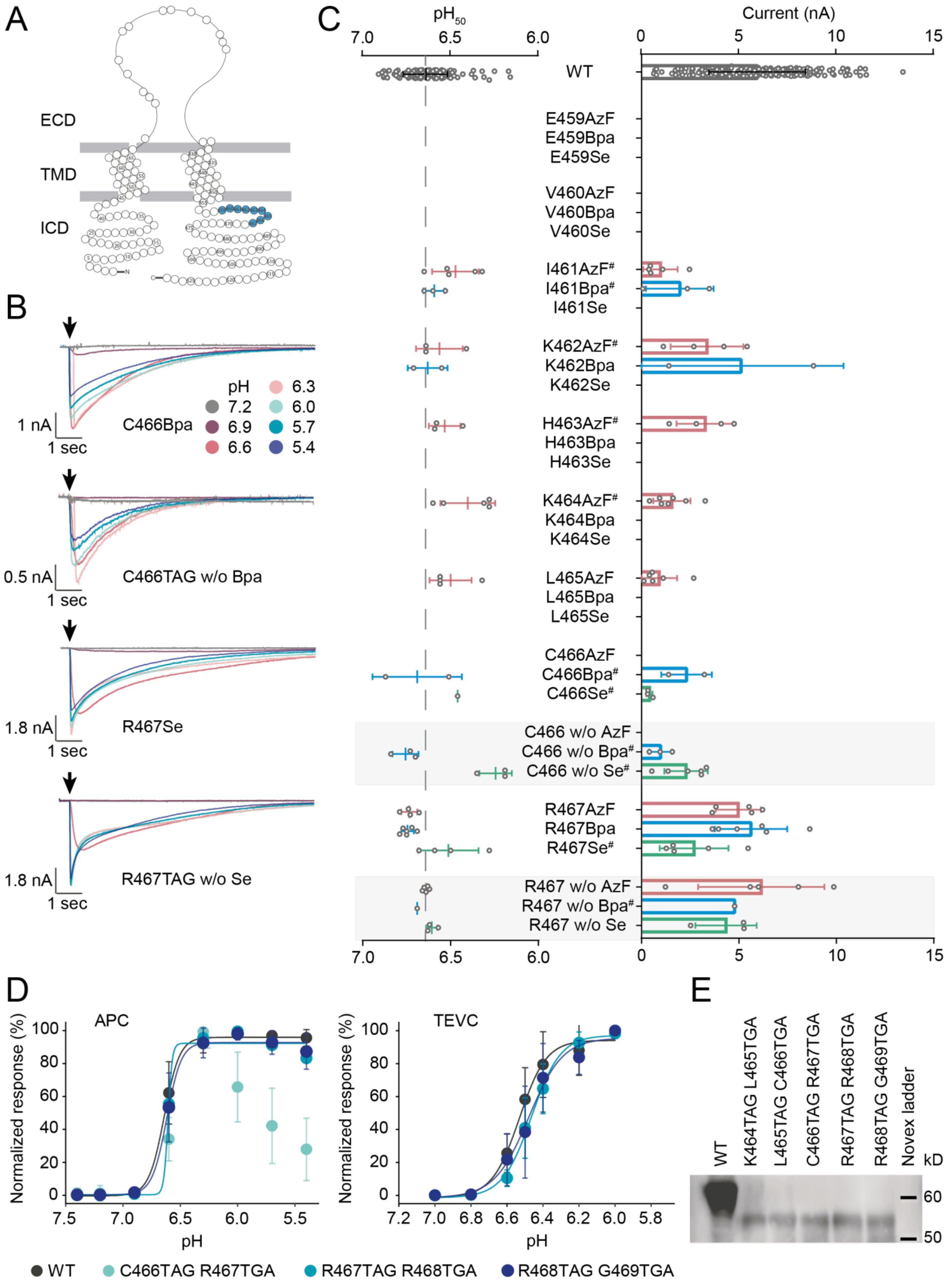
A
ECD
TMD
ICD
B
C466Bpa
pH
7.2
6.9
6.6
6.3
6.0
5.7
5.4
1 nA
1 sec
C466TAG w/o Bpa
0.5 nA
1 sec
R467Se
1.8 nA
1 sec
R467TAG w/o Se
1.8 nA
1 sec
C
pH50
Current (nA)
WT
E459AzF
E459Bpa
E459Se
V460AzF
V460Bpa
V460Se
I461AzF#
I461Bpa#
I461Se
K462AzF#
K462Bpa
K462Se
H463AzF#
H463Bpa
H463Se
K464AzF#
K464Bpa
K464Se
L465AzF
L465Bpa
L465Se
C466AzF
C466Bpa#
C466Se#
C466 w/o AzF
C466 w/o Bpa#
C466 w/o Se#
R467AzF
R467Bpa
R467Se#
R467 w/o AzF
R467 w/o Bpa#
R467 w/o Se
D
APC
TEVC
Normalized response (%)
pH
WT
C466TAG R467TGA
R467TAG R468TGA
R468TAG G469TGA
E
WT
K464TAG L465TGA
L465TAG C466TGA
C466TAG R467TGA
R467TAG R468TGA
R468TAG G469TGA
Novex ladder
kD
60
50

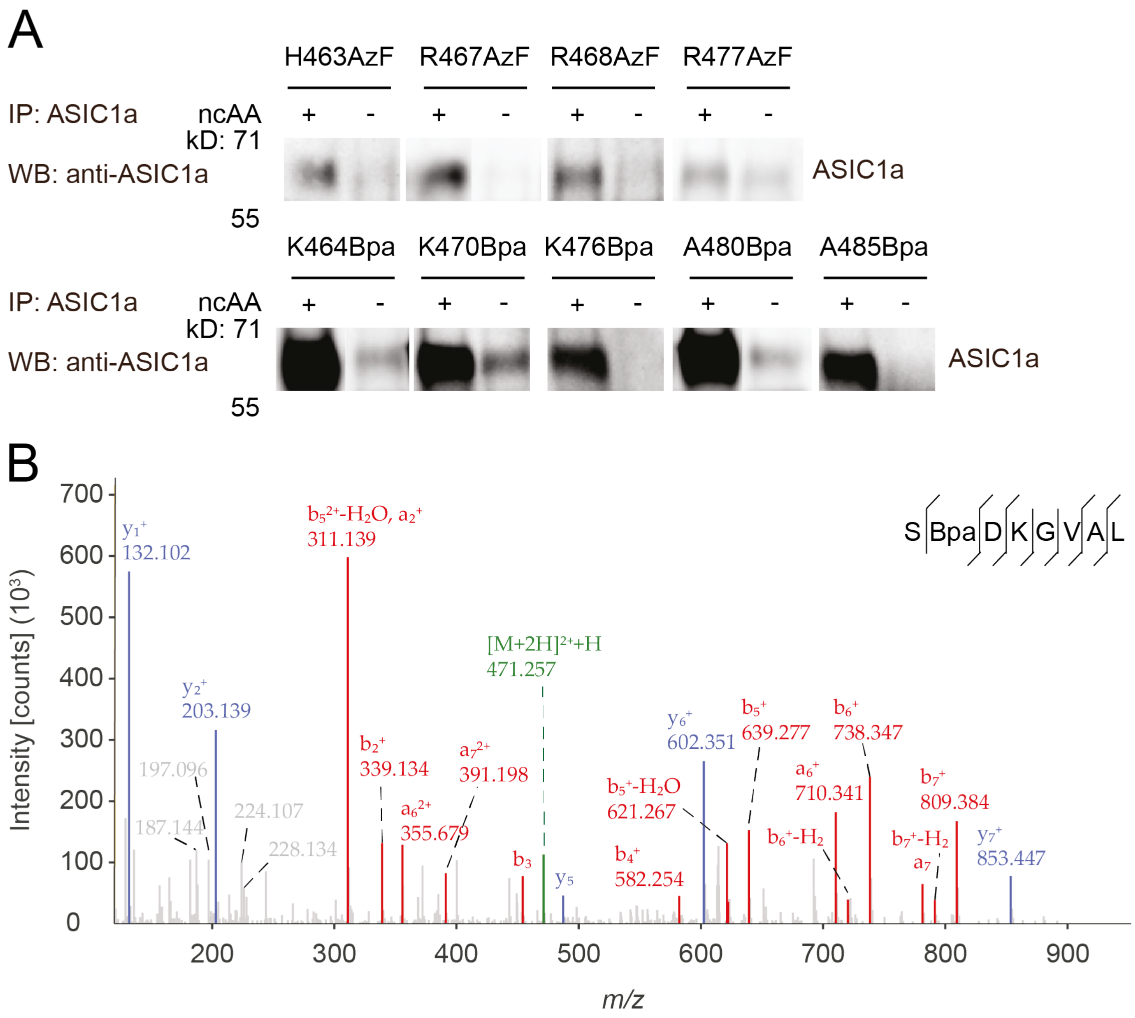

Figure S6: (A) Incorporation of AzF and Bpa in the C-terminus is efficient with position-dependent specificity. Selected hASIC1a TAG variants were expressed in presence or absence of 10 µM AzF-ME or 1 mM Bpa in HEK 293T ASIC1a-KO cells for 48 hrs, the full-length protein was purified via a C-terminal 1D4-tag and visualized by western blotting using the indicated antibody (AB). Only small amounts of full-length protein were detected in the absence of ncAA, indicating efficient incorporation. (B) Mass spectrometry confirms incorporation of Bpa at position 480. HCD fragment ion mass spectrum of the precursor peptide SBpaDKGVAL (positions 479-486, green) and the corresponding fragment ions (a- and b-ion series red, y ion series blue). Theoretical peptide mass 940.478, experimental *m/z* 470.743 (+0.53 ppm), charge +2.

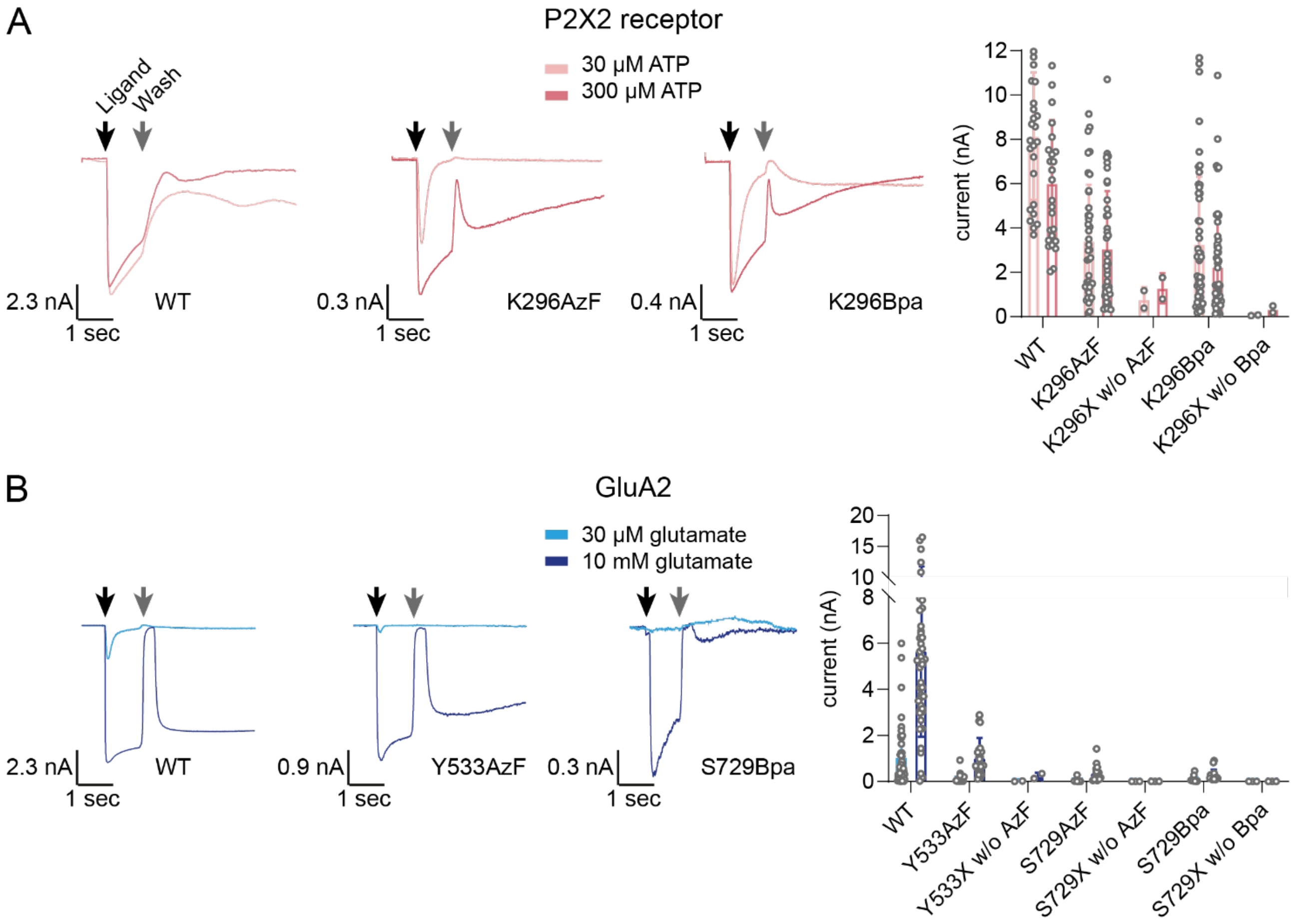

Figure S7: The established FACS APC workflow is suitable to assess ncAA incorporation into other ligand-gated ion channels. Example current traces and dot plots comparing current sizes for WT and ncAA-containing variants of the P2X2 receptor (A) and GluA2 (B) at different concentrations of ATP or glutamate, respectively. Cells expressing GluA2 were incubated with 100 µM cyclothiazide (0.8% v/v DMSO) for one minute before glutamate addition to reduce rapid desensitization [3]. Black arrow indicates ligand application, grey arrow indicates addition of wash solution. As apparent from the current traces, application of the wash solution removes the ligand from the channels temporarily, but as it is still present in the well, channels can re-open and desensitize over time. In order to enable concentration-response curve measurements, all ligand has to be removed from the well. Bar graphs are mean ± S.D., values shown in Table S2.

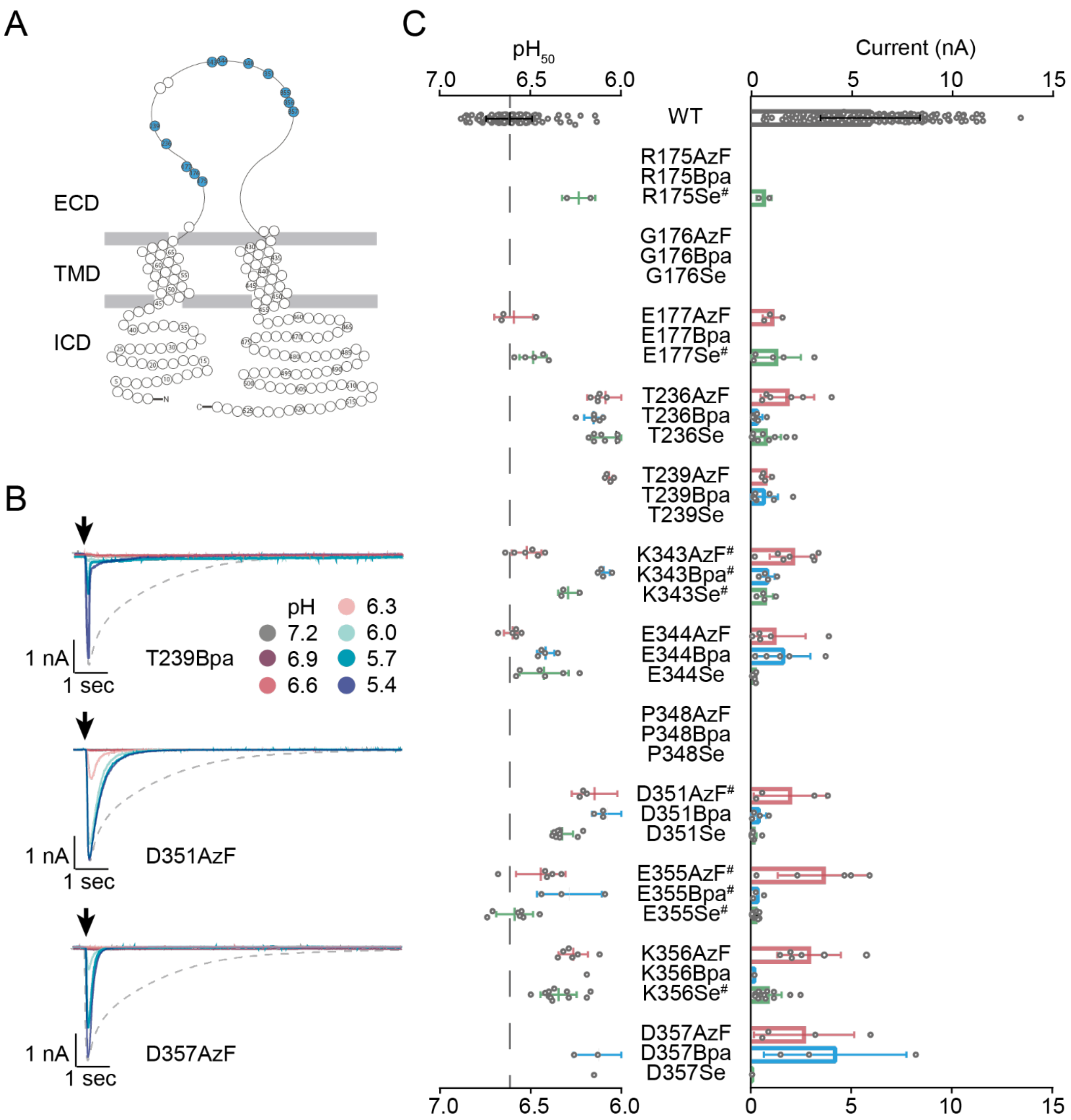

Figure S8: Evaluation of twelve positions around the PcTx1 binding site reveals several channel variants with lowered proton sensitivity and accelerated current decay. (A) Snake plot of hASIC1a highlighting assessed positions in blue. (B) Representative current traces of T239Bpa, D351AzF and D357AzF as recorded on the SyncroPatch 384PE, with arrows indicating time of proton application. Dashed lines indicate WT current in response to pH 6.0 application. (C) Dot plots comparing $pH_{50}$ (left) and peak current sizes (right), bars indicate mean ± S.D., (#) marks >20% tachyphylaxis (see also Table S1).

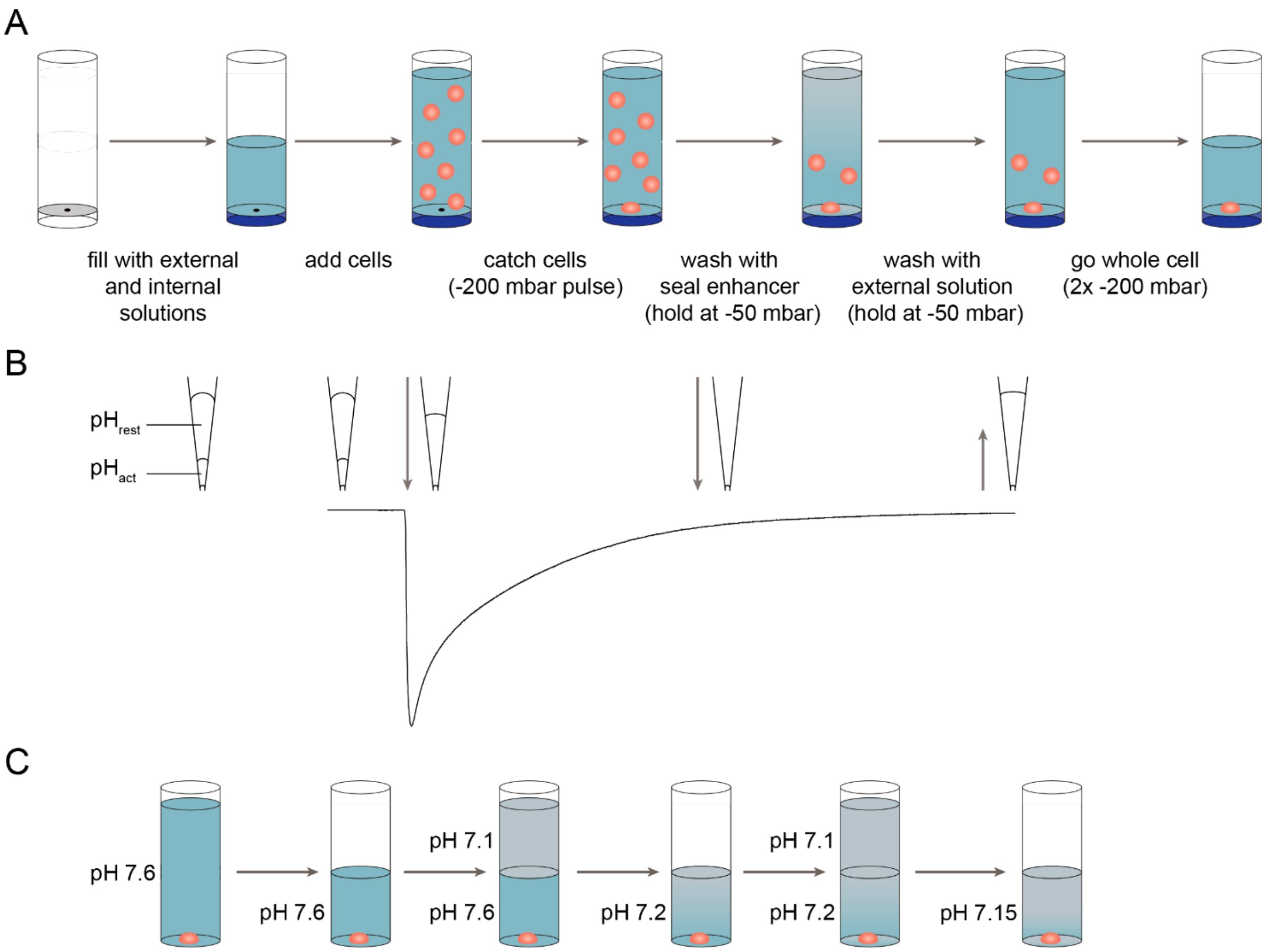

Figure S9: Schematic representations of the patching process, stacked ligand addition and exchange of the conditioning pH on the SyncroPatch 384PE. (A) After filling the wells with external and internal solution (light and dark blue), cells (orange) are added and caught on the hole by brief application of -200 mbar pressure. The cells are held in place with -50 mbar during the wash steps with seal enhancer and external solution before going into whole cell configuration via two pulses at -200 mbar. (B) For the stacked ligand application, pipettes are filled with 45 µl of resting pH ($pH_{rest}$) followed by 5 µl solution of activating pH ($pH_{act}$). Dispension of $pH_{act}$ leads to channel activation and desensitization in the presence of ligand, followed by dispension of $pH_{rest}$ with a delay of 5 sec to wash out the ligand. Solution is slowly taken back up into the pipette at the end of each sweep, followed by a wash step with $pH_{rest}$ (not shown). (C) When measuring SSD curves, the open-well system of the SyncroPatch 384PE requires repeated mixing steps to approximate the target conditioning pH without disturbing the cell (orange). 50% of the liquid (blue) are aspirated and replaced by lower pH solution (grey) twice to obtain the final conditioning pH, here pH 7.15.

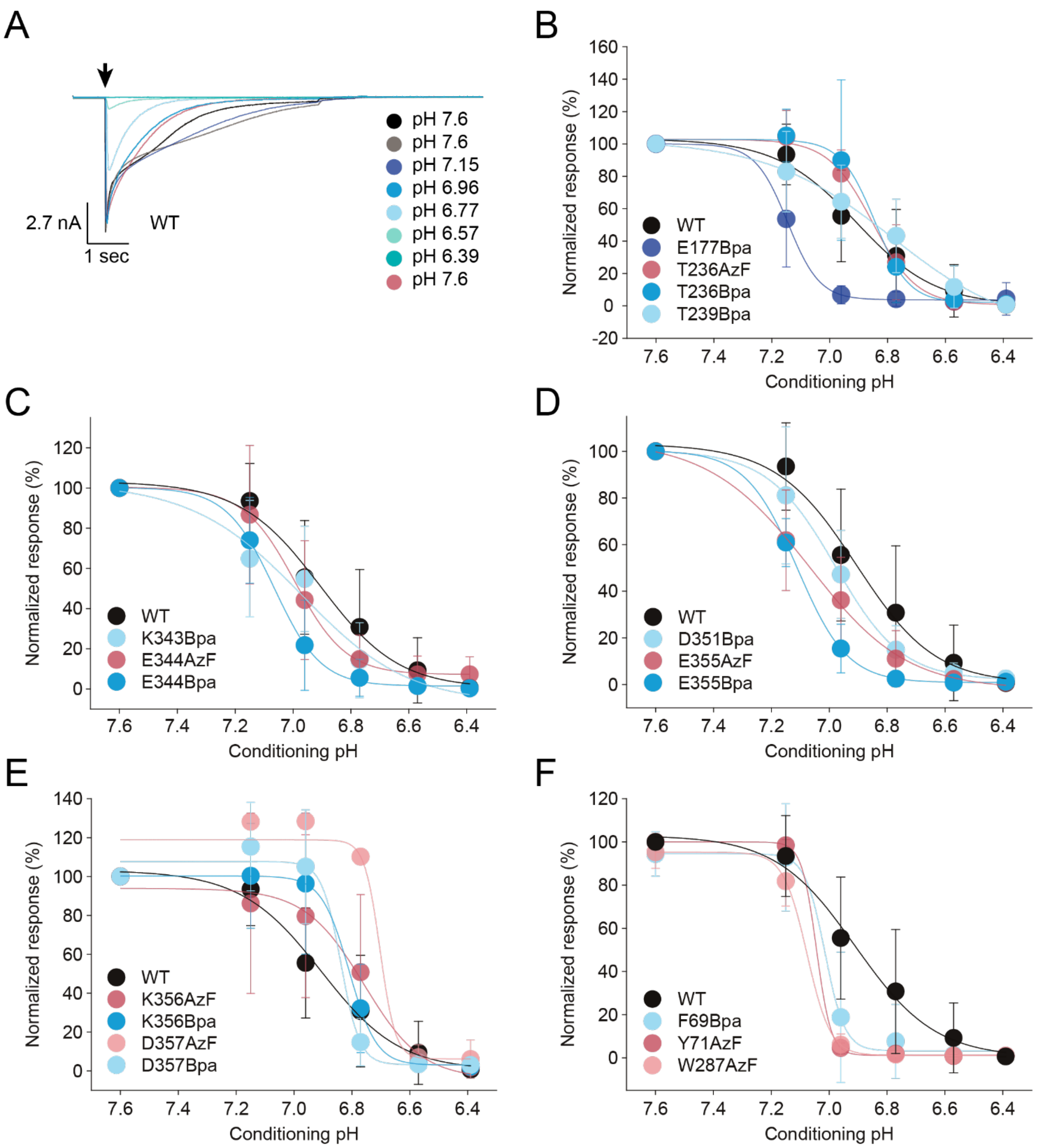

Figure S10: Steady-state desensitization of hASIC1a WT and 17 variants containing AzF or Bpa in the acidic pocket or interface region can be efficiently assessed using APC. (A) Example current trace for hASIC1a WT. Currents were evoked by application of pH 5.6 after conditioning at the indicated pH values for 2 min. Current recovery was assessed at the end of the protocol and cells that did not regain current were excluded from the analysis. (B-F) SSD curves of WT, 14 variants carrying AzF or Bpa in the acidic pocket, and three control positions in the interface region. Currents were normalized to the mean of the first two applications. See Table S4 for $pH_{50}$ SSD values, nH and n.

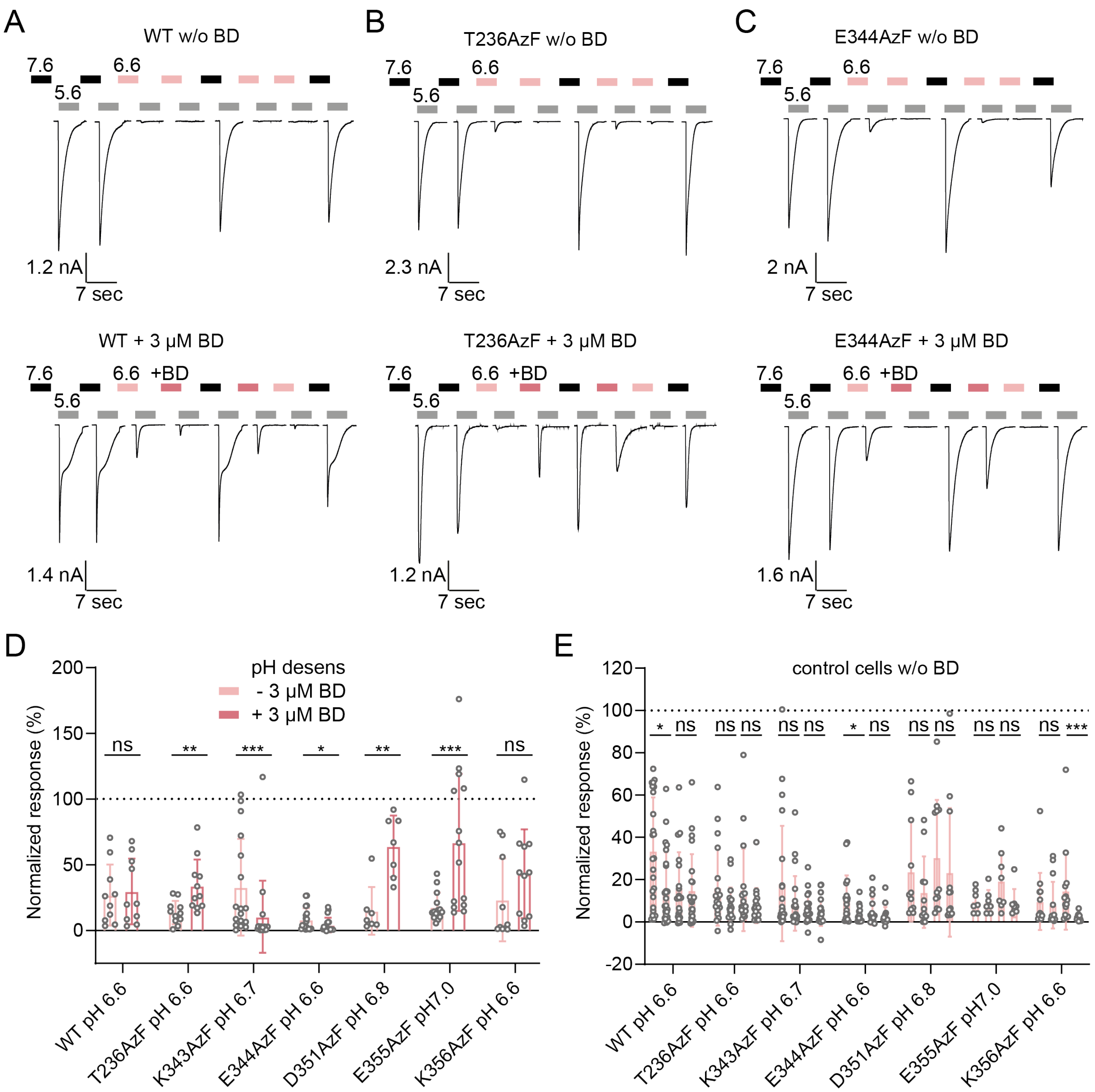

Figure S11: BigDyn modulation of hASIC1a WT and six variants carrying AzF in the acidic pocket. (A-C) Characteristic current traces of the full APC protocols for WT (A), T236AzF (B) and E344AzF (C) with and without 3 µM BigDyn (lower vs. upper panel). Cells were first exposed to two activation pulses with pH 5.6 (grey bars, 5 sec) after conditioning at pH 7.6 (black bars) to determine the control current, followed by two rounds of activation after 2 min conditioning with a pH that induces SSD (light pink bars) and a control pulse to evaluate current recovery. For half of the cell population, 3 µM BigDyn (dark pink bars) were co-applied during the second conditioning period to measure rescue from SSD. This assessment of SSD and recovery was repeated with peptide co-application during the first SSD-conditioning to also evaluate peptide wash out. Currents were normalized to the average of the first two control pulses to compare modulation at different conditions (values in Table S5, pink and black bars not to scale). (D) Bar graph comparing current after SSD in absence and presence of 3 µM BigDyn (traces 3+4 in A-C, lower panel). (E) Bar graph comparing current after

SSD for control cells not exposed to BigDyn (traces 3+4 and 6+7 in A-C, upper panel). Bar graphs show mean ± S.D, dashed line indicates 100%, values in Table S5. (*) denotes significant difference between groups, $p < 0.05$; (**): $p < 0.01$; (***): $p < 0.001$; ns: not significant; Mann Whitney test.

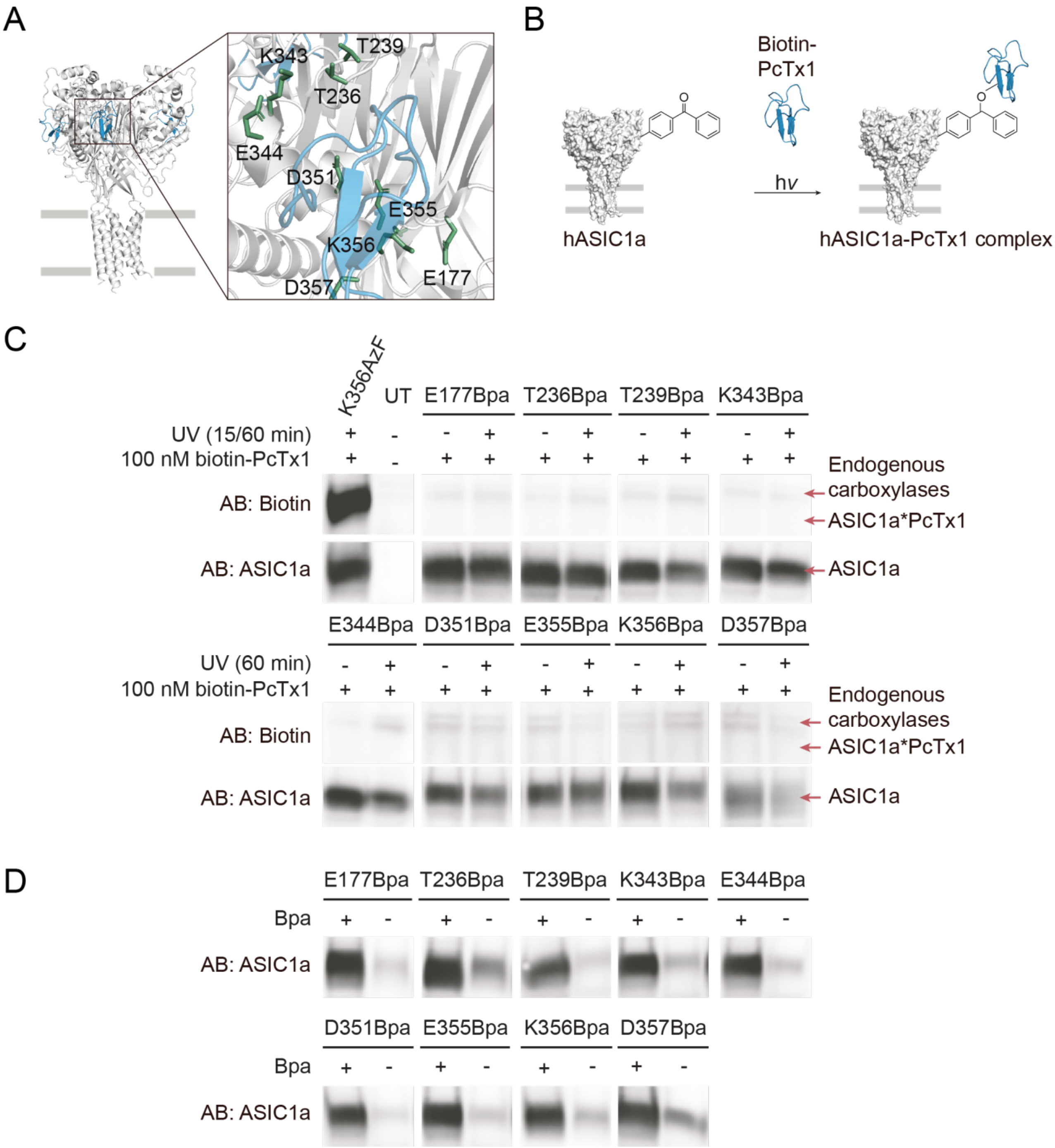

Figure S12: PcTx1 is not detected in the ASIC1a acidic pocket when Bpa is used for photocrosslinking. (A) Structure of chicken ASIC1 (white) in complex with PcTx1 (blue, PDB: 4FZ0), inset shows individual side chains replaced by Bpa in the acidic pocket (green), none of which crosslinked to biotin-PcTx1. (B) Schematic workflow for Bpa crosslinking to biotin-PcTx1 (see also Figure 5). (C) Western blot of purified hASIC1a K356AzF, untransfected cells (UT) and variants carrying Bpa in the extracellular domain detected using the specified antibodies (AB). Biotin-PcTx1 is only detected in the control sample containing AzF at position 356 (15 min UV exposure), but not in any of the nine positions containing Bpa (60 min UV exposure, positions coloured green in A). The detected double band by the anti-biotin AB originates from endogenous biotin-dependent carboxylases [4, 5]. (D) Control experiments demonstrating efficient Bpa incorporation at all positions tested for crosslinking in C. Stop-codon containing hASIC1a mutants were grown in the presence or absence of 1 mM Bpa in HEK 293T cells for 48 hours, after which the resulting full-length protein was purified via a C-terminal 1D4-tag and visualized by western blotting using the indicated antibody (AB). With the exception of positions 236 and 357, only small amounts of full-length protein were detected in absence of Bpa (compared to those obtained in its presence), demonstrating efficient incorporation.

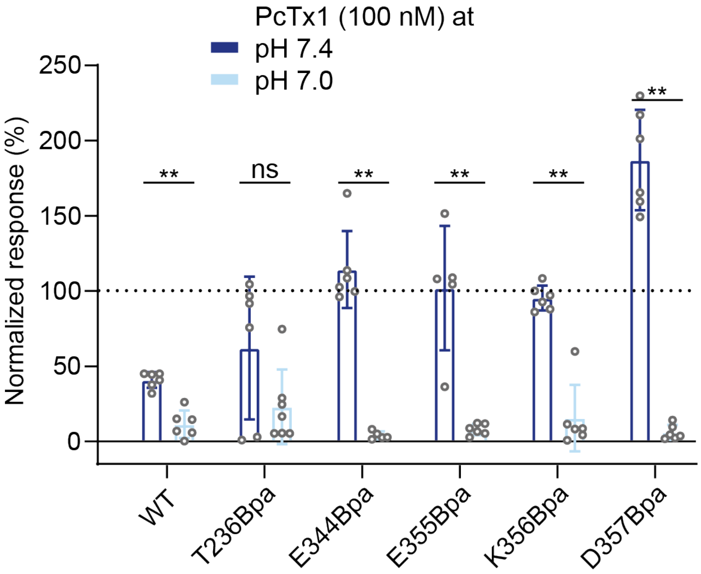

Figure S13: Bar graph for PcTx1 modulation of hASIC1a WT and selected variants containing Bpa in the acidic pocket at different pH. Cells were incubated with 100 nM PcTx1 at varying conditioning pH for 2 min before activation at pH 5.6 and the current was normalized to the average of the four preceding and following control currents after conditioning at pH 7.4. Bar graph shows mean ± S.D, dashed line indicates 100%, values shown in Table S6. (**) denotes significant difference between groups, $p < 0.01$; ns: not significant; Mann Whitney test.

Figure S14 (next two pages): Original Western blots for AzF crosslinking. Black bars indicate protein ladders for clarity (left panel), original markers (coomassie) are overlayed with the blot (chemiluminescence, right panel). Areas cropped for Figure 5 are marked with boxes. (A) Western blot for positions 236, 239, 343, 356 and 357. (B) Western blot for positions 177, 239 and 344. (C) Western blot for positions 71, 287, 69, 80, 253, 413, 351 and 355. (D) Western blot for the F352L K356AzF double mutant. Data is representative of 2-3 individual experiments. Control experiments demonstrating efficient AzF incorporation for all above positions are published in [6].

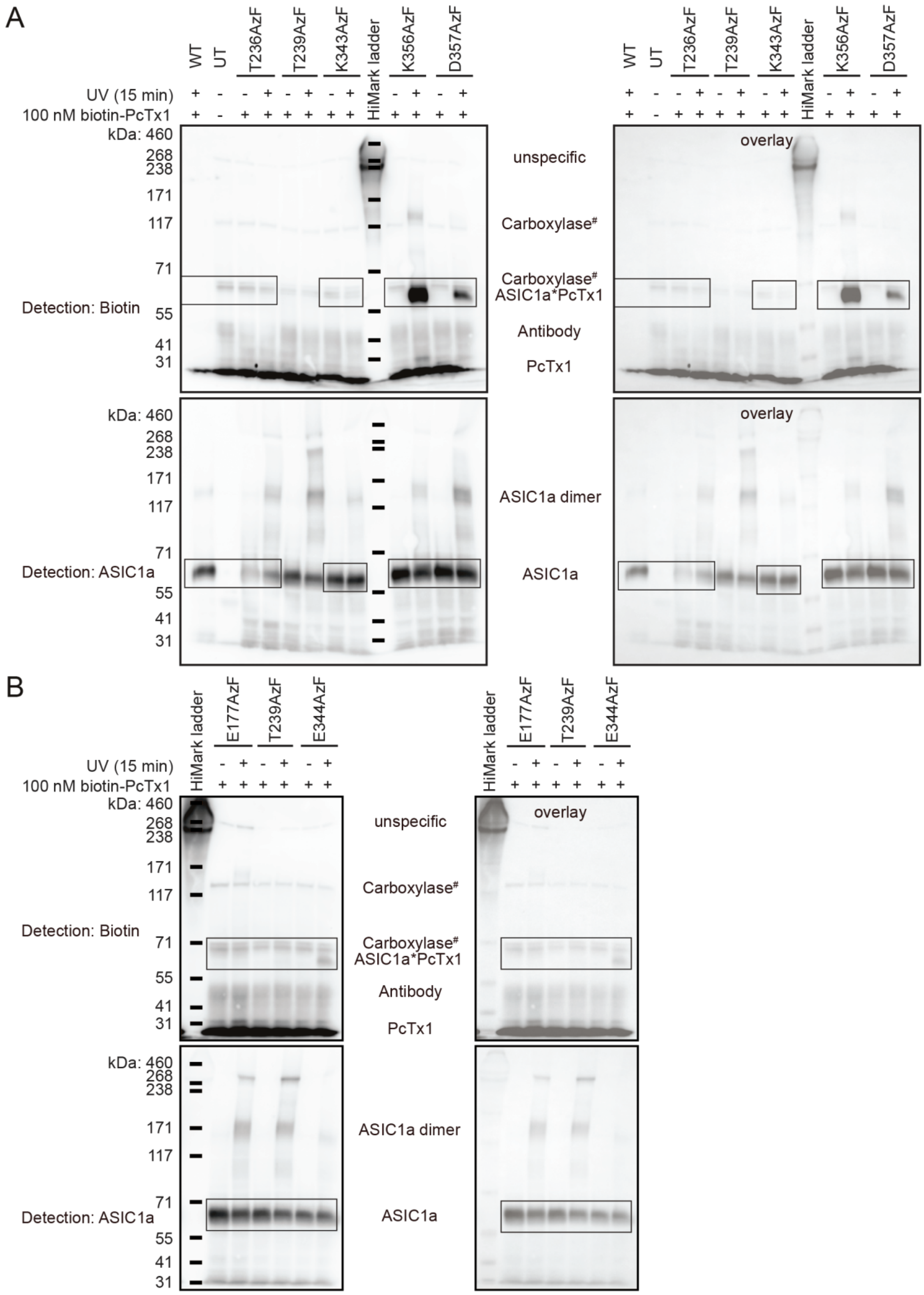
A
WT
UT
T236AzF
T239AzF
K343AzF
HiMark ladder
K356AzF
D357AzF
UV (15 min)
100 nM biotin-PcTx1
kDa: 460
268
238
171
117
71
55
41
31
Detection: Biotin
Detection: ASIC1a
unspecific
Carboxylase#
Carboxylase#
ASIC1a*PcTx1
Antibody
PcTx1
ASIC1a dimer
ASIC1a
overlay
B
E177AzF
T239AzF
E344AzF

C

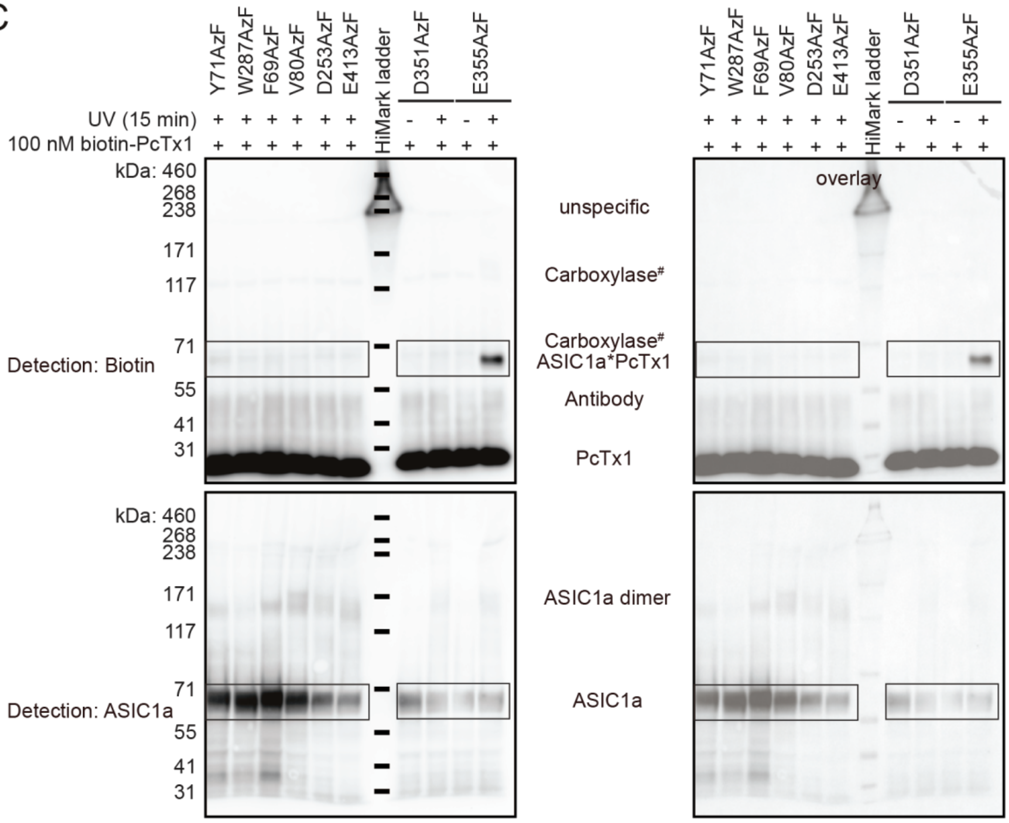
Y71AzF
W287AzF
F69AzF
V80AzF
D253AzF
E413AzF
HiMark ladder
D351AzF
E355AzF
UV (15 min)
100 nM biotin-PcTx1
kDa: 460
268
238
171
117
71
55
41
31
Detection: Biotin
Detection: ASIC1a
unspecific
Carboxylase#
Carboxylase#
ASIC1a*PcTx1
Antibody
PcTx1
ASIC1a dimer
ASIC1a
overlay

D

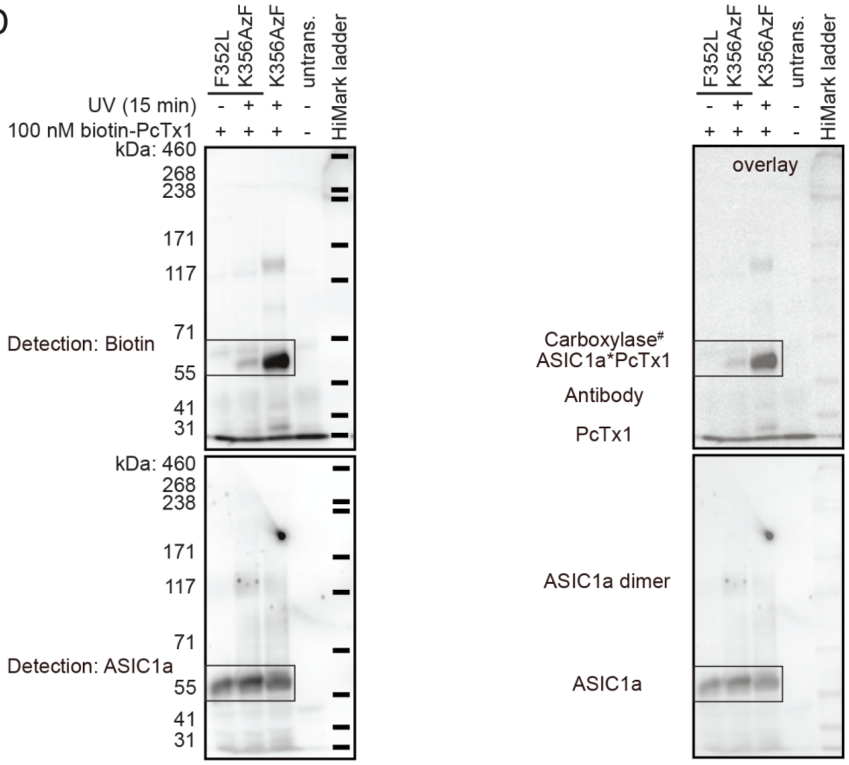
F352L
K356AzF
K356AzF
untrans.
HiMark ladder
UV (15 min)
100 nM biotin-PcTx1
kDa: 460
268
238
171
117
71
55
41
31
Detection: Biotin
Detection: ASIC1a
Carboxylase#
ASIC1a*PcTx1
Antibody
PcTx1
ASIC1a dimer
ASIC1a
overlay

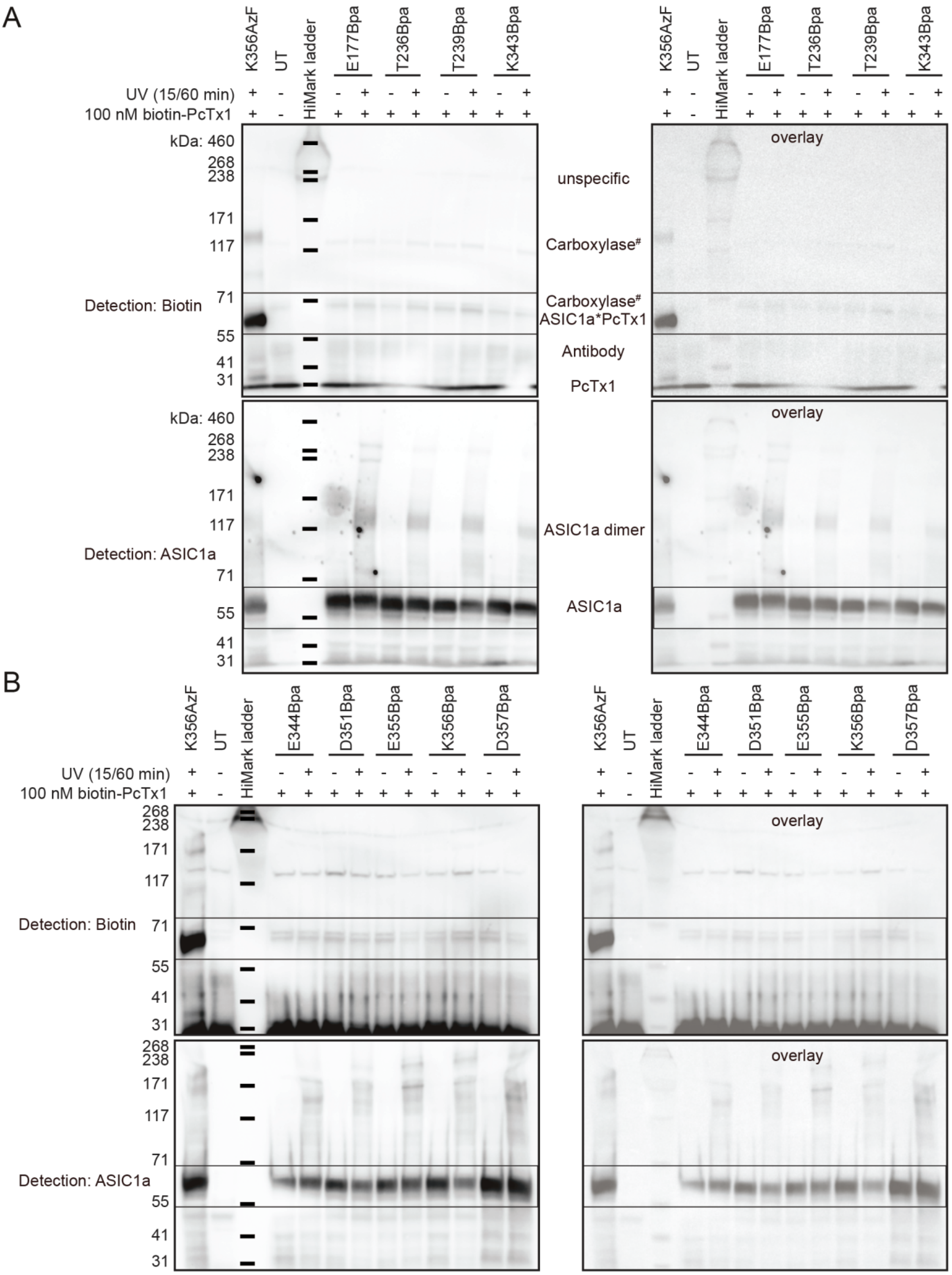
A
K356AzF
UT
HiMark ladder
E177Bpa
T236Bpa
T239Bpa
K343Bpa
UV (15/60 min)
100 nM biotin-PcTx1
kDa: 460
268
238
171
117
71
55
41
31
Detection: Biotin
Detection: ASIC1a
overlay
unspecific
Carboxylase#
Carboxylase#
ASIC1a*PcTx1
Antibody
PcTx1
ASIC1a dimer
ASIC1a
B
K356AzF
UT
HiMark ladder
E344Bpa
D351Bpa
E355Bpa
K356Bpa
D357Bpa
UV (15/60 min)
100 nM biotin-PcTx1
268
238
171
117
71
55
41
31
Detection: Biotin
Detection: ASIC1a
overlay

C

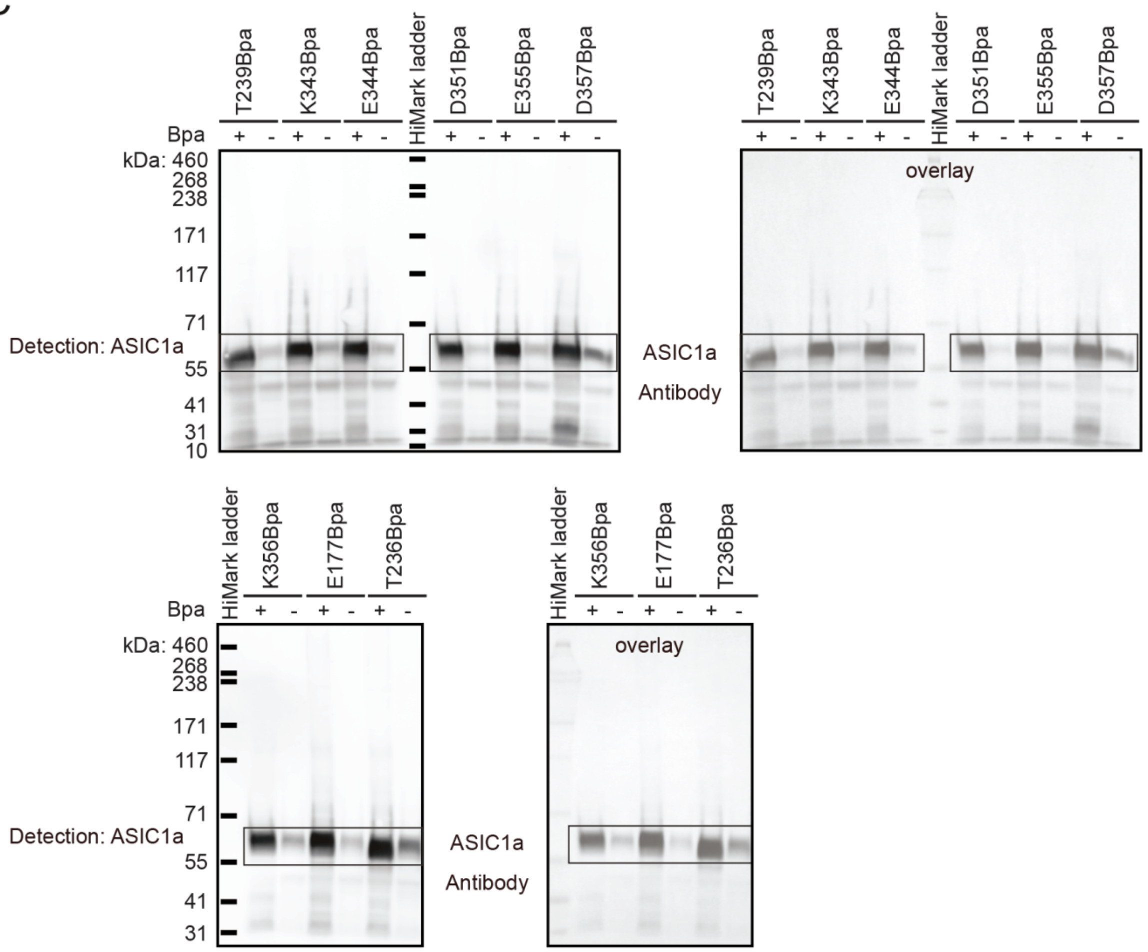

Figure S15: Original Western blots for Bpa crosslinking. Black bars indicate protein ladders for clarity (left panel), original markers (coomassie) are overlayed with the blot (chemiluminescence, right panel). Areas cropped for Figure S12 are marked with boxes. (A) Western blot for positions 177, 236, 239 and 343, including K356AzF as a positive control. (B) Western blot for positions 344, 351, 355, 356 and 357, including K356AzF as a positive control. (C) Western blot demonstrating efficient Bpa incorporation at positions 239, 343, 344, 351, 355 and 357 (upper panel) and at positions 177, 236 and 356 (lower panel). Data is representative of 2-3 individual experiments.

**Supplementary tables S1-6**

Table S1 (next 15 pages): Electrophysiological characterization of the hASIC1a TAG variant library as assessed on the SyncroPatch 384PE. Values for $pH_{50}$, $n_H$ and Imax are shown as mean ± S.D. for n ≥ 3 and as averages for n=1–2. ($^{\#}$) indicates pronounced tachyphylaxis (final sweep <80% of normalized peak current). Average transfection efficiency (TE) is shown in (%) as measured via GFP fluorescence on the FACS instrument.

| Position replaced by TAG | ncAA | mean $pH_{50}$ ± S.D. (n) | mean $n_H$ ± S.D. (n) | mean Imax (nA) ± S.D. (n) | average TE (%) |
|---|---|---|---|---|---|
| E6 | AzF# | 6.69 ± 0.07 (8) | 8.66 ± 2.76 (8) | 3.82 ± 1.65 (10) | 12.2 |
| | AzF neg. | n.a. | n.a. | 0 (0) | n.a. |
| | Bpa | 6.55 (2) | 8.76 (2) | 2.87 (2) | 6.8 |
| | Bpa neg. | 6.44 (2) | 8.42 (2) | 1.14 (2) | n.a. |
| | Se-AbK# | 6.57 ± 0.09 (6) | 7.66 ± 3.31 (3) | 1.8 ± 2.2 (8) | 5.8 |
| | Se-AbK neg. | n.a. | n.a. | 0 (0) | n.a. |
| E7 | AzF# | 6.58 ± 0.08 (7) | 7.78 ± 6.04 (6) | 2.47 ± 1.56 (9) | 25.2 |
| | AzF neg. | n.a. | n.a. | 1.01 (1) | n.a. |
| | Bpa# | 6.54 ± 0.07 (6) | 6.8 ± 3.45 (6) | 1.12 ± 1.1 (7) | 12.7 |
| | Bpa neg. # | 6.45 (1) | 7.81 (1) | 0.12 (2) | n.a. |
| | Se-AbK# | 6.6 ± 0.04 (5) | 7.54 (2) | 2.41 ± 2.58 (5) | 5 |
| | Se-AbK neg. | 6.59 (2) | 32.34 (2) | 0.99 (2) | n.a. |
| E8 | AzF | 6.51 ± 0.08 (4) | 6.29 ± 0.98 (3) | 4.7 ± 4.53 (4) | 13.5 |
| | AzF neg. | n.a. | n.a. | 0 (0) | n.a. |
| | Bpa | 6.44 ± 0.07 (10) | 5.61 ± 1.4 (10) | 1.86 ± 1.12 (10) | 10.5 |
| | Bpa neg. | 6.44 (1) | 6.13 (1) | 0.23 (1) | n.a. |
| | Se-AbK# | 6.55 ± 0.06 (4) | 9.3 ± 2.55 (4) | 2.17 ± 2.22 (4) | 3.9 |
| | Se-AbK neg. | n.a. | n.a. | 0 (0) | n.a. |
| E9 | AzF | 6.56 ± 0.06 (10) | 6.66 ± 4.37 (8) | 2.33 ± 1.37 (10) | 13.3 |
| | AzF neg. | n.a. | n.a. | 0 (0) | n.a. |
| | Bpa# | 6.59 ± 0.04 (10) | 7.80 ± 1.78 (7) | 1.99 ± 1.96 (11) | 13.8 |
| | Bpa neg. | 6.43 ± 0.16 (3) | 10.8 ± 3.09 (3) | 1.57 ± 1.13 (3) | n.a. |
| | Se-AbK# | 6.60 ± 0.04 (7) | 7.32 ± 2.06 (4) | 0.42 ± 0.23 (8) | 6 |
| | Se-AbK neg.# | 6.55 ± 0.06 (3) | 7.25 (2) | 0.85 ± 0.4 (3) | n.a. |
| V10 | AzF# | 6.54 ± 0.07 (12) | 6.66 ± 2.98 (7) | 1.56 ± 1.12 (13) | 6.1 |
| | AzF neg. | n.a. | n.a. | 0 (0) | n.a. |
| | Bpa | n.a. | n.a. | 0 (0) | 20.7 |
| | Bpa neg. | n.a. | n.a. | 0 (0) | n.a. |
| | Se-AbK# | 6.58 ± 0.01 (3) | 9.94 (1) | 0.91 ± 0.89 (4) | 7.7 |
| | Se-AbK neg. | n.a. | n.a. | 0 (0) | n.a. |
| G11 | AzF# | 6.51 ± 0.06 (6) | 8.0 ± 2.56 (5) | 0.72 ± 0.22 (7) | 6.9 |
| | AzF neg. | n.a. | n.a. | 0 (0) | n.a. |
| | Bpa | 6.44 ± 0.12 (9) | 6.21 ± 1.96 (9) | 2.35 ± 1.04 (10) | 18.3 |
| | Bpa neg. | n.a. | n.a. | 0 (0) | n.a. |
| | Se-AbK# | 6.6 (1) | 18.1 (1) | 0.28 (2) | 4.1 |
| | Se-AbK neg. | n.a. | n.a. | 0 (0) | n.a. |
| G12# | AzF# | 6.54 ± 0.06 (13) | 7.61 ± 2.53 (13) | 2.04 ± 1.68 (14) | 14.6 |
| | AzF neg. | n.a. | n.a. | 0 (0) | n.a. |
| | Bpa# | 6.57 ± 0.07 (4) | 8.53 ± 5.01 (3) | 1.45 ± 1.56 (6) | 15.1 |
| | Bpa neg. | n.a. | n.a. | 0 (0) | n.a. |
| | Se-AbK# | 6.54 ± 0.08 (5) | 3.93 (2) | 0.47 ± 0.35 (6) | 6.7 |
| | Se-AbK neg. | n.a. | n.a. | 0 (0) | n.a. |

| Position replaced by TAG | ncAA | mean $pH_{50}$ ± S.D. (n) | mean $n_H$ ± S.D. (n) | mean Imax (nA) ± S.D. (n) | average TE (%) |
|---|---|---|---|---|---|
| V13 | AzF | 6.60 ± 0.07 (8) | 7.28 ± 1.77 (6) | 3.66 ± 2.31 (8) | 6.9 |
| | AzF neg. | n.a. | n.a. | 0 (0) | n.a. |
| | Bpa | 6.61 ± 0.08 (10) | 5.28 ± 1.44 (10) | 4.92 ± 2.16 (10) | 12.1 |
| | Bpa neg. | n.a. | n.a. | 0.1 (1) | n.a. |
| | Se-AbK# | 6.58 ± 0.01 (7) | 8.3 ± 1.02 (4) | 1.02 ± 1.25 (8) | 5 |
| | Se-AbK neg. | n.a. | n.a. | 0 (0) | n.a. |
| Q14 | AzF | 6.61 ± 0.05 (13) | 7.23 ± 1.66 (13) | 3.75 ± 1.48 (15) | 8.4 |
| | AzF neg. | n.a. | n.a. | 0 (0) | n.a. |
| | Bpa | 6.67 ± 0.2 (3) | 4.49 (2) | 4.05 ± 0.96 (3) | 6.6 |
| | Bpa neg. | n.a. | n.a. | 0 (0) | n.a. |
| | Se-AbK# | 6.63 ± 0.04 (8) | 6.85 ± 1.38 (6) | 1.47 ± 1.09 (10) | 8.4 |
| | Se-AbK neg. | n.a. | n.a. | 0 (0) | n.a. |
| P15 | AzF# | 6.59 ± 0.06 (11) | 6.27 ± 2.16 (11) | 3.52 ± 1.47 (12) | 20.4 |
| | AzF neg. | n.a. | n.a. | 0 (0) | n.a. |
| | Bpa | 6.52 ± 0.1 (4) | 6.47 ± 3.5 (4) | 2.2 ± 2.2 (5) | 14 |
| | Bpa neg. | n.a. | n.a. | 0 (0) | n.a. |
| | Se-AbK | n.a. | n.a. | 0 (0) | 6.2 |
| | Se-AbK neg. | n.a. | n.a. | 0 (0) | n.a. |
| V16 | AzF# | 6.60 (2) | 6.81 (2) | 0.82 ± 0.82 (5) | 5.2 |
| | AzF neg. | n.a. | n.a. | 0 (0) | n.a. |
| | Bpa | 6.58 ± 0.06 (7) | 5.95 ± 2.08 (7) | 3.60 ± 2.49 (7) | 12.2 |
| | Bpa neg. | n.a. | n.a. | 0 (0) | n.a. |
| | Se-AbK# | 6.57 ± 0.03 (4) | 18.1 ± 13.38 (4) | 1.14 ± 0.63 (4) | 13.1 |
| | Se-AbK neg. | n.a. | n.a. | 0 (0) | n.a. |
| S17 | AzF | n.a. | n.a. | 0 (0) | 4.7 |
| | AzF neg. | n.a. | n.a. | 0 (0) | n.a. |
| | Bpa | n.a. | n.a. | 0 (0) | 12.6 |
| | Bpa neg. | n.a. | n.a. | 0 (0) | n.a. |
| | Se-AbK | n.a. | n.a. | 0 (0) | 5.4 |
| | Se-AbK neg. | n.a. | n.a. | 0 (0) | n.a. |
| I18 | AzF | 6.57 ± 0.07 (7) | 5.71 ± 1.92 (7) | 3.07 ± 1.47 (8) | 4.5 |
| | AzF neg. | n.a. | n.a. | 0 (0) | n.a. |
| | Bpa# | 6.58 ± 0.08 (9) | 6.4 ± 1.6 (8) | 3.81 ± 1.48 (9) | 16.3 |
| | Bpa neg. | n.a. | n.a. | 0 (0) | n.a. |
| | Se-AbK | n.a. | n.a. | 0.14 (2) | 3.4 |
| | Se-AbK neg. | n.a. | n.a. | 0 (0) | n.a. |
| Q19 | AzF# | n.a. | n.a. | 0.39 ± 0.46 (3) | 3.6 |
| | AzF neg. | n.a. | n.a. | 0 (0) | n.a. |
| | Bpa# | 6.60 (2) | 8.99 (2) | 2.19 ± 1.72 (4) | 10.2 |
| | Bpa neg. | n.a. | n.a. | 0 (0) | n.a. |
| | Se-AbK | n.a. | n.a. | 0 (0) | 5.3 |
| | Se-AbK neg. | n.a. | n.a. | 0 (0) | n.a. |

| Position replaced by TAG | ncAA | mean $pH_{50}$ ± S.D. (n) | mean $n_H$ ± S.D. (n) | mean Imax (nA) ± S.D. (n) | average TE (%) |
|---|---|---|---|---|---|
| A20 | AzF# | n.a. | n.a. | 0.13 ± 0.13 (4) | 6.1 |
| | AzF neg. | n.a. | n.a. | 0 (0) | n.a. |
| | Bpa# | 6.54 (1) | 6.54 (1) | 0.27 (2) | 17.2 |
| | Bpa neg. | n.a. | n.a. | 0 (0) | n.a. |
| | Se-AbK | n.a. | n.a. | 0 (0) | 6.3 |
| | Se-AbK neg. | n.a. | n.a. | 0 (0) | n.a. |
| F21 | AzF | 6.57 ± 0.06 (5) | 7.13 ± 5.67 (5) | 3.47 ± 1.03 (5) | 15.6 |
| | AzF neg. | n.a. | n.a. | 0 (0) | n.a. |
| | Bpa# | 6.56 ± 0.06 (3) | 4.96 (2) | 1.01 ± 1.08 (9) | 17.3 |
| | Bpa neg. | n.a. | n.a. | 0 (0) | n.a. |
| | Se-AbK | n.a. | n.a. | 0 (0) | 5.3 |
| | Se-AbK neg. | n.a. | n.a. | 0 (0) | n.a. |
| A22 | AzF# | n.a. | n.a. | 0.43 ± 0.25 (7) | 12.3 |
| | AzF neg. | n.a. | n.a. | 0 (0) | n.a. |
| | Bpa# | 6.68 (1) | 7.28 (1) | 0.3 (2) | 9.6 |
| | Bpa neg. | n.a. | n.a. | 0 (0) | n.a. |
| | Se-AbK | n.a. | n.a. | 0 (0) | 8.1 |
| | Se-AbK neg. | n.a. | n.a. | 0 (0) | n.a. |
| S23# | AzF# | 6.58 (1) | 26.2 (1) | 0.38 ± 0.21 (3) | 8.2 |
| | AzF neg. | n.a. | n.a. | 0 (0) | n.a. |
| | Bpa# | n.a. | n.a. | 0.36 ± 0.45 (4) | 16.9 |
| | Bpa neg. | n.a. | n.a. | 0 (0) | n.a. |
| | Se-AbK# | 6.56 ± 0.04 (4) | 16.7 ± 9.85 (4) | 0.46 ± 0.59 (6) | 15.1 |
| | Se-AbK neg. | n.a. | n.a. | 0 (0) | n.a. |
| S24# | AzF# | 6.58 ± 0.03 (5) | 21.1 ± 6.56 (5) | 0.99 ± 1.08 (9) | 16 |
| | AzF neg. | n.a. | n.a. | 0 (0) | n.a. |
| | Bpa# | 6.57 ± 0.03 (5) | 10.8 ± 8.04 (5) | 3.32 ± 1.26 (6) | 12.1 |
| | Bpa neg. | n.a. | n.a. | 0 (0) | n.a. |
| | Se-AbK# | 6.60 ± 0.04 (9) | 19.4 ± 10.2 (9) | 3.02 ± 2.37 (11) | 13.7 |
| | Se-AbK neg. | n.a. | n.a. | 0 (0) | n.a. |
| S25 | AzF | n.a. | n.a. | 0 (0) | 13.8 |
| | AzF neg. | n.a. | n.a. | 0 (0) | n.a. |
| | Bpa | n.a. | n.a. | 0 (0) | 16.6 |
| | Bpa neg. | n.a. | n.a. | 0 (0) | n.a. |
| | Se-AbK | n.a. | n.a. | 0 (0) | 6.2 |
| | Se-AbK neg. | n.a. | n.a. | 0 (0) | n.a. |
| T26 | AzF | n.a. | n.a. | 0 (0) | 5.7 |
| | AzF neg. | n.a. | n.a. | 0 (0) | n.a. |
| | Bpa | n.a. | n.a. | 0 (0) | 12.6 |
| | Bpa neg. | n.a. | n.a. | 0 (0) | n.a. |
| | Se-AbK | n.a. | n.a. | 0 (0) | 6.7 |
| | Se-AbK neg. | n.a. | n.a. | 0 (0) | n.a. |

| Position replaced by TAG | ncAA | mean $pH_{50}$ ± S.D. (n) | mean $n_H$ ± S.D. (n) | mean Imax (nA) ± S.D. (n) | average TE (%) |
|---|---|---|---|---|---|
| L27 | AzF# | n.a. | n.a. | 1.06 ± 1.03 (4) | 11.6 |
| | AzF neg. | n.a. | n.a. | 0 (0) | n.a. |
| | Bpa# | n.a. | n.a. | 1.91 ± 3.03 (3) | 16.7 |
| | Bpa neg. | n.a. | n.a. | 0.15 (1) | n.a. |
| | Se-AbK | n.a. | n.a. | 0 (0) | 3.7 |
| | Se-AbK neg. | n.a. | n.a. | 0 (0) | n.a. |
| H28 | AzF | n.a. | n.a. | 0 (0) | 11.6 |
| | AzF neg. | n.a. | n.a. | 0 (0) | n.a. |
| | Bpa | n.a. | n.a. | 0 (0) | 15.5 |
| | Bpa neg. | n.a. | n.a. | 0 (0) | n.a. |
| | Se-AbK | n.a. | n.a. | 0 (0) | 5.8 |
| | Se-AbK neg. | n.a. | n.a. | 0 (0) | n.a. |
| G29 | AzF# | 6.59 (2) | 19.9 (2) | 0.11 ± 0.04 (4) | 13.5 |
| | AzF neg. | n.a. | n.a. | 0 (0) | n.a. |
| | Bpa# | 6.64 (1) | 6.14 (1) | 0.27 ± 0.11 (4) | 15.8 |
| | Bpa neg. | n.a. | n.a. | 0 (0) | n.a. |
| | Se-AbK | n.a. | n.a. | 0 (0) | 5.8 |
| | Se-AbK neg. | n.a. | n.a. | 0 (0) | n.a. |
| L30 | AzF# | 6.59 (2) | 6.16 (1) | 0.78 ± 1.19 (4) | 10.8 |
| | AzF neg. | n.a. | n.a. | 0 (0) | n.a. |
| | Bpa# | 6.61 ± 0.07 (5) | 5.09 ± 1.89 (5) | 2.91 ± 1.68 (5) | 15.8 |
| | Bpa neg. | n.a. | n.a. | 0 (0) | n.a. |
| | Se-AbK | n.a. | n.a. | 0 (0) | 8.6 |
| | Se-AbK neg. | n.a. | n.a. | 0 (0) | n.a. |
| A31# | AzF# | n.a. | n.a. | 0.19 ± 0.09 (5) | 13.1 |
| | AzF neg. | n.a. | n.a. | 0 (0) | n.a. |
| | Bpa# | n.a. | n.a. | 0.23 ± 0.1 (3) | 17.8 |
| | Bpa neg. | n.a. | n.a. | 0 (0) | n.a. |
| | Se-AbK# | 6.48 ± 0.09 (3) | 19.4 ± 12.2 (5) | 1.19 ± 2.06 (12) | 5.7 |
| | Se-AbK neg. | n.a. | n.a. | 0 (0) | n.a. |
| H32 | AzF# | n.a. | n.a. | 0.18 ± 0.08 (4) | 16.2 |
| | AzF neg. | n.a. | n.a. | 0 (0) | n.a. |
| | Bpa# | n.a. | n.a. | 0.11 (2) | 17.6 |
| | Bpa neg. | n.a. | n.a. | 0 (0) | n.a. |
| | Se-AbK | n.a. | n.a. | 0 (0) | 11.3 |
| | Se-AbK neg. | n.a. | n.a. | 0 (0) | n.a. |
| I33 | AzF# | 6.62 (1) | 4.86 (1) | 1.21 (1) | 20.2 |
| | AzF neg. | n.a. | n.a. | 0 (0) | n.a. |
| | Bpa | n.a. | n.a. | 0 (0) | 17.4 |
| | Bpa neg. | n.a. | n.a. | 0 (0) | n.a. |
| | Se-AbK | n.a. | n.a. | 0 (0) | 9.4 |
| | Se-AbK neg. | n.a. | n.a. | 0 (0) | n.a. |

| Position replaced by TAG | ncAA | mean $pH_{50}$ ± S.D. (n) | mean $n_H$ ± S.D. (n) | mean Imax (nA) ± S.D. (n) | average TE (%) |
|---|---|---|---|---|---|
| F34 | AzF | 6.48 ± 0.06 (8) | 12.6 ± 9.94 (9) | 2.31 ± 1.41 (9) | 12.6 |
| | AzF neg. | n.a. | n.a. | 0 (0) | n.a. |
| | Bpa# | 6.59 ± 0.07 (5) | 11.8 ± 10.4 (5) | 1.59 ± 1.55 (6) | 27.5 |
| | Bpa neg. | n.a. | n.a. | 0 (0) | n.a. |
| | Se-AbK | n.a. | n.a. | 0 (0) | 4.4 |
| | Se-AbK neg. | n.a. | n.a. | 0 (0) | n.a. |
| S35 | AzF# | 6.58 (1) | 29.6 (1) | 0.44 ± 0.53 (5) | 13.5 |
| | AzF neg. | n.a. | n.a. | 0 (0) | n.a. |
| | Bpa | n.a. | n.a. | 0 (0) | 28.9 |
| | Bpa neg. | n.a. | n.a. | 0 (0) | n.a. |
| | Se-AbK# | 6.48 ± 0.18 (3) | 6.67 ± 4.28 (3) | 0.37 ± 0.2 (6) | 16.3 |
| | Se-AbK neg. | n.a. | n.a. | 0 (0) | n.a. |
| Y36 | AzF# | n.a. | n.a. | 0.17 ± 0.12 (5) | 11 |
| | AzF neg. | n.a. | n.a. | 0 (0) | n.a. |
| | Bpa# | 6.72 ± 0.03 (3) | 4.32 ± 1.47 (3) | 0.66 ± 0.54 (4) | 14 |
| | Bpa neg. | n.a. | n.a. | 0 (0) | n.a. |
| | Se-AbK | n.a. | n.a. | 0 (0) | 7.2 |
| | Se-AbK neg. | n.a. | n.a. | 0 (0) | n.a. |
| E37 | AzF# | n.a. | n.a. | 0.4 (2) | 15.1 |
| | AzF neg. | n.a. | n.a. | 0 (0) | n.a. |
| | Bpa# | 6.75 (2) | 37.3 (2) | 0.61 ± 1.03 (4) | 17.2 |
| | Bpa neg. | n.a. | n.a. | 0 (0) | n.a. |
| | Se-AbK | n.a. | n.a. | 0 (0) | 12 |
| | Se-AbK neg. | n.a. | n.a. | 0 (0) | n.a. |
| R38 | AzF# | n.a. | n.a. | 0.54 ± 0.63 (4) | 18.4 |
| | AzF neg. | n.a. | n.a. | 0 (0) | n.a. |
| | Bpa# | 6.65 (1) | 23.7 (1) | 0.29 ± 0.21 (3) | 26.9 |
| | Bpa neg. | n.a. | n.a. | 0 (0) | n.a. |
| | Se-AbK | n.a. | n.a. | 0 (0) | 13.3 |
| | Se-AbK neg. | n.a. | n.a. | 0 (0) | n.a. |
| L39 | AzF | 6.44 ± 0.08 (16) | 5.92 ± 2.96 (16) | 3.2 ± 1.66 (16) | 9.2 |
| | AzF neg. | 6.44 (1) | 3.49 (1) | 0.57 (1) | n.a. |
| | Bpa# | 6.5 ± 0.11 (5) | 10.3 ± 12.5 (4) | 1.73 ± 1.48 (5) | 17.2 |
| | Bpa neg. | n.a. | n.a. | 0 (0) | n.a. |
| | Se-AbK | 6.45 ± 0.14 (8) | 7.87 ± 5.25 (10) | 1.39 ± 0.93 (10) | 11.7 |
| | Se-AbK neg. | n.a. | n.a. | 0 (0) | n.a. |
| S40 | AzF | 6.49 ± 0.11 (10) | 7.84 ± 9.64 (11) | 3.54 ± 1.51 (11) | 10.1 |
| | AzF neg. | 6.36 ± 0.19 (3) | 14.0 ± 16.6 (3) | 0.6 ± 0.68 (3) | n.a. |
| | Bpa | 6.63 ± 0.05 (3) | 15.9 ± 16.7 (3) | 4.01 ± 1.75 (3) | 27.3 |
| | Bpa neg.# | 6.42 ± 0.04 (3) | 5.97 ± 0.57 (3) | 0.48 ± 0.49 (3) | n.a. |
| | Se-AbK | 6.45 ± 0.1 (5) | 11.2 ± 8.07 (6) | 1.91 ± 0.99 (7) | 12.9 |
| | Se-AbK neg. | 6.42 ± 0.01 (3) | 4.19 ± 1.07 (3) | 0.1 ± 0.02 (3) | n.a. |

| Position replaced by TAG | ncAA | mean $pH_{50}$ ± S.D. (n) | mean $n_H$ ± S.D. (n) | mean Imax (nA) ± S.D. (n) | average TE (%) |
|---|---|---|---|---|---|
| L41 | AzF | 6.48 ± 0.07 (13) | 5.03 ± 1.61 (13) | 3.94 ± 1.69 (13) | 15.1 |
| | AzF neg. | n.a. | n.a. | 0.18 (1) | n.a. |
| | Bpa# | 6.62 ± 0.04 (4) | 9.98 ± 6.8 (4) | 3.31 ± 3.08 (4) | 27.6 |
| | Bpa neg. # | 6.49 (2) | 6.41 (2) | 2.03 (2) | n.a. |
| | Se-AbK | 6.51 ± 0.18 (6) | 6.44 ± 5.76 (7) | 1.61 ± 1.36 (7) | 11.2 |
| | Se-AbK neg. | n.a. | n.a. | 0 (0) | n.a. |
| K42 | AzF | n.a. | n.a. | 0 (0) | 13.4 |
| | AzF neg. | n.a. | n.a. | 0 (0) | n.a. |
| | Bpa | n.a. | n.a. | 0 (0) | 12.4 |
| | Bpa neg. | n.a. | n.a. | 0 (0) | n.a. |
| | Se-AbK# | 6.58 ± 0.08 (4) | 8.34 ± 5.15 (4) | 1.91 ± 1.5 (4) | 2 |
| | Se-AbK neg. | n.a. | n.a. | 0 (0) | n.a. |
| R43 | AzF | n.a. | n.a. | 0 (0) | 9.2 |
| | AzF neg. | n.a. | n.a. | 0 (0) | n.a. |
| | Bpa | n.a. | n.a. | 0 (0) | 7.4 |
| | Bpa neg. | n.a. | n.a. | 0 (0) | n.a. |
| | Se-AbK | n.a. | n.a. | 0 (0) | 7.2 |
| | Se-AbK neg. | n.a. | n.a. | 0 (0) | n.a. |
| A44 | AzF | 6.56 ± 0.2 (3) | 10.4 ± 9.48 (4) | 7.65 ± 4.45 (4) | 15.1 |
| | AzF neg. | n.a. | n.a. | 0 (0) | n.a. |
| | Bpa | 6.6 ± 0.05 (6) | 5.8 ± 1.9 (6) | 3.9 ± 2.04 (6) | 14.8 |
| | Bpa neg. | 6.26 (1) | 2.24 (1) | 3.41 (1) | n.a. |
| | Se-AbK# | 6.52 ± 0.04 (3) | 15.5 ± 9.84 (5) | 1.19 ± 0.38 (6) | 16.1 |
| | Se-AbK neg. | n.a. | n.a. | 0 (0) | n.a. |
| L45 | AzF# | 6.74 ± 0.1 (7) | 6.67 ± 2.76 (7) | 5.31 ± 1.91 (7) | 11.3 |
| | AzF neg. | n.a. | n.a. | 0 (0) | n.a. |
| | Bpa | 6.58 ± 0.05 (5) | 6.05 ± 4.19 (5) | 2.25 ± 0.52 (5) | 12.8 |
| | Bpa neg. | n.a. | n.a. | 0 (0) | n.a. |
| | Se-AbK | 6.35 ± 0.15 (8) | 4.47 ± 1.51 (8) | 2.42 ± 3.09 (8) | 16.5 |
| | Se-AbK neg. | n.a. | n.a. | 0 (0) | n.a. |
| W46 | AzF | n.a. | n.a. | 0 (0) | 13.9 |
| | AzF neg. | n.a. | n.a. | 0 (0) | n.a. |
| | Bpa | n.a. | n.a. | 0 (0) | 14 |
| | Bpa neg. | n.a. | n.a. | 0 (0) | n.a. |
| | Se-AbK | n.a. | n.a. | 0 (0) | 15.3 |
| | Se-AbK neg. | n.a. | n.a. | 0 (0) | n.a. |
| A47# | AzF# | 6.46 ± 0.2 (3) | 8.34 (1) | 2.49 ± 1.81 (8) | 15.9 |
| | AzF neg. | n.a. | n.a. | 0 (0) | n.a. |
| | Bpa# | 6.38 (2) | 8.88 (1) | 0.46 ± 0.34 (6) | 23 |
| | Bpa neg. | n.a. | n.a. | 0.31 (1) | n.a. |
| | Se-AbK | n.a. | n.a. | 0 (0) | 15.2 |
| | Se-AbK neg. | n.a. | n.a. | 0 (0) | n.a. |

| Position replaced by TAG | ncAA | mean $pH_{50}$ ± S.D. (n) | mean $n_H$ ± S.D. (n) | mean Imax (nA) ± S.D. (n) | average TE (%) |
|---|---|---|---|---|---|
| L48 | AzF | 6.56 ± 0.03 (3) | 7.72 ± 7.92 (3) | 5.9 ± 2.37 (3) | 12.8 |
| | AzF neg. | n.a. | n.a. | 0 (0) | n.a. |
| | Bpa | 6.57 ± 0.09 (5) | 7.45 ± 2.21 (5) | 3.38 ± 1.02 (5) | 18.2 |
| | Bpa neg. | n.a. | n.a. | 0.65 (2) | n.a. |
| | Se-AbK | 6.51 ± 0.1 (9) | 9.95 ± 7.87 (10) | 1.14 ± 1.29 (12) | 15.5 |
| | Se-AbK neg. | n.a. | n.a. | 0 (0) | n.a. |
| V65 | AzF | n.a. | n.a. | 0 (0) | 12.6 |
| | AzF neg. | n.a. | n.a. | 0 (0) | n.a. |
| | Bpa | 6.54 ± 0.07 (10) | 5.02 ± 1.46 (10) | 2.16 ± 1.46 (10) | 15.3 |
| | Bpa neg. | n.a. | n.a. | 0 (0) | n.a. |
| | Se-AbK[#] | n.a. | n.a. | 0.49 ± 0.05 (3) | 15.8 |
| | Se-AbK neg. | n.a. | n.a. | 0 (0) | n.a. |
| Q66 | AzF | 6.63 ± 0.14 (3) | 5.95 ± 3.13 (3) | 2.15 ± 0.58 (3) | 12.7 |
| | AzF neg. | n.a. | n.a. | 0 (0) | n.a. |
| | Bpa | 6.19 (1) | 5.79 (1) | 3.39 (2) | 11.6 |
| | Bpa neg. | n.a. | n.a. | 0 (0) | n.a. |
| | Se-AbK[#] | 6.51 ± 0.03 (5) | 4.82 ± 2.79 (5) | 0.74 ± 0.85 (9) | 12.2 |
| | Se-AbK neg. | n.a. | n.a. | 0 (0) | n.a. |
| Y67 | AzF | n.a. | n.a. | 0 (0) | 11.6 |
| | AzF neg. | n.a. | n.a. | 0 (0) | n.a. |
| | Bpa | 6.33 ± 0.09 (6) | 6.13 ± 1.15 (6) | 4.55 ± 1.40 (6) | 18.5 |
| | Bpa neg. | n.a. | n.a. | 0 (0) | n.a. |
| | Se-AbK[#] | 6.43 ± 0.16 (8) | 15.2 ± 10.3 (9) | 2.63 ± 3.81 (9) | 5.3 |
| | Se-AbK neg. | n.a. | n.a. | 0 (0) | n.a. |
| Y68 | AzF | n.a. | n.a. | 0 (0) | 10.0 |
| | AzF neg. | n.a. | n.a. | 0 (0) | n.a. |
| | Bpa[#] | 6.52 ± 0.12 (4) | 14.2 ± 9.48 (4) | 2.66 ± 1.68 (4) | 17.8 |
| | Bpa neg. | n.a. | n.a. | 0 (0) | n.a. |
| | Se-AbK | 6.23 ± 0.22 (3) | 2.61 ± 1.01 (3) | 0.16 ± 0.05 (3) | 6.8 |
| | Se-AbK neg. | n.a. | n.a. | 0 (0) | n.a. |
| F69 | AzF | 6.6 ± 0.09 (19) | 9.43 ± 8.98 (22) | 4.22 ± 2.25 (22) | 12.2 |
| | AzF neg. | n.a. | n.a. | 0 (0) | n.a. |
| | Bpa | 6.74 ± 0.08 (5) | 13.4 ± 13.6 (5) | 2.67 ± 1.42 (5) | 10.3 |
| | Bpa neg. | n.a. | n.a. | 0 (0) | n.a. |
| | Se-AbK[#] | 6.44 ± 0.05 (4) | 7.69 ± 0.52 (4) | 2.36 ± 1.84 (4) | 5.6 |
| | Se-AbK neg. | n.a. | n.a. | 0 (0) | n.a. |
| Y71 | AzF | 6.64 ± 0.1 (12) | 9.71 ± 8.17 (12) | 3.77 ± 2.00 (12) | 12.8 |
| | AzF neg. | 6.47 ± 0.16 (7) | 2.49 ± 1.1 (7) | 1.12 ± 1.04 (7) | n.a. |
| | Bpa[#] | 6.61 ± 0.15 (3) | 6.55 (2) | 2.97 ± 1.32 (3) | 11.5 |
| | Bpa neg. | n.a. | n.a. | 0 (0) | n.a. |
| | Se-AbK | 6.3 ± 0.14 (9) | 2.74 ± 1.44 (9) | 1.04 ± 1.36 (9) | 5.8 |
| | Se-AbK neg. | n.a. | n.a. | 0 (0) | n.a. |

| Position replaced by TAG | ncAA | mean $pH_{50}$ ± S.D. (n) | mean $n_H$ ± S.D. (n) | mean Imax (nA) ± S.D. (n) | average TE (%) |
|---|---|---|---|---|---|
| R175 | AzF | n.a. | n.a. | 0 (0) | 9.0 |
| | AzF neg. | n.a. | n.a. | 0 (0) | n.a. |
| | Bpa | n.a. | n.a. | 0 (0) | 13.0 |
| | Bpa neg. | n.a. | n.a. | 0 (0) | n.a. |
| | Se-AbK[#] | 6.21 ± 0.06 (12) | 6.65 ± 8.52 (13) | 0.93 ± 1.13 (15) | 13.3 |
| | Se-AbK neg. | n.a. | n.a. | 0 (0) | n.a. |
| G176 | AzF | n.a. | n.a. | 0 (0) | 9.5 |
| | AzF neg. | n.a. | n.a. | 0 (0) | n.a. |
| | Bpa | n.a. | n.a. | 0 (0) | 28.9 |
| | Bpa neg. | n.a. | n.a. | 0 (0) | n.a. |
| | Se-AbK | n.a. | n.a. | 0 (0) | 15 |
| | Se-AbK neg. | n.a. | n.a. | 0 (0) | n.a. |
| E177 | AzF | 6.59 ± 0.11 (3) | 6.29 ± 3.53 (3) | 1.07 ± 0.47 (3) | 8.7 |
| | AzF neg. | n.a. | n.a. | 0 (0) | n.a. |
| | Bpa | n.a. | n.a. | 0 (0) | 13.4 |
| | Bpa neg. | n.a. | n.a. | 0 (0) | n.a. |
| | Se-AbK[#] | 6.49 ± 0.08 (5) | 15.7 ± 11.2 (5) | 1.26 ± 1.23 (5) | 7.7 |
| | Se-AbK neg. | n.a. | n.a. | 0 (0) | n.a. |
| T236 | AzF | 6.17 ± 0.14 (10) | 15.7 ± 14.8 (10) | 1.47 ± 1.1 (10) | 12.0 |
| | AzF neg. | n.a. | n.a. | 0 (0) | n.a. |
| | Bpa | 6.17 ± 0.08 (12) | 9.85 ± 10.7 (12) | 1.24 ± 0.89 (14) | 18.3 |
| | Bpa neg. | n.a. | n.a. | 0 (0) | n.a. |
| | Se-AbK | 6.25 ± 0.07 (8) | 13.5 ± 13.2 (8) | 1.60 ± 1.10 (8) | 11.8 |
| | Se-AbK neg. | n.a. | n.a. | 0 (0) | n.a. |
| T239 | AzF | 6.05 ± 0.12 (12) | 3.82 ± 1.38 (13) | 0.83 ± 0.84 (14) | 10.7 |
| | AzF neg. | n.a. | n.a. | 0 (0) | n.a. |
| | Bpa | 5.49 ± 0.13 (6) | 2.61 ± 0.62 (6) | 1.16 ± 1.17 (6) | 16.3 |
| | Bpa neg. | n.a. | n.a. | 0 (0) | n.a. |
| | Se-AbK | n.a. | n.a. | 0 (0) | 13.6 |
| | Se-AbK neg. | n.a. | n.a. | 0 (0) | n.a. |
| P286 | AzF | n.a. | n.a. | 0 (0) | 8.8 |
| | AzF neg. | n.a. | n.a. | 0 (0) | n.a. |
| | Bpa | n.a. | n.a. | 0 (0) | 19.8 |
| | Bpa neg. | n.a. | n.a. | 0 (0) | n.a. |
| | Se-AbK | n.a. | n.a. | 0 (0) | 12.4 |
| | Se-AbK neg. | n.a. | n.a. | 0 (0) | n.a. |
| W287 | AzF | 6.63 ± 0.1 (12) | 8.54 ± 7.35 (12) | 3.07 ± 1.75 (13) | 11.5 |
| | AzF neg. | n.a. | n.a. | 0 (0) | n.a. |
| | Bpa | n.a. | n.a. | 0 (0) | 14.9 |
| | Bpa neg. | n.a. | n.a. | 0 (0) | n.a. |
| | Se-AbK | n.a. | n.a. | 0 (0) | 12.9 |
| | Se-AbK neg. | n.a. | n.a. | 0 (0) | n.a. |

| Position replaced by TAG | ncAA | mean $pH_{50}$ ± S.D. (n) | mean $n_H$ ± S.D. (n) | mean Imax (nA) ± S.D. (n) | average TE (%) |
|---|---|---|---|---|---|
| K343# | AzF# | 6.46 ± 0.09 (6) | 12.0 ± 12.9 (6) | 0.63 ± 0.43 (7) | 11.9 |
| | AzF neg. | n.a. | n.a. | 0 (0) | n.a. |
| | Bpa# | 6.19 ± 0.04 (6) | 4.65 ± 0.96 (6) | 1.89 ± 1.26 (6) | 20.4 |
| | Bpa neg. | n.a. | n.a. | 0 (0) | n.a. |
| | Se-AbK# | 6.32 (2) | 29.7 (2) | 0.51 (2) | 12 |
| | Se-AbK neg. | n.a. | n.a. | 0 (0) | n.a. |
| E344 | AzF | 6.6 ± 0.05 (5) | 17.3 ± 8.73 (5) | 1.18 ± 1.55 (5) | 9.2 |
| | AzF neg. | n.a. | n.a. | 0 (0) | n.a. |
| | Bpa | 6.42 ± 0.05 (4) | 8.67 ± 6.19 (5) | 1.62 ± 1.34 (5) | 20.8 |
| | Bpa neg. | n.a. | n.a. | 0.5 ± 0.35 (3) | n.a. |
| | Se-AbK | 6.43 ± 0.14 (6) | 13.2 ± 11.7 (7) | 0.20 ± 0.09 (7) | 8.2 |
| | Se-AbK neg. | n.a. | n.a. | 0 (0) | n.a. |
| P348 | AzF | n.a. | n.a. | 0 (0) | 10.8 |
| | AzF neg. | n.a. | n.a. | 0 (0) | n.a. |
| | Bpa | n.a. | n.a. | 0 (0) | 20.1 |
| | Bpa neg. | n.a. | n.a. | 0 (0) | n.a. |
| | Se-AbK | n.a. | n.a. | 0 (0) | 7.0 |
| | Se-AbK neg. | n.a. | n.a. | 0 (0) | n.a. |
| D351 | AzF# | 6.23 ± 0.06 (7) | 10.1 ± 10.4 (7) | 0.84 ± 0.77 (8) | 10.9 |
| | AzF neg. | n.a. | n.a. | 0 (0) | n.a. |
| | Bpa | 6.17 ± 0.12 (3) | 5.52 ± 1.46 (3) | 0.24 ± 0.24 (3) | 17.8 |
| | Bpa neg. | n.a. | n.a. | 0 (0) | n.a. |
| | Se-AbK | 6.33 ± 0.06 (9) | 6.72 ± 7.76 (10) | 0.14 ± 0.15 (10) | 12.0 |
| | Se-AbK neg. | n.a. | n.a. | 0 (0) | n.a. |
| E355# | AzF# | 6.46 ± 0.11 (8) | 4.24 ± 1.69 (8) | 1.5 ± 1.91 (10) | 12.6 |
| | AzF neg. | n.a. | n.a. | 0 (0) | n.a. |
| | Bpa# | 6.53 ± 0.1 (11) | 6.51 ± 8.43 (11) | 1.33 ± 1.52 (14) | 17.0 |
| | Bpa neg. | n.a. | n.a. | 0 (0) | n.a. |
| | Se-AbK# | 6.87 ± 0.22 (3) | 21.5 (2) | 0.77 ± 0.73 (4) | 11.9 |
| | Se-AbK neg. | n.a. | n.a. | 0 (0) | n.a. |
| K356 | AzF | 6.18 ± 0.22 (8) | 9.18 ± 9.19 (8) | 2.37 ± 2.23 (8) | 10.4 |
| | AzF neg. | n.a. | n.a. | 0 (0) | n.a. |
| | Bpa | 6.07 ± 0.28 (8) | 11.2 ± 13.4 (8) | 1.45 ± 1.42 (8) | 16.8 |
| | Bpa neg. | n.a. | n.a. | 0.46 (1) | n.a. |
| | Se-AbK# | 6.26 ± 0.22 (6) | 10.2 ± 12.7 (6) | 0.73 ± 0.84 (7) | 9.8 |
| | Se-AbK neg. | n.a. | n.a. | 0 (0) | n.a. |
| D357 | AzF | 5.66 ± 0.26 (10) | 2.51 ± 0.57 (10) | 1.28 ± 1.3 (11) | 13.0 |
| | AzF neg. | n.a. | n.a. | 0 (0) | n.a. |
| | Bpa | 5.91 ± 0.1 (8) | 2.85 ± 0.65 (8) | 1.84 ± 2.02 (10) | 16.8 |
| | Bpa neg. | n.a. | n.a. | 0 (0) | n.a. |
| | Se-AbK | 5.81 ± 0.17 (6) | 2.19 ± 0.38 (6) | 0.35 ± 0.43 (6) | 11.4 |
| | Se-AbK neg. | n.a. | n.a. | 0 (0) | n.a. |

| Position replaced by TAG | ncAA | mean $pH_{50}$ ± S.D. (n) | mean $n_H$ ± S.D. (n) | mean Imax (nA) ± S.D. (n) | average TE (%) |
|---|---|---|---|---|---|
| Y426 | AzF | n.a. | n.a. | 0 (0) | 8.2 |
| | AzF neg. | n.a. | n.a. | 0 (0) | n.a. |
| | Bpa | n.a. | n.a. | 0 (0) | 33.5 |
| | Bpa neg. | n.a. | n.a. | 0 (0) | n.a. |
| | Se-AbK[#] | 6.43 ± 0.09 (4) | 6.37 ± 4.49 (4) | 0.68 ± 0.94 (5) | 9.5 |
| | Se-AbK neg. | n.a. | n.a. | 0 (0) | n.a. |
| E427 | AzF | n.a. | n.a. | 0 (0) | 10.9 |
| | AzF neg. | n.a. | n.a. | 0 (0) | n.a. |
| | Bpa | n.a. | n.a. | 0 (0) | 20.1 |
| | Bpa neg. | n.a. | n.a. | 0 (0) | n.a. |
| | Se-AbK | n.a. | n.a. | 0 (0) | 7.6 |
| | Se-AbK neg. | n.a. | n.a. | 0 (0) | n.a. |
| I428 | AzF[#] | 6.48 ± 0.14 (4) | 18.3 ± 15.1 (3) | 1.11 ± 0.82 (6) | 10.5 |
| | AzF neg. | n.a. | n.a. | 0 (0) | n.a. |
| | Bpa | n.a. | n.a. | 0 (0) | 25.3 |
| | Bpa neg. | n.a. | n.a. | 0 (0) | n.a. |
| | Se-AbK | n.a. | n.a. | 0 (0) | 6.4 |
| | Se-AbK neg. | n.a. | n.a. | 0 (0) | n.a. |
| A429 | AzF | n.a. | n.a. | 0 (0) | 12.3 |
| | AzF neg. | n.a. | n.a. | 0 (0) | n.a. |
| | Bpa | n.a. | n.a. | 0 (0) | 7.8 |
| | Bpa neg. | n.a. | n.a. | 0 (0) | n.a. |
| | Se-AbK | n.a. | n.a. | 0 (0) | 6.7 |
| | Se-AbK neg. | n.a. | n.a. | 0 (0) | n.a. |
| G430 | AzF | n.a. | n.a. | 0 (0) | 11.6 |
| | AzF neg. | n.a. | n.a. | 0 (0) | n.a. |
| | Bpa | n.a. | n.a. | 0 (0) | 7.7 |
| | Bpa neg. | n.a. | n.a. | 0 (0) | n.a. |
| | Se-AbK | 6.46 ± 0.13 (6) | 10.6 ± 13.3 (6) | 0.18 ± 0.24 (7) | 5.7 |
| | Se-AbK neg. | n.a. | n.a. | 0 (0) | n.a. |
| F442 | AzF[#] | 6.62 ± 0.06 (7) | 10.9 ± 6.8 (7) | 3.05 ± 2.5 (7) | 7.5 |
| | AzF neg. | n.a. | n.a. | 0 (0) | n.a. |
| | Bpa | 6.64 ± 0.03 (4) | 1.73 ± 6.19 (4) | 1.32 ± 1.4 (4) | 11.9 |
| | Bpa neg. | n.a. | n.a. | 0 (0) | n.a. |
| | Se-AbK | n.a. | n.a. | 0 (0) | 4.8 |
| | Se-AbK neg. | n.a. | n.a. | 0 (0) | n.a. |
| F454 | AzF | n.a. | n.a. | 0 (0) | 9.4 |
| | AzF neg. | n.a. | n.a. | 0 (0) | n.a. |
| | Bpa[#] | 6.59 ± 0.06 (4) | 7.97 ± 1.72 (4) | 2.51 ± 1.79 (4) | 12.7 |
| | Bpa neg. | n.a. | n.a. | 0 (0) | n.a. |
| | Se-AbK | n.a. | n.a. | 0 (0) | 7.4 |
| | Se-AbK neg. | n.a. | n.a. | 0 (0) | n.a. |

| Position replaced by TAG | ncAA | mean $pH_{50}$ ± S.D. (n) | mean $n_H$ ± S.D. (n) | mean Imax (nA) ± S.D. (n) | average TE (%) |
|---|---|---|---|---|---|
| D455 | AzF | 6.56 ± 0.14 (7) | 6.81 ± 2.44 (7) | 0.83 ± 0.82 (8) | 9.7 |
| | AzF neg. | n.a. | n.a. | 0 (0) | n.a. |
| | Bpa# | 6.46 ± 0.05 (4) | 5.82 ± 2.38 (4) | 0.91 ± 0.88 (4) | 13.9 |
| | Bpa neg. | n.a. | n.a. | 0 (0) | n.a. |
| | Se-AbK | n.a. | n.a. | 0 (0) | 9.4 |
| | Se-AbK neg. | n.a. | n.a. | 0 (0) | n.a. |
| Y456 | AzF# | 6.55 ± 0.06 (4) | 13.8 ± 8.49 (4) | 0.9 ± 1.07 (5) | 7.1 |
| | AzF neg. | n.a. | n.a. | 0 (0) | n.a. |
| | Bpa# | 6.41 (1) | 7.66 (1) | 0.56 ± 0.47 (3) | 14.8 |
| | Bpa neg. | n.a. | n.a. | 0 (0) | n.a. |
| | Se-AbK | n.a. | n.a. | 0 (0) | 9.8 |
| | Se-AbK neg. | n.a. | n.a. | 0 (0) | n.a. |
| Y458 | AzF# | 6.54 ± 0.08 (5) | 17.5 ± 12.3 (5) | 2.48 ± 1.61 (6) | 9.9 |
| | AzF neg. | n.a. | n.a. | 0 (0) | n.a. |
| | Bpa | 6.46 ± 0.12 (13) | 6.22 ± 1.52 (13) | 1.86 ± 2.2 (13) | 10.1 |
| | Bpa neg. | n.a. | n.a. | 0 (0) | n.a. |
| | Se-AbK | 6.44 ± 0.03 (4) | 4.63 ± 2.28 (4) | 0.26 ± 0.27 (4) | 10.0 |
| | Se-AbK neg. | n.a. | n.a. | 0 (0) | n.a. |
| E459 | AzF | n.a. | n.a. | 0 (0) | 6 |
| | AzF neg. | n.a. | n.a. | 0 (0) | n.a. |
| | Bpa | n.a. | n.a. | 0 (0) | 8 |
| | Bpa neg. | n.a. | n.a. | 0 (0) | n.a. |
| | Se-AbK | n.a. | n.a. | 0 (0) | 5.5 |
| | Se-AbK neg. | n.a. | n.a. | 0 (0) | n.a. |
| V460 | AzF | n.a. | n.a. | 0 (0) | 5.3 |
| | AzF neg. | n.a. | n.a. | 0 (0) | n.a. |
| | Bpa | n.a. | n.a. | 0 (0) | 10.4 |
| | Bpa neg. | n.a. | n.a. | 0 (0) | n.a. |
| | Se-AbK | n.a. | n.a. | 0 (0) | 3.7 |
| | Se-AbK neg. | n.a. | n.a. | 0 (0) | n.a. |
| I461 | AzF# | 6.47 ± 0.13 (5) | 3.93 ± 1.53 (4) | 0.97 ± 0.88 (5) | 6.9 |
| | AzF neg. | n.a. | n.a. | 0 (0) | n.a. |
| | Bpa# | 6.59 ± 0.06 (3) | 11.8 ± 6.01 (3) | 1.97 ± 1.74 (3) | 14.0 |
| | Bpa neg. | n.a. | n.a. | 0 (0) | n.a. |
| | Se-AbK | n.a. | n.a. | 0 (0) | 5.0 |
| | Se-AbK neg. | n.a. | n.a. | 0 (0) | n.a. |
| K462 | AzF# | 6.56 ± 0.13 (3) | 6.13 (2) | 3.37 ± 1.87 (4) | 7.6 |
| | AzF neg. | n.a. | n.a. | 0 (0) | n.a. |
| | Bpa | 6.63 (2) | 6.4 (2) | 5.12 (2) | 9.6 |
| | Bpa neg. | n.a. | n.a. | 0 (0) | n.a. |
| | Se-AbK | n.a. | n.a. | 0 (0) | 4.8 |
| | Se-AbK neg. | n.a. | n.a. | 0 (0) | n.a. |

| Position replaced by TAG | ncAA | mean $pH_{50}$ ± S.D. (n) | mean $n_H$ ± S.D. (n) | mean Imax (nA) ± S.D. (n) | average TE (%) |
|---|---|---|---|---|---|
| H463 | AzF# | 6.53 ± 0.09 (3) | 15.8 ± 17.0 (3) | 3.27 ± 1.47 (4) | 8.0 |
| | AzF neg. | n.a. | n.a. | 0 (0) | n.a. |
| | Bpa | n.a. | n.a. | 0 (0) | 7.6 |
| | Bpa neg. | n.a. | n.a. | 0 (0) | n.a. |
| | Se-AbK | n.a. | n.a. | 0 (0) | 13.0 |
| | Se-AbK neg. | n.a. | n.a. | 0 (0) | n.a. |
| K464 | AzF# | 6.40 ± 0.16 (5) | 12.6 ± 5.74 (3) | 1.56 ± 0.96 (7) | 8.4 |
| | AzF neg. | n.a. | n.a. | 0 (0) | n.a. |
| | Bpa | n.a. | n.a. | 0 (0) | 7.8 |
| | Bpa neg. | n.a. | n.a. | 0 (0) | n.a. |
| | Se-AbK | n.a. | n.a. | 0 (0) | 11.9 |
| | Se-AbK neg. | n.a. | n.a. | 0 (0) | n.a. |
| L465 | AzF# | 6.5 ± 0.12 (4) | 23.3 ± 6.24 (4) | 0.91 ± 0.93 (6) | 6.5 |
| | AzF neg. | n.a. | n.a. | 0 (0) | n.a. |
| | Bpa | n.a. | n.a. | 0 (0) | 5.6 |
| | Bpa neg. | n.a. | n.a. | 0 (0) | n.a. |
| | Se-AbK | n.a. | n.a. | 0 (0) | 16.3 |
| | Se-AbK neg. | n.a. | n.a. | 0 (0) | n.a. |
| C466 | AzF | n.a. | n.a. | 0 (0) | 5.8 |
| | AzF neg. | n.a. | n.a. | 0 (0) | n.a. |
| | Bpa# | 6.69 (2) | 29.7 (1) | 2.31 (2) | 6.6 |
| | Bpa neg. # | 6.76 ± 0.07 (3) | 11.2 ± 11.4 (3) | 0.98 ± 0.59 (3) | n.a. |
| | Se-AbK# | 6.46 (1) | 4.05 (1) | 0.41 ± 0.16 (3) | 12.4 |
| | Se-AbK neg.# | 6.24 ± 0.09 (3) | 10.5 ± 9.44 (3) | 2.29 ± 1.12 (6) | n.a. |
| R467 | AzF | 6.74 ± 0.05 (4) | 4.9 ± 1.55 (5) | 4.97 ± 1.16 (5) | 8.2 |
| | AzF neg. | 6.64 ± 0.02 (5) | 5.7 ± 2.76 (5) | 6.15 ± 3.24 (5) | n.a. |
| | Bpa | 6.75 ± 0.04 (6) | 6.26 ± 0.82 (6) | 5.62 ± 1.86 (6) | 6.4 |
| | Bpa neg. # | 6.69 (1) | 6.35 (1) | 4.78 (1) | n.a. |
| | Se-AbK# | 6.51 ± 0.17 (4) | 6.46 ± 0.76 (3) | 2.69 ± 1.77 (5) | 8.1 |
| | Se-AbK neg. | 6.61 ± 0.03 (3) | 5.68 ± 2.69 (3) | 4.34 ± 1.58 (3) | n.a. |
| R468 | AzF# | 6.72 ± 0.11 (6) | 7.01 ± 0.54 (6) | 4.07 ± 1.13 (6) | 10.0 |
| | AzF neg. | 6.71 ± 0.04 (7) | 7.71 ± 0.9 (7) | 7.22 ± 2.16 (7) | n.a. |
| | Bpa | 6.73 ± 0.1 (7) | 7.87 ± 6.96 (7) | 4.95 ± 2.05 (8) | 6.8 |
| | Bpa neg. | 6.76 (2) | 6.11 (2) | 4.91 (2) | n.a. |
| | Se-AbK | 6.51 ± 0.12 (5) | 9.53 ± 6.54 (5) | 3.42 ± 2.44 (5) | 10.2 |
| | Se-AbK neg. | 6.53 ± 0.1 (4) | 12.4 ± 12.8 (5) | 3.97 ± 2.27 (5) | n.a. |
| G469 | AzF | 6.75 ± 0.13 (6) | 8.51 ± 4.49 (6) | 5.16 ± 1.77 (6) | 7.3 |
| | AzF neg. | 6.74 ± 0.06 (5) | 6.77 ± 2.16 (5) | 7.01 ± 2.36 (5) | n.a. |
| | Bpa | 6.78 ± 0.04 (3) | 6.04 ± 0.8 (3) | 5.91 ± 1.62 (3) | 7.4 |
| | Bpa neg. # | 6.71 (1) | 7.63 (1) | 4.72 (1) | n.a. |
| | Se-AbK | 6.51 ± 0.08 (5) | 12.4 ± 12.1 (5) | 2.51 ± 2.8 (5) | 6.5 |
| | Se-AbK neg. | 6.51 ± 0.03 (3) | 5.79 ± 1.88 (3) | 8.71 ± 1.89 (3) | n.a. |

| Position replaced by TAG | ncAA | mean $pH_{50}$ ± S.D. (n) | mean $n_H$ ± S.D. (n) | mean Imax (nA) ± S.D. (n) | average TE (%) |
|---|---|---|---|---|---|
| K470 | AzF | 6.33 ± 0.07 (5) | 6.16 ± 1.7 (5) | 2.57 ± 1.33 (6) | 6.1 |
| | AzF neg. | 6.61 ± 0.08 (8) | 9.58 ± 10.46 (8) | 6.38 ± 2.83 (10) | n.a. |
| | Bpa | 6.77 ± 0.05 (4) | 5.10 ± 1.68 (4) | 4 ± 1.71 (4) | 6.3 |
| | Bpa neg. | 6.66 ± 0.04 (4) | 7.62 ± 2.08 (4) | 3.75 ± 0.86 (4) | n.a. |
| | Se-AbK# | 6.56 ± 0.05 (3) | 5.08 ± 2.05 (3) | 3.79 ± 1.75 (3) | 6.7 |
| | Se-AbK neg. | 6.56 ± 0.05 (5) | 10.6 ± 8.66 (6) | 4.70 ± 1.69 (7) | n.a. |
| C471 | AzF | 6.60 ± 0.11 (10) | 12.4 ± 12.1 (10) | 2.84 ± 1.27 (10) | 8.4 |
| | AzF neg. | 6.62 ± 0.06 (6) | 9.42 ± 2.33 (6) | 4.34 ± 0.88 (6) | n.a. |
| | Bpa | 6.76 ± 0.04 (4) | 7.15 ± 3.55 (4) | 5.92 ± 1.60 (4) | 6.1 |
| | Bpa neg. | 6.7 (1) | 2.35 (1) | 4.06 (1) | n.a. |
| | Se-AbK# | 6.57 ± 0.11 (5) | 8.08 ± 0.69 (5) | 3.84 ± 2.23 (5) | 7.9 |
| | Se-AbK neg. | 6.48 ± 0.08 (3) | 13.4 ± 14.2 (4) | 4.54 (5) | n.a. |
| Q472 | AzF | 6.81 ± 0.11 (7) | 15.1 ± 11.2 (7) | 5.35 ± 1.68 (7) | 8.9 |
| | AzF neg. | 6.71 ± 0.06 (6) | 6.64 ± 2.3 (6) | 7.33 ± 1.88 (6) | n.a. |
| | Bpa | 6.69 ± 0.04 (4) | 6.25 ± 2.01 (4) | 5.58 ± 1.73 (4) | 8.0 |
| | Bpa neg. | 6.70 ± 0.03 (4) | 6.38 ± 1.61 (4) | 2.70 ± 1.32 (4) | n.a. |
| | Se-AbK# | 6.49 ± 0.22 (3) | 6.59 (2) | 3.99 ± 0.16 (3) | 8.1 |
| | Se-AbK neg.# | 6.49 (2) | 15.4 (2) | 4.5 (2) | n.a. |
| K473 | AzF | 6.65 ± 0.05 (9) | 6.58 ± 2.14 (9) | 3.81 ± 1.34 (9) | 7.4 |
| | AzF neg. | 6.68 (2) | 7.94 (2) | 4.99 (2) | n.a. |
| | Bpa | 6.86 ± 0.04 (3) | 10.6 ± 8.35 (3) | 6 ± 0.97 (3) | 8.9 |
| | Bpa neg. | 6.91 (2) | 13.1 (2) | 5.58 (2) | n.a. |
| | Se-AbK# | 6.58 ± 0.03 (6) | 17.6 ± 13.1 (6) | 4.55 ± 2.91 (6) | 14.0 |
| | Se-AbK neg. | 6.58 ± 0.09 (4) | 7.24 ± 1.81 (4) | 6.66 ± 1.66 (4) | n.a. |
| E474 | AzF | 6.63 ± 0.07 (6) | 6.88 ± 3.5 (7) | 3.24 ± 1.02 (8) | 8.5 |
| | AzF neg. | 6.69 ± 0.13 (8) | 7.1 ± 3.61 (8) | 4.38 ± 2.66 (8) | n.a. |
| | Bpa | 6.69 ± 0.04 (4) | 5.94 ± 1.3 (4) | 7.62 ± 2.64 (4) | 9.5 |
| | Bpa neg. | 6.51 ± 0.19 (4) | 11.4 ± 9.67 (4) | 3.86 ± 4.06 (4) | n.a. |
| | Se-AbK | 6.60 ± 0.03 (3) | 5.05 ± 1.59 (3) | 2.59 ± 0.49 (3) | 9.5 |
| | Se-AbK neg. | 6.58 ± 0.09 (3) | 6.23 ± 1.66 (3) | 3.35 ± 1.32 (3) | n.a. |
| A475 | AzF | 6.72 ± 0.18 (6) | 4.95 ± 1.94 (6) | 5.17 ± 2.08 (6) | 10.2 |
| | AzF neg. | 6.64 ± 0.18 (6) | 8.66 ± 7.95 (6) | 5.97 ± 3.69 (6) | n.a. |
| | Bpa | 6.67 ± 0.07 (4) | 7.41 ± 1.77 (4) | 8.25 ± 0.8 (4) | 13.7 |
| | Bpa neg. | 6.66 ± 0.03 (3) | 6.82 ± 2.09 (3) | 5.79 ± 4.77 (3) | n.a. |
| | Se-AbK | 6.58 ± 0.08 (8) | 5.14 ± 2.32 (8) | 4.70 ± 1.36 (8) | 7.6 |
| | Se-AbK neg. | 6.62 (2) | 7.02 (2) | 2.75 (2) | n.a. |
| K476 | AzF | 6.64 ± 0.05 (7) | 8.22 ± 6.76 (7) | 2.32 ± 1.49 (8) | 12.7 |
| | AzF neg. | 6.60 ± 0.20 (6) | 14.4 ± 13.4 (7) | 3.69 ± 2.24 (7) | n.a. |
| | Bpa# | 6.66 (1) | 9.13 (1) | 6.78 (1) | 10.3 |
| | Bpa neg. | 6.59 ± 0.09 (4) | 9.38 ± 5.83 (4) | 4.41 ± 4.26 (4) | n.a. |
| | Se-AbK | 6.59 ± 0.05 (9) | 6.83 ± 2.77 (9) | 3.37 ± 2.13 (9) | 9.6 |
| | Se-AbK neg. | 6.46 ± 0.14 (5) | 8.82 ± 0.81 (5) | 3.07 ± 2.34 (5) | n.a. |

| Position replaced by TAG | ncAA | mean $pH_{50}$ ± S.D. (n) | mean $n_H$ ± S.D. (n) | mean Imax (nA) ± S.D. (n) | average TE (%) |
|---|---|---|---|---|---|
| R477 | AzF | 6.54 (1) | 3.45 (1) | 3.11 (1) | 6.2 |
| | AzF neg. | 6.69 ± 0.14 (4) | 5.98 ± 1.1 (4) | 3.71 ± 1.98 (4) | n.a. |
| | Bpa# | 6.64 (2) | 7.72 (2) | 6.14 (2) | 10.2 |
| | Bpa neg. | 6.59 (2) | 7.86 (2) | 5.49 (2) | n.a. |
| | Se-AbK | 6.60 ± 0.11 (7) | 5.32 ± 2.41 (7) | 3.05 ± 1.25 (7) | 9.3 |
| | Se-AbK neg. | 6.59 ± 0.04 (3) | 6.36 ± 2.51 (3) | 5.29 ± 2.39 (3) | n.a. |
| S478 | AzF | 6.71 ± 0.04 (3) | 5.17 ± 3.15 (3) | 5.00 ± 2.89 (3) | 6.0 |
| | AzF neg. # | 6.9 (1) | 13.5 (1) | 2.33 (2) | n.a. |
| | Bpa | 6.74 ± 0.1 (4) | 11.9 ± 10.8 (4) | 6.11 ± 2.03 (4) | 6.9 |
| | Bpa neg. | 6.65 (1) | 7.3 (1) | 7.38 (1) | n.a. |
| | Se-AbK | 6.61 ± 0.08 (10) | 7.34 ± 1.39 (9) | 4.35 ± 2.37 (10) | 8.7 |
| | Se-AbK neg. | 6.58 (2) | 5.22 (2) | 7.51 (2) | n.a. |
| S479 | AzF | 6.68 ± 0.11 (8) | 5.55 ± 1.70 (7) | 5.34 ± 3.30 (8) | 4.9 |
| | AzF neg. # | 6.62 ± 0.04 (3) | 13.6 ± 9.99 (3) | 3.6 ± 2.41 (3) | n.a. |
| | Bpa | 6.64 ± 0.06 (4) | 7.29 ± 2.05 (4) | 6.28 ± 2.42 (4) | 8.8 |
| | Bpa neg. | 6.47 ± 0.18 (3) | 5.11 ± 2.55 (3) | 3.49 ± 2.47 (3) | n.a. |
| | Se-AbK | 6.62 ± 0.07 (4) | 7.11 ± 1.99 (4) | 4.3 ± 2.73 (4) | 11.9 |
| | Se-AbK neg. | 6.59 ± 0.06 (4) | 4.39 ± 0.83 (4) | 6.32 ± 3.3 (4) | n.a. |
| A480 | AzF | 6.67 (1) | 4.58 (1) | 10.4 (1) | 4.3 |
| | AzF neg. # | 6.71 (1) | 5.73 (1) | 5.5 (2) | n.a. |
| | Bpa | 6.67 ± 0.03 (4) | 6.01 ± 0.78 (4) | 7.80 ± 1.41 (4) | 13.1 |
| | Bpa neg. | 6.5 (2) | 7.94 (2) | 6.48 (2) | n.a. |
| | Se-AbK | 6.48 ± 0.1 (3) | 5.25 ± 4.47 (3) | 4.05 ± 3.03 (3) | 8.8 |
| | Se-AbK neg. | 6.61 (1) | 18.8 (2) | 1.53 (2) | n.a. |
| D481 | AzF | 6.51 (1) | 5.52 (1) | 3.4 (1) | 4.1 |
| | AzF neg. | 6.67 ± 0.04 (3) | 5.88 ± 0.42 (3) | 8.92 ± 2.66 (3) | n.a. |
| | Bpa | 6.71 ± 0.05 (4) | 6.83 ± 0.86 (4) | 6.27 ± 1.38 (4) | 8.2 |
| | Bpa neg. | 6.69 ± 0.12 (3) | 17.6 ± 15.4 (4) | 3.38 ± 3.92 (4) | n.a. |
| | Se-AbK | 6.63 ± 0.04 (8) | 9.97 ± 9.09 (8) | 3.4 ± 2.04 (8) | 12.9 |
| | Se-AbK neg. | 6.3 (1) | 6.06 (1) | 2.96 (1) | n.a. |
| K482 | AzF | 6.59 ± 0.27 (6) | 5.74 ± 2.9 (6) | 4.04 ± 1.11 (6) | 5.6 |
| | AzF neg. | 6.64 ± 0.07 (8) | 7.65 ± 2.16 (8) | 4.24 ± 1.14 (8) | n.a. |
| | Bpa | 6.67 (2) | 6.32 (2) | 6.6 ± 2.37 (3) | 7.7 |
| | Bpa neg. | 6.62 (2) | 15.8 ± 15.6 (3) | 4.50 ± 3.55 (3) | n.a. |
| | Se-AbK | 6.69 ± 0.07 (7) | 8.34 ± 7.09 (8) | 4.58 ± 2.04 (8) | 7.7 |
| | Se-AbK neg. | 6.5 ± 0.08 (4) | 5.52 ± 0.87 (4) | 5.1 ± 1.86 (4) | n.a. |
| G483 | AzF# | 6.66 ± 0.08 (11) | 8.24 ± 7.35 (11) | 3.73 ± 2.04 (12) | 5.3 |
| | AzF neg. | 6.68 ± 0.11 (7) | 10.1 ± 8.32 (7) | 3.2 ± 1.34 (7) | n.a. |
| | Bpa | 6.65 ± 0.09 (4) | 7.42 ± 2.44 (4) | 6.87 ± 3.35 (4) | 8.4 |
| | Bpa neg. | 6.59 ± 0.04 (5) | 13.7 ± 12.2 (5) | 7.94 ± 1.69 (5) | n.a. |
| | Se-AbK# | 6.67 ± 0.1 (10) | 8.0 ± 2.33 (10) | 4.54 ± 2.43 (10) | 7.8 |
| | Se-AbK neg. | 6.56 ± 0.06 (4) | 5.96 ± 1.34 (4) | 4.86 ± 2.09 (4) | n.a. |

| Position replaced by TAG | ncAA | mean $pH_{50}$ ± S.D. (n) | mean $n_H$ ± S.D. (n) | mean Imax (nA) ± S.D. (n) | average TE (%) |
|---|---|---|---|---|---|
| V484 | AzF | n.a. | n.a. | 0 (0) | 8.1 |
| | AzF neg. | n.a. | n.a. | 0 (0) | n.a. |
| | Bpa# | 6.56 ± 0.09 (4) | 7.39 ± 1.6 (4) | 2.47 ± 2.91 (4) | 9.8 |
| | Bpa neg. | n.a. | n.a. | 0 (0) | n.a. |
| | Se-AbK | n.a. | n.a. | 0 (0) | 10.8 |
| | Se-AbK neg. | n.a. | n.a. | 0 (0) | n.a. |
| A485 | AzF# | 6.74 ± 0.09 (6) | 9.34 ± 8.72 (6) | 4.04 ± 1.21 (7) | 6.1 |
| | AzF neg. | 6.73 ± 0.13 (8) | 14.5 ± 11.5 (8) | 3.25 ± 1.4 (9) | n.a. |
| | Bpa | 6.51 ± 0.12 (5) | 14.7 ± 11.0 (5) | 4.75 ± 2.9 (5) | 14.7 |
| | Bpa neg. | 6.7 ± 0.06 (4) | 6.55 ± 1.32 (4) | 3.2 ± 1.97 (4) | n.a. |
| | Se-AbK | 6.6 ± 0.14 (5) | 11.9 ± 8.85 (5) | 5.17 ± 2.44 (5) | 11.1 |
| | Se-AbK neg. | 6.53 ± 0.06 (4) | 6.69 ± 3.22 (4) | 2.84 ± 2.82 (4) | n.a. |
| L486 | AzF | 6.8 ± 0.08 (8) | 7.68 ± 5.14 (8) | 6.23 ± 2.71 (9) | 6.8 |
| | AzF neg. | 6.73 ± 0.07 (7) | 7.42 ± 0.92 (7) | 5.95 ± 2.58 (7) | n.a. |
| | Bpa# | 6.73 ± 0.11 (7) | 14.2 ± 9.56 (7) | 3.69 ± 2 (7) | 14.4 |
| | Bpa neg. | 6.6 (2) | 6.31 (2) | 6.35 (2) | n.a. |
| | Se-AbK# | 6.59 (2) | 7.05 (2) | 4.01 (2) | 10.2 |
| | Se-AbK neg. | 6.6 ± 0.06 (4) | 5.14 ± 1.04 (4) | 2.9 ± 2.01 (5) | n.a. |
| S487 | AzF# | n.a. | n.a. | 1.01 ± 1.13 (5) | 9.3 |
| | AzF neg. | n.a. | n.a. | 0 (0) | n.a. |
| | Bpa# | n.a. | n.a. | 2.28 ± 1.98 (6) | 18.1 |
| | Bpa neg. # | 6.49 (2) | 11.6 (2) | 0.49 (2) | n.a. |
| | Se-AbK | n.a. | n.a. | 0 (0) | 9.4 |
| | Se-AbK neg. | n.a. | n.a. | 0 (0) | n.a. |
| L488 | AzF# | 6.78 (1) | 6.25 (1) | 4.91 (1) | 6.0 |
| | AzF neg. # | 6.69 (1) | 7.85 (1) | 4.51 (2) | n.a. |
| | Bpa | 6.66 ± 0.19 (3) | 12.0 ± 12.1 (3) | 4.14 ± 2.14 (3) | 10.7 |
| | Bpa neg. | 6.6 (1) | 5.31 (1) | 6.84 (1) | n.a. |
| | Se-AbK# | 6.65 ± 0.02 (3) | 13.4 ± 10.3 (3) | 5.56 ± 3.17 (3) | 8.0 |
| | Se-AbK neg. | 6.67 (1) | 11.6 (1) | 5.9 (1) | n.a. |

Table S2: Mean current size of P2X2 and GluA2 variants evaluated in the APC screen. Values are depicted as mean ± S.D., number in brackets indicates number of cells.

| Clone | Current (nA) at 30 µM agonist | Current (nA) at saturating agonist |
|---|---|---|
| P2X2 WT | 8.14 ± 2.87 (26) | 6.01 ± 2.86 (26) |
| P2X2 K296AzF | 3.79 ± 3.44 (40) | 3.07 ± 2.60 (40) |
| P2X2 K296X w/o AzF | 0.78 ± 0.56 (2) | 1.28 ± 0.69 (2) |
| P2X2 K296Bpa | 3.27 ± 3.04 (51) | 2.25 ± 2.31 (51) |
| P2X2 K296X w/o Bpa | 0.06 ± 0.03 (2) | 0.34 ± 0.21 (2) |
| GluA2 WT | 1.04 ± 1.31 (46) | 5.69 ± 3.76 (47) |
| GluA2 Y533AzF | 0.09 ± 0.22 (27) | 1.21 ± 0.96 (30) |
| GluA2 Y533X w/o AzF | 0.01 ± 0.01 (2) | 0.23 ± 0.15 (2) |
| GluA2 S729AzF | 0.03 ± 0.08 (15) | 0.39 ± 0.33 (17) |
| GluA2 S729X w/o AzF | No current | No current |
| GluA2 S729Bpa | 0.07 ± 0.13 (13) | 0.28 ± 0.24 (16) |
| GluA2 S729X w/o Bpa | No current | No current |

Table S3: Incorporation of ncAA photocrosslinkers in the acidic pocket of hASIC1a results in channel variants with accelerated current decay. Displayed are mean and S.D. of t½, (n) equals number of cells. (*) denotes significant difference between t½ of current decay compared to WT, $p < 0.05$; (***): $p < 0.001$; Mann-Whitney test.

| Clone | t½ ± S.D. (ms) | P value | n |
|---|---|---|---|
| hASIC1a WT | 818 ± 750 | - | 9 |
| T239Bpa | 224 ± 176* | 0.036 | 6 |
| D357AzF | 93.8 ± 40.9*** | 0.0002 | 10 |

Table S4: SSD curve recordings of different hASIC1a variants measured by APC. Displayed are mean and S.D. of half-maximal inactivation ($pH_{50}$ SSD) and Hill slope ($n_H$) as well as number of experiments (n). (*) denotes significant difference from WT, $p < 0.05$, (**): $p < 0.005$, (***): $p < 0.001$, (****): $p < 0.0001$, Mann-Whitney test.

| Clone | $pH_{50}$ SSD ± S.D. | P value | $n_H$ SSD ± S.D. | P value | n |
|---|---|---|---|---|---|
| WT | 6.91 ± 0.02 | - | 3.16 ± 0.42 | - | 40 |
| E177Bpa | 7.15 ± 0.01**** | <0.0001 | 7.95 ± 3.3** | 0.0035 | 12 |
| T236AzF | 6.85 ± 0.02 | 0.5541 | 5.68 ± 1.35 | 0.1134 | 4 |
| T236Bpa | 6.85 ± 0.02 | 0.3506 | 7.37 ± 1.67** | 0.0092 | 16 |
| T239Bpa | 6.79 ± 0.07 | 0.3447 | 2.02 ± 0.61 | 0.1587 | 12 |
| K343Bpa | 6.97 ± 0.04 | 0.2135 | 2.31 ± 0.59 | 0.3279 | 14 |
| E344AzF | 7.0 ± 0.04 | 0.1558 | 4.87 ± 1.9 | 0.4739 | 5 |
| E344Bpa | 7.07 ± 0.02*** | 0.0004 | 5.31 ± 0.76* | 0.0876 | 12 |
| D351Bpa | 6.98 ± 0.03 | 0.7032 | 3.73 ± 0.87 | 0.7474 | 7 |
| E355AzF | 7.07 ± 0.03* | 0.0459 | 2.56 ± 0.6 | 0.1640 | 8 |
| E355Bpa | 7.11 ± 0.01**** | <0.0001 | 5.0 ± 0.52 | 0.4647 | 7 |
| K356AzF | 6.76 ± 0.06 | 0.2881 | 4.32 ± 2.5 | 0.3639 | 7 |
| K356Bpa | 6.81 ± 0.02 | 0.1406 | 9.07 ± 4.45 | 0.4252 | 12 |
| D357AzF | 6.70 ± 0.50 | 0.2394 | 16.2 ± 11.9 | 0.7819 | 6 |
| D357Bpa | 6.84 ± 0.03 | 0.3937 | 12.9 ± 4.8**** | <0.0001 | 13 |
| F69Bpa | 7.01 ± 0.06* | 0.0175 | 13.3 ± 13.9*** | 0.0002 | 8 |
| Y71AzF | 7.05 ± 0.02* | 0.0175 | 17.1 ± 2.93** | 0.0018 | 3 |
| W287AzF | 7.08 ± 0.01*** | 0.0010 | 10.9 ± 1.5** | 0.0019 | 6 |

Table S5: BigDyn modulation of hASIC1a WT and six variants containing AzF in the acidic pocket. Cells were incubated at SSD-inducing pH for 2 min with or without 3 µM BigDyn before activation at pH 5.6 and the currents were normalized to the average of two preceding control pulses after conditioning at pH 7.6. Values are indicated as mean ± S.D., (n) equals number of cells. (*) denotes significant difference between currents with and without 3 µM BigDyn, p < 0.05; (**): p < 0.01; (***): p < 0.001; ns: not significant; Mann-Whitney test.

| Clone | pH | 3 µM BigDyn | Current after $pH_{SSD}$ 1 (%) | Current after $pH_{SSD}$ 2 (±BigDyn, %) | P value | Current after $pH_{SSD}$ 3 (±BigDyn, %) | Current after $pH_{SSD}$ 4 (%) | P value | n |
|---|---|---|---|---|---|---|---|---|---|
| WT | 6.6 | No | 33.6 ± 25.3 | 15.4 ± 17.1* | 0.0106 | 16.0 ± 17.0 | 14.9 ± 17.2$^{ns}$ | 0.7114 | 24 |
| WT | 6.6 | Yes | 27.0 ± 23.2 | 29.7 ± 25.0$^{ns}$ | 0.9118 | 38.0 ± 34.8 | 20.7 ± 39.3$^{ns}$ | 0.0939 | 10 |
| T236AzF | 6.6 | No | 16.3 ± 18.1 | 7.65 ± 8.63$^{ns}$ | 0.07 | 15.1 ± 19.5 | 7.79 ± 8.33$^{ns}$ | 0.1479 | 19 |
| T236AzF | 6.6 | Yes | 13.7 ± 8.85 | 34.0 ± 20.0** | 0.0032 | 46.0 ± 31.2 | 17.3 ± 15.5$^{**}$ | 0.0039 | 11 |
| K343AzF | 6.7 | No | 18.2 ± 27.3 | 8.75 ± 12.9$^{ns}$ | 0.2480 | 15.9 ± 41.7 | 4.49 ± 6.36$^{ns}$ | 0.1738 | 22 |
| K343AzF | 6.7 | Yes | 54.1 ± 96.5 | 10.5 ± 27.4$^{***}$ | 0.0006 | 42.2 ± 75.9 | 7.61 ± 12.2$^{**}$ | 0.0013 | 18 |
| E344AzF | 6.6 | No | 9.65 ± 12.0 | 2.23 ± 6.39$^{*}$ | 0.0253 | 1.50 ± 11.9 | 0.78 ± 11.1$^{ns}$ | 0.3888 | 19 |
| E344AzF | 6.6 | Yes | 8.29 ± 8.69 | 4.21 ± 5.56$^{*}$ | 0.0326 | 18.0 ± 19.0 | 5.18 ± 7.36$^{**}$ | 0.0077 | 19 |
| D351AzF | 6.8 | No | 23.8 ± 23.7 | 14.1 ± 16.9$^{ns}$ | 0.1371 | 30.5 ± 27.2 | 23.5 ± 30.4$^{ns}$ | 0.2982 | 12 |
| D351AzF | 6.8 | Yes | 14.9 ± 18.2 | 64.1 ± 23.4** | 0.0041 | 80.9 ± 47.0 | 26.8 ± 15.8$^{***}$ | 0.0006 | 7 |
| E355AzF | 7.0 | No | 9.85 ± 5.36 | 9.38 ± 5.73$^{ns}$ | 0.7366 | 19.4 ± 14.1 | 8.87 ± 6.69$^{ns}$ | 0.0516 | 8 |
| E355AzF | 7.0 | Yes | 17.9 ± 11.0 | 67.1 ± 52.2*** | 0.0008 | 125 ± 90.7 | 80.4 ± 99.3* | 0.0241 | 14 |
| K356AzF | 6.6 | No | 9.70 ± 13.5 | 7.94 ± 11.1$^{ns}$ | 0.1465 | 14.8 ± 18.4 | 2.20 ± 1.57*** | 0.0003 | 14 |
| K356AzF | 6.6 | Yes | 23.4 ± 31.5 | 43.2 ± 33.8$^{ns}$ | 0.1145 | 70.2 ± 46.8 | 10.6 ± 12.1*** | 0.0003 | 11 |

Table S6: PcTx1 modulation of hASIC1a WT and five variants containing AzF in the acidic pocket at different pH. Cells were incubated with 100 nM PcTx1 for 2 min before activation at pH 5.6 and the current was normalized to the average of the four preceding and following control pulses after conditioning at pH 7.4. Values are indicated as mean ± S.D., (n) equals number of cells. (*) denotes significant difference between currents at different pH, $p < 0.05$; (**): $p < 0.01$; ns: not significant; one-way ANOVA with Tukey's multiple comparisons test (AzF variants) or Mann-Whitney test (Bpa variants).

| | pH 1 (7.4) | | pH 2 (7.3) | | pH 3 (7.0; 7.2 for WT) | | P value | | |
|---|---|---|---|---|---|---|---|---|---|
| Clone | Normalized response (%) ± S.D. | n | Normalized response (%) ± S.D. | n | Normalized response (%) ± S.D. | n | pH 1 vs pH 2 | pH 2 vs pH 3 | pH 1 vs pH 3 |
| WT | 38.2 ± 31.7 | 8 | 23.7 ± 15.4 | 11 | 2.06 ± 2.50 | 7 | $0.2907^{ns}$ | $0.0907^{ns}$ | 0.0060** |
| T236AzF | 188 ± 82.1 | 5 | 64.0 ± 31.5 | 4 | 46.5 ± 46.4 | 8 | 0.0147* | $0.8708^{ns}$ | 0.0017** |
| E344AzF | 58.0 ± 37.4 | 3 | 23.9 ± 17.2 | 11 | 4.14 ± 4.41 | 6 | 0.0303* | $0.1191^{ns}$ | 0.0020** |
| E355AzF | 66.9 ± 55.5 | 13 | 48.8 ± 63.4 | 8 | 5.03 ± 5.60 | 8 | $0.7051^{ns}$ | $0.2071^{ns}$ | 0.0281* |
| K356AzF | 92.2 ± 40.1 | 12 | 51.7 ± 33.2 | 6 | 43.6 ± 51.6 | 11 | $0.1737^{ns}$ | $0.9298^{ns}$ | 0.0343* |
| D357AzF | 902 ± 926 | 17 | 331 ± 441 | 6 | 260 ± 383 | 9 | $0.6407^{ns}$ | $>0.9999^{ns}$ | $0.3072^{ns}$ |

| D357AzF | pH 4 (6.8) | | pH 5 (6.7) | | pH 6 (6.6) | | pH 7 (6.5) | |
|---|---|---|---|---|---|---|---|---|
| | 1.80 ± 1.78 | 6 | 0.72 ± 1.68 | 5 | 3.58 ± 6.22 | 7 | 2.20 ± 2.16 | 7 |

P values D357AzF

| pH 1 vs pH 4 | pH 1 vs pH 5 | pH 1 vs pH 6 | pH 1 vs pH 7 | pH 2 vs pH 4 | pH 2 vs pH 5 |
|---|---|---|---|---|---|
| 0.0187* | 0.0186* | 0.0191* | 0.0188* | $0.7327^{ns}$ | $0.7298^{ns}$ |
| pH 2 vs pH 6 | pH 2 vs pH 7 | pH 3 vs pH 4 | pH 3 vs pH 5 | pH 3 vs pH 6 | pH 3 vs pH 7 |
| $0.7376^{ns}$ | $0.7338^{ns}$ | $0.6245^{ns}$ | $0.6196^{ns}$ | $0.6326^{ns}$ | $0.6263^{ns}$ |
| pH 4 vs pH 5 | pH 4 vs pH 6 | pH 4 vs pH 7 | pH 5 vs pH 6 | pH 5 vs pH 7 | pH 6 vs pH 7 |
| $0.9943^{ns}$ | $0.9998^{ns}$ | $>0.9999^{ns}$ | $0.9797^{ns}$ | $0.9567^{ns}$ | $>0.9999^{ns}$ |

Bpa variants

| Clone | pH 1 (7.4) | | pH 2 (7.0) | | P value |
|---|---|---|---|---|---|
| | Normalized response (%) ± S.D. | n | Normalized response (%) ± S.D. | n | pH 1 vs pH 2 |
| WT | 40.9 ± 5.25 | 6 | 11.4 ± 9.15 | 6 | 0.0022** |
| T236Bpa | 62.1 ± 47.5 | 6 | 23.0 ± 24.8 | 7 | $0.3660^{ns}$ |
| E344Bpa | 114 ± 25.6 | 6 | 4.00 ± 2.64 | 5 | 0.0043** |
| E355Bpa | 102 ± 41.4 | 5 | 7.89 ± 3.72 | 6 | 0.0043** |
| K356Bpa | 95.4 ± 8.24 | 6 | 15.6 ± 22.1 | 6 | 0.0022** |
| D357Bpa | 187 ± 33.4 | 6 | 6.01 ± 4.92 | 6 | 0.0022** |
